# From variability to consensus: rescoring harmonizes peptide identification across diverse search engines and datasets

**DOI:** 10.64898/2026.03.04.709532

**Authors:** Dirk Winkelhardt, Sven Berres, Julian Uszkoreit

## Abstract

Peptide-spectrum match (PSM) rescoring has become standard in proteomics workflows, improving peptide identification accuracy across diverse search engines. Despite the availability of multiple rescoring strategies, systematic comparisons spanning several search engines, datasets, and database configurations remain limited. Here, we benchmarked seven publicly available search engines, evaluating standard target-decoy-based false discovery rate (FDR) estimation alongside Percolator, MS2Rescore, and Oktoberfest across four datasets acquired on different mass spectrometry platforms in data-dependent mode and searched against protein databases of varying size and composition.

Rescoring substantially increased identification consensus and reduced variability between search engines, with prediction-based approaches yielding the largest gains. While database size had limited impact for human datasets, it significantly affected identification rates on a metaproteomic dataset. Entrapment-based evaluation indicated generally adequate FDR control across methods, although prediction-based rescoring exhibited a higher tendency toward FDR underestimation in specific configurations.

Overall, advanced rescoring strategies harmonize peptide identification outcomes across search engines, thereby enhancing robustness and comparability in proteomics analyses. However, careful feature selection and appropriate database choice remain essential to ensure reliable FDR control and optimal performance across diverse experimental settings.

## Introduction

Rescoring of peptide-spectrum matches (PSMs) has become standard practice in data-dependent acquisition (DDA) proteomics data analysis in recent years. Established methods such as Percolator^1^ and PeptideProphet^2^ have been integrated into proteomics workflows for more than two decades. These approaches aim to distinguish correct from incorrect PSM identifications. The original PeptideProphet used linear models based mainly on search engine scores and a limited set of additional features. In contrast, Percolator introduced a semi-supervised learning strategy that leverages the target–decoy approach (TDA)^3^, where decoy sequences are used to represent the distribution of incorrect or randomly matched peptides, thereby enabling more accurate estimation of identification confidence.

More recent methods incorporate additional features derived from predicted peptide fragment ion spectra, including MS2Rescore^4^ and Oktoberfest^5^. In these approaches, fragment ion intensities are predicted using dedicated algorithms^6,7^ and compared to experimentally measured spectra for each PSM. Similarity metrics between predicted and observed spectra are then used as additional features for rescoring, typically within Percolator or alternative implementations such as Mokapot^8^, to improve discrimination between correct and incorrect identifications.

In parallel with developments in rescoring methodologies, peptide-spectrum matching algorithms, commonly referred to as peptide search engines, have also evolved substantially. Numerous search engines are now available and are integrated into diverse workflow environments. The selection of a particular algorithm often depends on user preference, computational infrastructure, and project-specific requirements.

The performance gains respective to number of identified PSMs achieved by individual rescoring methods have been demonstrated in their respective original publications. Furthermore, their application to emerging proteomics fields characterized by large search spaces, such as immunopeptidomics and proteogenomics, has recently been reported^9^. A recent comparative study^10^ evaluated MS2Rescore, Oktoberfest, and inSPIRE^11^. However, these comparisons were limited to a single peptide spectrum matching algorithm and typically a single protein database.

Here, we present a comprehensive comparison across seven publicly available DDA search engines: X!Tandem^12^, Andromeda (MaxQuant)^13^, Comet^14^, MS Amanda^15^, MS-GF+^16^, MSFragger^17^, and Sage^18^. For each engine, we evaluated standard target-decoy-based scoring as well as rescoring using Percolator alone, MS2Rescore, and Oktoberfest. These three rescoring strategies represent the most widely used approaches currently available. Benchmarking was performed on four datasets acquired on different mass spectrometry platforms and searched against protein databases of varying sizes. The analysis focused exclusively on tandem MS (MS2)-based DDA peptide identification; no quantitative analyses or analyses of data-independent acquisition (DIA) samples were performed.

Identification results were compared at both the PSM and peptidoform levels across search engines, databases, and rescoring strategies. While identification rates based solely on raw search score, derived false discovery rate (FDR) estimation varied substantially between search engines, rescoring markedly increased consensus and reduced performance differences across methods.

## Materials and methods

### Pipeline

To investigate the performance of the peptide identification tools, a custom Nextflow^19^ pipeline called *mspepid* was built. As Thermo Fisher Scientific and Bruker are among the most commonly used vendors in proteomics, the pipeline was designed to accept data from both platforms. Both formats were converted into the open standard mzML format using ProteoWizard^20^ and tdf2mzml (https://github.com/mafreitas/tdf2mzml), respectively, to simplify data handling and homogenize access.

Protein databases are required to adhere to the UniProt FASTA header scheme. If a given database contains only target sequences, decoys are generated using the DecoyDatabase tool of OpenMS^21^ reversing the peptide sequences to support target-decoy based false discovery estimation^3,22^. In addition to conventional target–decoy databases, mspepid can also generate entrapment databases to assess the false discovery control by leveraging FDRBench (commit 146f77)^23^. In the described analysis, threefold shuffled entrapment databases were generated.

The given MS2 spectra are identified using the seven publicly available search engines: Comet, MaxQuant, MS Amanda, MSFragger, MS-GF+, Sage, and X!Tandem. As some search engines report the resulting peptide–spectrum matches (PSMs) in non-standardized formats, psm-utils^24^ is used to collect the PSMs into its lossless TSV format. Based on this, false discovery rate (FDR) estimation is performed using the raw PSM scores of the search engines and custom implementation in a Python notebook allowing the selection of the search engine score for the calculation (see GitHub link at end of the section). To have scores following a “higher score better” metric, the following scores were selected for the FDR estimation of the raw PSM scoring (referred to as score in the figures).

- Comet: *−ln*(*E-value*)
- MaxQuant: *MaxQuant Score*
- MS Amanda: *Amanda score*
- MS Fragger: *−ln*(*expect*)
- MS-GF+: *−ln*(*spec E-value*)
- Sage: *Discriminant Score*
- X!Tandem: *−ln*(*expect*)

After export of the search results into Percolator PIN files provided by psm-utils, rescoring is performed by three popular tools: Percolator^1^, MS2Rescore^4^ (respectively TIMS2Rescore for timsTOF data^25^), and Oktoberfest^5^.

To ensure consistent processing of the provided data, *mspepid* aligns shared search engine parameters, such as mass tolerances, as closely as possible across tools. Search engine-specific parameters are adjusted depending on the input data to ensure optimal performance of each algorithm (compare settings given in databases). Upon conversion of the search results into the psm-utils TSV format, only the PSMs with the best score per spectrum were retained (on a tie, all best scoring PSMs per spectrum) and used for the analyses in this study.

Although the pipeline was carefully designed to handle data consistently across all search engine and rescoring tool combinations, certain exceptions were necessary due to runtime or execution constraints. For example, when analyzing Bruker TIMS-TOF data with MaxQuant, the Bruker .d-directory is not converted to mzML because this conversion causes issues that affect subsequent rescoring: The applied version of MS2Rescore could not match the spectra to PSMs in this case, as the PSM indices were wrong. However, using the vendor file format did not affect the rescoring. To reduce the runtime required by MS-GF+ when searching very large protein databases (such as the ProHap^26^ database and the DB1 of CAMPI), the pipeline splits the databases prior to the search into 10 (ProHap) respectively 20 (DB1) chunks of equal amounts of protein entries. The resulting mzIdentML files are then merged afterward using MzidMerger (1.4.26, https://github.com/PNNL-Comp-Mass-Spec/MzidMerger). MzidMerger did also take care to adjust the spectrum-E-value and E-value of MS-GF+ for each PSM, as this value is database dependent. This approach is similar to the procedures implemented in MSFragger and Sage, which perform database splitting and merging automatically with the respective parameters. Additionally, to further improve computation time for MS-GF+, mzML files were split to contain a maximum of 10,000 spectra, and the results were merged accordingly.

MS2Rescore and Oktoberfest were used to create their additional features and add them as Percolator input files, while the actual rescoring was performed by externally running Percolator. This allowed for a more modular approach and returns identical results as the integrated Percolator routines in each tool. A fair comparison of post-processing steps was ensured, by removing the linear discriminant analysis (LDA) score calculated by Oktoberfest before passing the generated features to Percolator. The LDA score represents Oktoberfest’s attempt to separate targets from decoys and would otherwise provide a near-perfect separation label to Percolator.

The pipeline was implemented to be as reproducible as possible by using containerized versions of each software component (Supplementary Table 1). If no containerized version was available, an appropriate image was created and made publicly accessible via GitHub and Quay.io. The only exception is MSFragger, whose license prevents redistribution of the executable. Consequently, the container image must be built locally after downloading the binary manually and accepting the license. The build instructions are provided in the *mspepid* repository Makefile.

Final data analyses and visualizations were generated using a Python notebook (available at https://github.com/medbioinf/mspepid-analysis). For several analyses, peptidoforms were considered instead of individual PSMs. A peptidoform was defined by aggregating all PSMs sharing an identical amino acid sequence and identical modifications into a single entity. During this aggregation step, leucine and isoleucine were treated as indistinguishable. The FDR at the peptidoform level was calculated after aggregation, and results were filtered accordingly.

### Datasets and databases

In this study, *mspepid* was used to reanalyze a total of four DDA datasets. All datasets were downloaded from the public repositories MassIVE^27^ and PRIDE^28^. The datasets were selected to represent a broad range of human-based studies, including tissue samples, cell cultures, and metaproteomics analyses. Two datasets originate from human subsets measured by an Orbitrap and a TIMS-TOF of the label-free quantification benchmark dataset published by Van Puyvelde et al.^29^ (PRIDE ID PXD028735). In this datasets, the identical sample was measured on two different machine setups in technical triplicates. The third analysed dataset is a subset of the Cancer Array dataset from Tüshaus et al.^30^ (MassIVE ID MSV000095036), representing a typical clinical measurement of tissue samples on one single machine setup. For these human datasets three protein databases of different sizes were used: the human Swiss-Prot and the human reference proteome, both containing isoforms, provided by UniProt^31^, as well as the ProHap database^26^, which contains numerous annotated protein sequence variants. Before spectrum identification, the UniProt databases were enriched with contaminants from the database compiled by Frankenfield^32^, as summarized in Table 1. ProHap provides already a database with common contaminants.

In addition to the human datasets, we analyzed run S07 from the metaproteomic benchmarking study *Critical Assessment of MetaProteome Investigation* (CAMPI) by Van Den Bossche^33^ (PRIDE ID PXD023217). This dataset was searched against the two protein databases *GUT_DB1IGC* (DB1) and *GUT_DB2MG* (DB2) provided by the CAMPI authors. Unlike the human samples investigated, the CAMPI dataset comprises metaproteomics experiments and therefore requires these specially constructed protein database integrating sequences from multiple species. DB1 consists of proteins translated from the “integrated gene catalog (IGC)”, which comprises genes of the microbiome of independently measured gut samples^34^. DB2 was created using genomics and transcriptomics sequencing data of the same sample which also underwent the proteomics measurement (for more details refer to the original publication^33^). As the provided databases contained asterisks in the amino acid sequences, these needed to be removed for compatibility with all search engines. Inside the sequence, the asterisks were exchanged with the amino acid X (placeholder for any amino acid in the applied search engines), at the protein C-terminus they were simply removed.

An overview of the datasets, the reanalyzed files, and the nomenclature used throughout this manuscript is provided in Table 2. For the entrapment analyses, all datasets were reanalyzed once more against one of the respective databases: for the Cancer Array, Orbitrap, and timsTOF datasets, the human reference proteome was used; for the CAMPI dataset, DB1 was used.

**Table 1:** Protein database names, counts (including contaminants), sources and versions.

| name | nr entries | source | date/version |
| --- | --- | --- | --- |
| Human SwissProt | 42,900 | https://www.uniprot.org/ | 2025_01 |
| Human Reference Proteome | 105,867 | https://www.uniprot.org/ | 2025_01 |
| ProHap-all | 349,894 | https://zenodo.org/records/12671237 | 240527_ProHap_ALL |
| GUT_DB1IGC (DB1) | 9,879,896 | PRIDE archive (PXD023217) | 18/12/2020 (upload) |
| GUT_DB2MG (DB2) | 441,558 | PRIDE archive (PXD023217) | 18/12/2020 (upload) |

**Table 2:** The reanalyzed datasets in this study.

| dataset | repository and accession | reanalyzed MS runs (MS2 spectra per run) | instrument, tolerances and databases |
| --- | --- | --- | --- |
| Cancer Array | MassIVE, MSV000095036 | P065440_15SPD_FMS-30-40-50-60-70_R2 (102,963),<br>P065440_15SPD_FMS-35-45-55-65-75_R2 (102,803),<br>P065441_15SPD_FMS-30-40-50-60-70_R2 (103,280),<br>P065441_15SPD_FMS-35-45-55-65-75_R2 (102,568),<br>P065442_15SPD_FMS-30-40-50-60-70_R2 (103,418),<br>P065442_15SPD_FMS-35-45-55-65-75_R2 (102,987),<br>P065443_15SPD_FMS-30-40-50-60-70_R1 (100,719),<br>P065443_15SPD_FMS-35-45-55-65-75_R1 (101,013),<br>P065444_15SPD_FMS-30-40-50-60-70_R1 (102,020),<br>P065444_15SPD_FMS-35-45-55-65-75_R1 (102,320) | Orbitrap Exploris 480<br>precursor: 40 ppm,<br>fragment: 0.05 Da<br>Human Swiss-Prot,<br>Human Reference Proteome,<br>ProHap-all |
| CAMPI | PRIDE, PXD023217 | S07 (46,847) | Q Exactive HF<br>precursor: 10 ppm<br>fragment: 0.02 Da<br>DB1 and DB2 |
| Orbitrap | PRIDE, PXD028735 | LFQ_Orbitrap_DDA_Human_01 (118,387),<br>LFQ_Orbitrap_DDA_Human_02 (115,801),<br>LFQ_Orbitrap_DDA_Human_03 (114,340) | Q Exactive HF-X<br>precursor: 20 ppm<br>fragment: 0.02 Da<br>Human Swiss-Prot,<br>Human Reference Proteome,<br>ProHap-all |
| timsTOF | PRIDE, PXD028735 | LFQ_timsTOFPro_PASEF_Human_01 (379,781),<br>LFQ_timsTOFPro_PASEF_Human_02 (378,725),<br>LFQ_timsTOFPro_PASEF_Human_03 (379,776) | timsTOF Pro<br>precursor: 20 ppm<br>fragment: 0.05 Da<br>Human Swiss-Prot,<br>Human Reference Proteome,<br>ProHap-all |

For all datasets, carbamidomethylation of cysteine (C) was used as the only static modification, and oxidation of methionine (M) as only variable modification. Trypsin was used as the cleavage agent, and an automatic target-decoy search was deactivated (where applicable). The precursor and fragment tolerances were applied according to the datasets as stated in Table 2. Besides certain choices for output files and the parallel processing of split databases for the ProHap database and DB1 (Sage and MS Fragger), all other settings were left to the respective tool’s default values.

The results of the reanalyses together with their respective mass spectrometry data have been deposited to the ProteomeXchange Consortium (http://proteomecentral.proteomexchange.org) via the PRIDE partner repository^28^ with the following dataset identifiers:

- PXD075141 and DOI 10.6019/PXD075141 for the CAMPI dataset,
- PXD075199 and DOI 10.6019/PXD075199 for the Cancer Array dataset,
- PXD075209 and DOI 10.6019/PXD075209 for the Orbitrap dataset, and
- PXD075206 and DOI 10.6019/PXD075206 for the timsTOF dataset.

## Results

After the initial PSM identification for each combination of dataset, protein database, and search engine, we generated comparative plots to visualize the results. In the following sections, we describe these plots and highlight the key findings from each analysis. While few plots are shown in the main manuscript, plots for all combinations are shown in the supplemental material.

### Target-decoy distribution plots

To assess whether the TDA assumptions appear to be violated, we generated target–decoy distribution plots following the methodology described in^35^. The assumption under scrutiny here is that low scoring incorrect PSMs and decoys follow a similar distribution to be able to estimate the FDR and cut the ranked identification on a given 1% FDR q-value position^3^. For most combinations of dataset, database, and search engine, these plots exhibited distributions consistent with the expected behavior, as illustrated in Figure 1. Separation between putatively correct and incorrect identifications was least pronounced when relying solely on raw search scoring and improved with Percolator and prediction-based rescoring methods. For Sage, however, little difference was observed between the search engine’s scoring and Percolator rescoring, likely because Sage already applies an internal linear discriminant model to compute its primary score.

**Figure 1:**
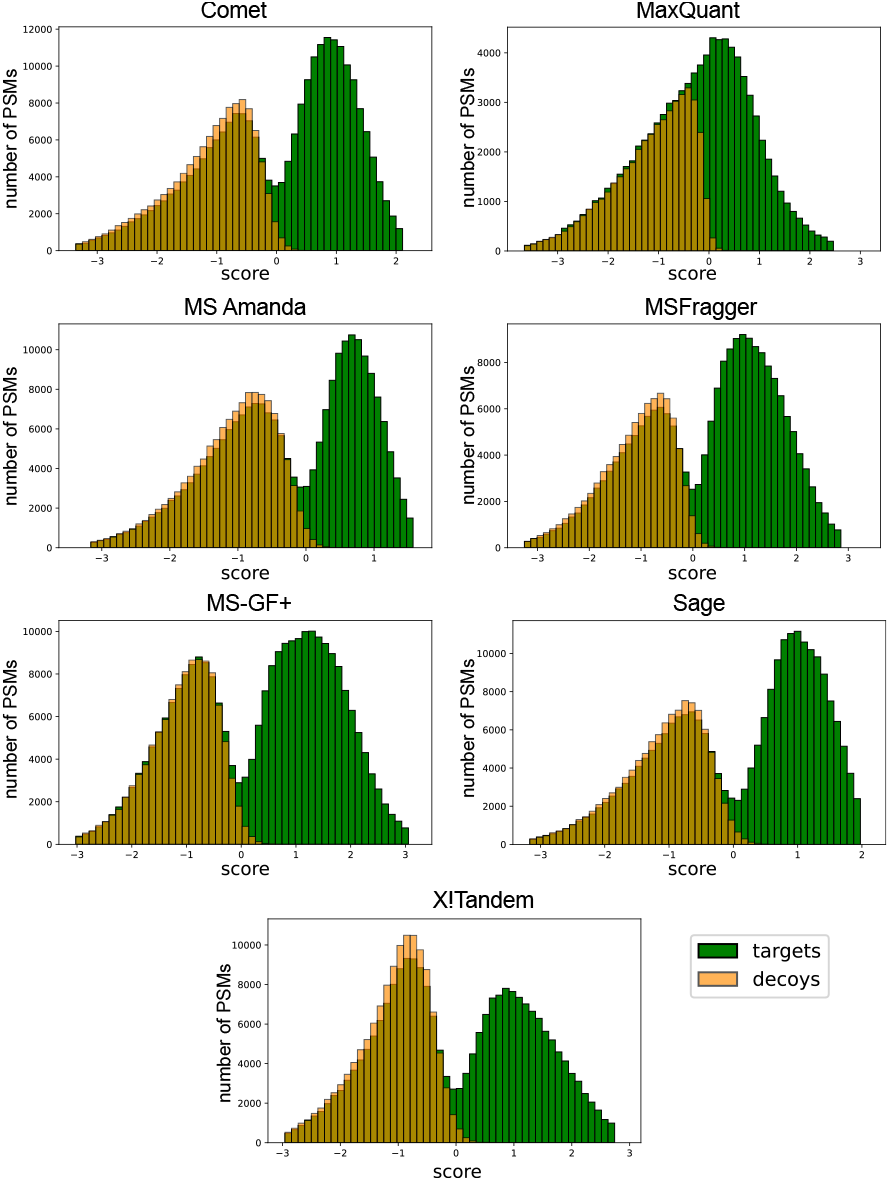
Target–decoy distribution of peptide–spectrum matches (PSMs). Shown is the first MS run of the timsTOF dataset searched against the human proteome database using MS2Rescore. Each search engine exhibits slightly different distributions, but in general, the assumption that low scoring incorrect target PSMs and decoys follow a similar distribution is supported. Although for this dataset, MaxQuant showed less pronounced separation into two distributions, all methods effectively separate correct from incorrect identifications.

Some combinations, however, deviated from this general pattern. For the timsTOF dataset, ProHap database, and MS-GF+, Percolator without spectrum prediction-based features slightly violated the assumption that decoys and incorrect target PSMs share the same distribution, introducing minor bias in separation. More strikingly, rescoring with MS2Rescore or Oktoberfest produced nearly perfect separation between target and decoy PSMs, clearly violating the assumptions underlying FDR estimation (see Supplemental Figure S21 for the details). For the analyses described in this work, the slightly violating Percolator rescoring data were retained as they were instead of performing a fix like the following, which would be advised if this data should be published on its own (Supplemental Figure S20b). The issue with MS2Rescore and Oktoberfest was resolved by removing a single feature before rescoring - the negative logarithm of the MS-GF+ E-value - and the adjusted data were used in all subsequent analyses. The MS-GF+ E-value has the special characteristic, that it actually is not calculated for one spectrum (like the MS-GF+ spec-E-value), but over all PSMs, similar to the also excluded lda-score of Oktoberfest. This might also be the reason for it to perform as a score to separate the target and decoy PSMs on rescoring in a similar way, at least in the mentioned datasets.

### Identified PSMs as a function of q-value thresholds

To provide an initial overview of identification performance, we generated “pseudo-ROC” plots in which the number of identified target PSMs is plotted against the corresponding q-value threshold. Figure 2 shows these plots for the Orbitrap dataset. These plots enable comparison of each combination of search engine and rescoring strategy across datasets and databases.

**Figure 2:**
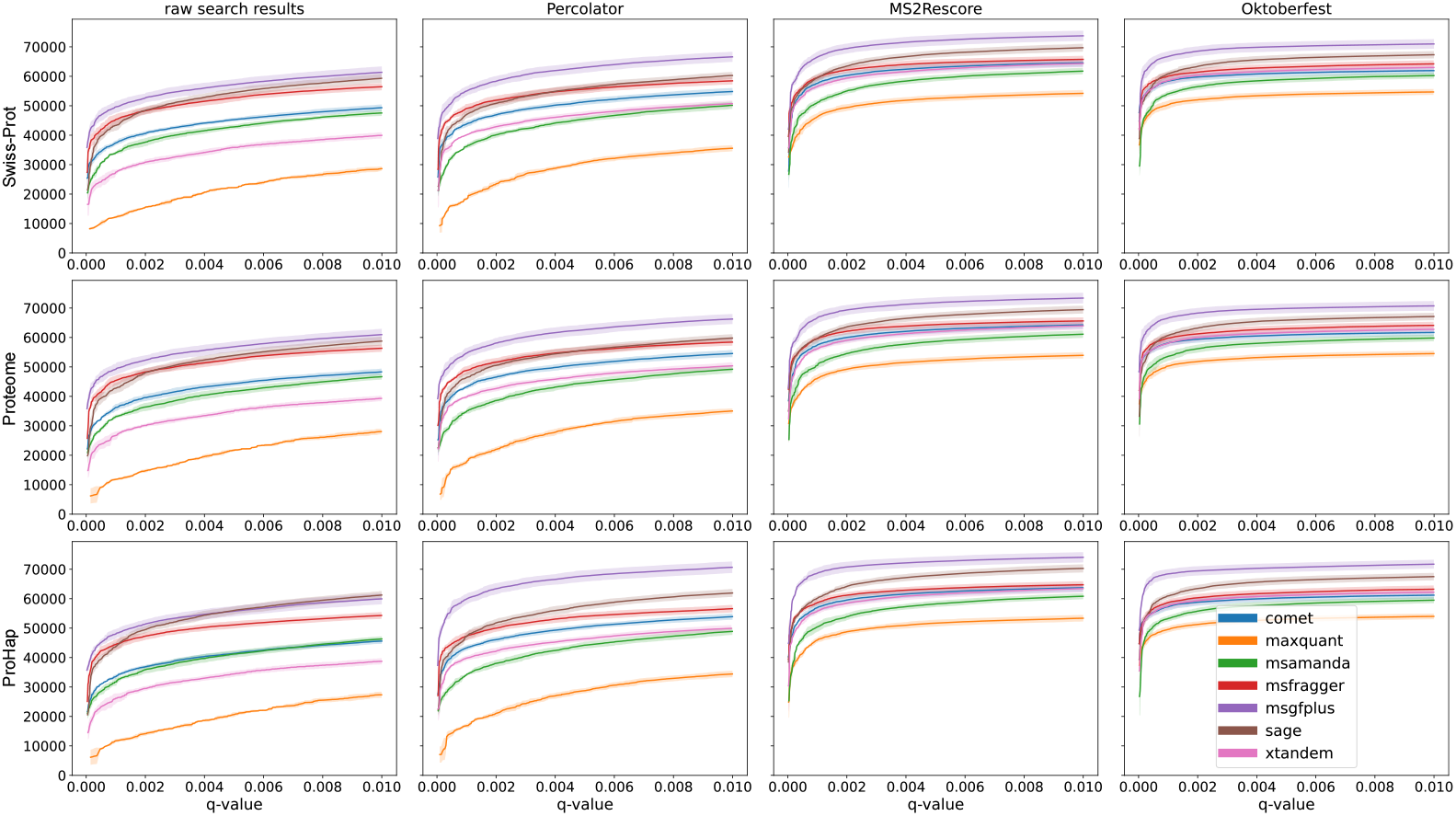
Number of identified target PSMs as a function of the q-value threshold. Depicted are the mean values (dark hue) as well as the maximum and minimum (lighter shades) observed across all runs. The plots shown correspond to the Orbitrap dataset and illustrate general trends across search engines. In particular, MS-GF+ and MaxQuant consistently exhibit the highest and lowest identification rates, respectively, across most conditions. Additionally, advanced rescoring methods not only increase the total number of identifications but also reduce differences between search engines, resulting in more comparable identification counts.

Some trends emerge already from these analyses. When relying solely on the raw PSM scoring for FDR estimation, the number of PSMs below the 1% FDR threshold differs substantially between search engines. Although these differences persist to some extent after applying Percolator, they are markedly reduced following the prediction-based rescoring approaches, which generally increase the total number of identifications while decreasing variability between search engines.

When considering the overall number of identified PSMs per search engine, the relative ranking remains largely consistent, with MS-GF+ yielding the highest numbers and MaxQuant the lowest across most conditions. Some notable deviations were observed, however. For the timsTOF dataset, X!Tandem without rescoring did not produce any PSMs fulfilling the 1% FDR threshold due to high-ranking decoy hits. In the CAMPI dataset, MS Amanda yielded the fewest identifications in most combinations, although the absolute differences between search engines were smaller than in other datasets (Supplemental Figure S23).

### Number of identifications per search engine and rescoring method

To provide an intuitive comparison of rescoring effects for each dataset–database combination, we generated bar charts summarizing the total number of identified peptidoforms per search engine and rescoring strategy (Figure 3). The reported values represent the mean number of identified peptidoforms across runs within each dataset, with error bars indicating the standard deviation.

**Figure 3:**
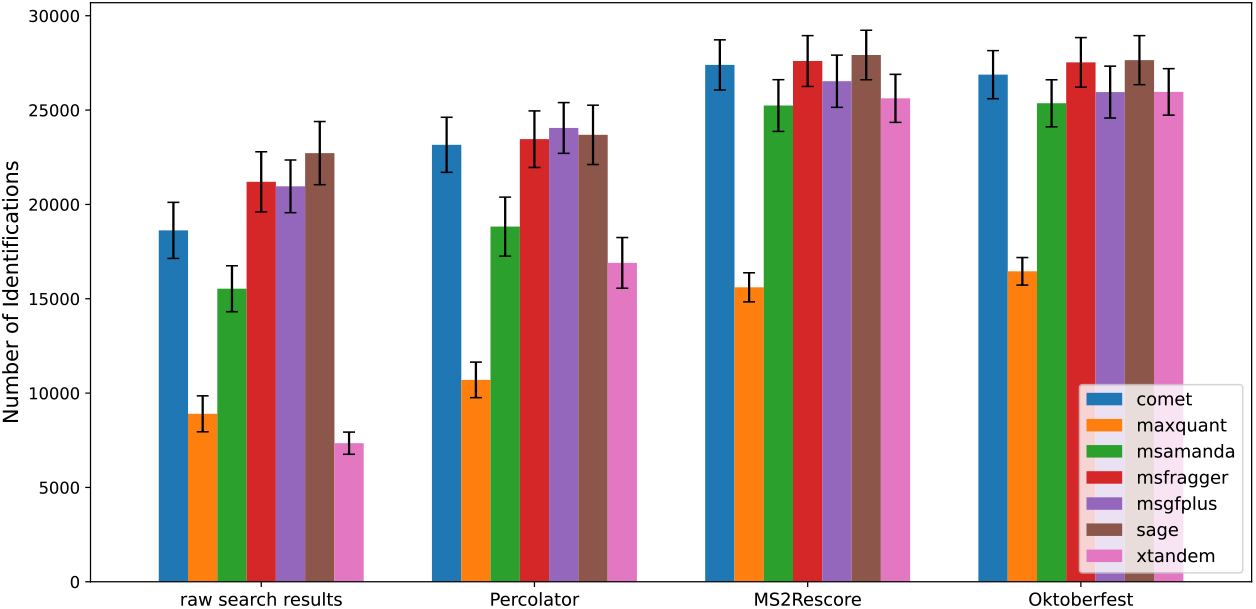
Number of identified peptidoforms for the Cancer Array dataset and Proteome database. The bar charts show the mean number of identified 1% FDR-valid peptidoforms across all runs of the dataset, with error bars indicating the standard deviation. When using the raw search results only or Percolator-based rescoring for FDR estimation, substantial differences between search engines are observed. Prediction-based rescoring methods reduce these discrepancies drastically, resulting in reduced variability in identification numbers across most search engines. This plot furthermore highlights the ability to totally “rescue” the identifications of the very low performing X!Tandem by rescoring.

Overall, these plots recapitulate the trends observed at the PSM level (e.g. for the peptidoform data in Figure 3 compare Supplemental Figure S22 showing this trend for the same dataset), demonstrating that the effects of rescoring propagate consistently to the peptidoform level. Across all datasets, advanced rescoring approaches not only increased the total number of identifications but also reduced the difference in numbers of identifications between search engines, resulting in more comparable performance.

A quantitative inspection of the Cancer Array dataset searched against the proteome database illustrates this effect (Figure 3). When applying raw PSM scoring for FDR estimation only, the largest difference between the search engines’ average number of results per run was 15,373 peptidoforms. This difference decreased to 13,351 with Percolator, or to 7,150 when excluding MaxQuant. After rescoring with MS2Rescore and Oktoberfest, the differences were further reduced to 12,313 and 11,191, respectively, or to 2,679 and 2,282 when MaxQuant was excluded. Expressed as a percentage of the highest number of identifications per search engine, this corresponds to a reduction from 67% (raw search results) to 9.6% (MS2Rescore) and 8.2% (Oktoberfest).

For the Orbitrap dataset (proteome database) (Supplemental Figure S32), the relative difference decreased from 48% when applying only the search engine-based scoring to 20% and 16% with MS2Rescore and Oktoberfest, respectively. When excluding MaxQuant, the reduction was from 32% to 12% and 13%.

In the timsTOF dataset (Supplemental Figure S35 for proteome database, but similar numbers on the other two databases), X!Tandem was substantially outperformed by the other search engines when applying FDR estimation with raw search results only, resulting in a maximal difference of 95%. Rescoring reduced this discrepancy to 39% (MS2Rescore) and 44% (Oktoberfest), and even to 12% and 11%, respectively, when excluding MaxQuant.

For the CAMPI dataset, MaxQuant yielded more identifications than MS Amanda after MS2Rescore and Oktoberfest rescoring (Supplemental Figure S29). The overall difference for search engines using DB1 decreased from 61% applying raw search scores to 24% (MS2Rescore) and 19% (Oktoberfest), or further to 14% and 11%, respectively, when excluding MS Amanda.

For the human datasets, database size had minimal influence on the number of identified peptidoforms, with only minor differences observed between databases of varying sizes (Supplemental Figures S26 - S28 and S31 - S36). In contrast, the CAMPI dataset exhibited more pronounced differences between DB1 and DB2 (Supplemental Figures S29 and S30, be aware that the y-axes are scaled differently), with the substantially larger DB1 consistently yielding a higher number of identifications. After rescoring with MS2Rescore and Oktoberfest, the numbers of identifications were 22% - 31% percent higher in DB1 in comparison to DB2.

### Overlap of databases per search engine

To further assess the influence of database selection, we generated UpSet plots for each search engine to visualize overlaps between protein databases and FDR estimation strategies at the peptidoform level (see supplemental data section 4). With these plots, it is possible to inspect the influence of the databases and the rescoring methods on the resulting number of identifications and especially the overlap of identifications. For most combinations of dataset and search engine, rescoring had a greater impact on .the number of additional overlapping identifications than database choice. The only exception was the CAMPI dataset, for which DB1 consistently yielded substantially more identified peptidoforms than DB2 and also the largest gain of overlapping identifications was added by searching against DB1 with any FDR control method, instead of applying rescoring methods.

This observation is consistent with the trends shown in the corresponding bar plots described before. Notably, the effect of yielding more overlapped identifications by rescoring was more pronounced for datasets acquired on Orbitrap instruments compared to the timsTOF dataset (compare Supplemental Figures S39 and S40).

### Overlap of all search engines per rescoring method

Finally, UpSet plots were generated for each combination of database and rescoring strategy to visualize differences between search engines under the respective conditions. For these analyses, peptidoforms, including modifications, were compared, while distinctions between leucine and isoleucine were disregarded. The plots report the average number of peptidoforms per MS run for each intersection group. The plots for all combinations can be found in Supplemental Figures S41 - S51, while Figure 4 only highlights the numbers in the intersection of all search engines per rescoring respectively raw PSM scoring, database and dataset. The overview Figure 4 gives a quick representation of the fact, that the number of identifications in the intersection of all search engines and hence the consensus is always increased by rescoring in comparison to the raw PSM scoring, and also more by the prediction-based methods than by Percolator alone.

**Figure 4:**
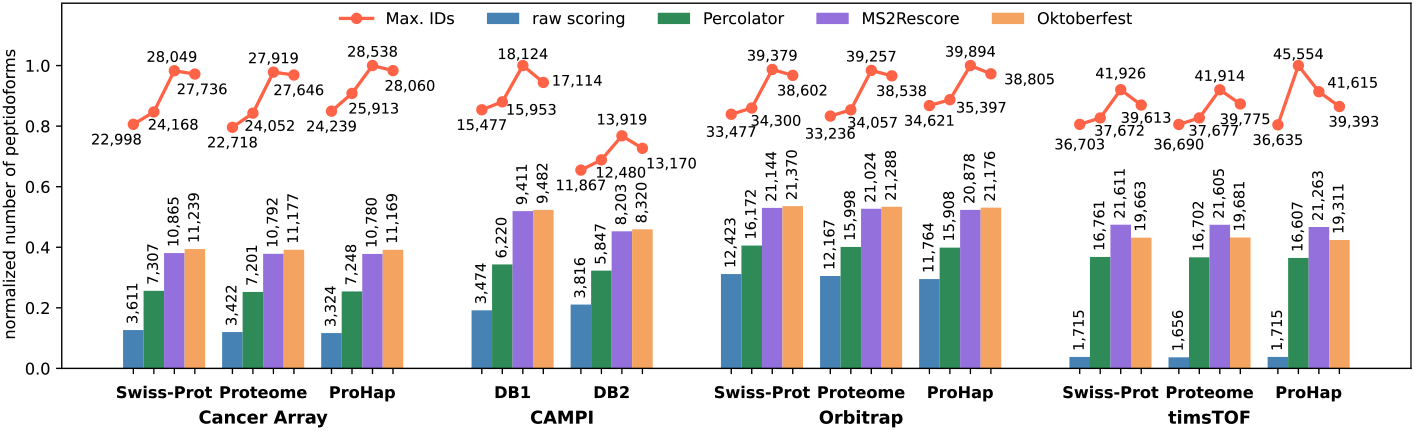
Overlap of peptidoforms identified by all search engines for each combination of dataset and database. The plot shows as bar charts the number of peptidoform identifications per rescoring method, respective the usage of raw identifications, for each combination of database and dataset. Additionally, the highest number of identified peptidoforms for the single best search engine is given (as the line plot “Max. IDs”). The heights of the plot are normalized per dataset to the highest “Max. IDs”. The single numbers are taken from Supplemental figures S41 - S51. The plot highlights the general trend, that Percolator already increases the overlap and consensus, while this is even improved by the prediction-based methods increase the total overlap vastly. Especially for the timsTOF dataset the increase from the total overlap is almost tenfold, though this is due to the fact that X!Tandem had very low identifications with raw PSM scoring

Several observations can be made by inspecting these supplemental plots. The Cancer Array and CAMPI datasets exhibit more dispersed overlap patterns, characterized by a larger number of intersection groups exceeding the reporting threshold. In contrast, the Orbitrap and timsTOF datasets show higher consensus among search engines, reflected by fewer distinct intersection groups. The timsTOF dataset demonstrates the highest overall consensus despite having the largest number of PSMs per run. The main exception is MaxQuant, which almost consistently reports fewer identifications across runs, whereas MS-GF+ typically yields a subset of additional peptidoforms not identified by other search engines.

In the timsTOF dataset, X!Tandem’s scoring function fails to differentiate between good and bad (respectively decoy) identifications, which is shown by the number of identifications when controlling the FDR relying solely on the raw PSM scores. In this case, high-scoring decoys raise the 1% FDR threshold, resulting in a markedly reduced number of identified peptidoforms, which is well reflected in the overview Figure 4. All rescoring methods substantially mitigate this effect, yielding identification numbers comparable to those of other search engines and increasing the total overlap.

The CAMPI dataset (Supplemental Figures S44 and S45)shows a distinct pattern. Without rescoring (raw search results), all search engines except MaxQuant show high consensus. After rescoring, however, MS Amanda emerges as a more pronounced outlier. Nevertheless, the consensus among the remaining search engines, i.e. all except MaxQuant and MS Amanda, remains high.

### Assessment of FDR estimation using entrapment

To evaluate the FDR control across methods, the FDRbench framework described in^23^ was applied to one representative database per dataset and to all FDR estimation strategies. The introduction of entrapment protein sequences in the original database allows the estimation of the actual proportion of false positives as the “false discovery proportion” (FDP) besides the estimation of the FDR using the target-decoy approach. Ideally, the FDR and FDP should be identically for the positions in a ranked list of identifications. The resulting upper and lower bounds of the FDP of peptidoforms are shown in Supplemental Figures S52–S55.

For most assessed methods, the absolute lower bound of the FDP lies below the FDP–FDR diagonal, satisfying this key requirement for a valid FDR control. Some of the benchmarked combinations though fail this requirement, besides others MaxQuant slightly for all FDR estimations in the Cancer Array and the CAMPI dataset, but also Comet and X!Tandem with MS2Rescore and Oktoberfest in the CAMPI dataset and MS Fragger with Oktoberfest on all combinations except the timsTOF dataset.

The upper bound is frequently located below or close to the diagonal, although also here notable deviations are observed. MS Amanda represents the most prominent case, consistently remaining below the control limit across methods, indicating stable, maybe occasionally even conservative, FDR control. The only exception is the CAMPI dataset, where the entrapment methods assumes a much higher FDP than any estimated FDR for MS Amanda. This is due to the fact, that the scoring function of MS Amanda ranks many entrapment peptides directly at the top position. To better visualize the offset, a larger FDP and FDR range is plotted in Supplemental Figure S56a.

Interestingly, MS-GF+, which in all other datasets performs rather conservative, underestimates the FDR with the upper bound in the timsTOF dataset (Supplemental Figure S55e). This effect might be due to the fact, that the applied version of MS-GF+ does not have a model for timsTOF data, but the common HCD model was used, which is well suited for all other machines.

Overall, for most remaining combinations, FDR control appears adequate, although in some cases the observed FDP slightly exceeds the nominal FDR. In particular, for MSFragger in the Cancer Array, CAMPI, and Orbitrap (compare Figure 5) datasets, Oktoberfest shows an initial increase in FDP at low FDR thresholds before approaching the diagonal at higher FDR values. In Figure S56b it can be seen, that also this offset returns below the FDP diagonal, but at higher FDR q-values. Overall, the prediction-based rescoring methods MS2Rescore and Oktoberfest exhibit a greater tendency to underestimate the true FDR compared to raw PSM scoring and Percolator.

**Figure 5:**
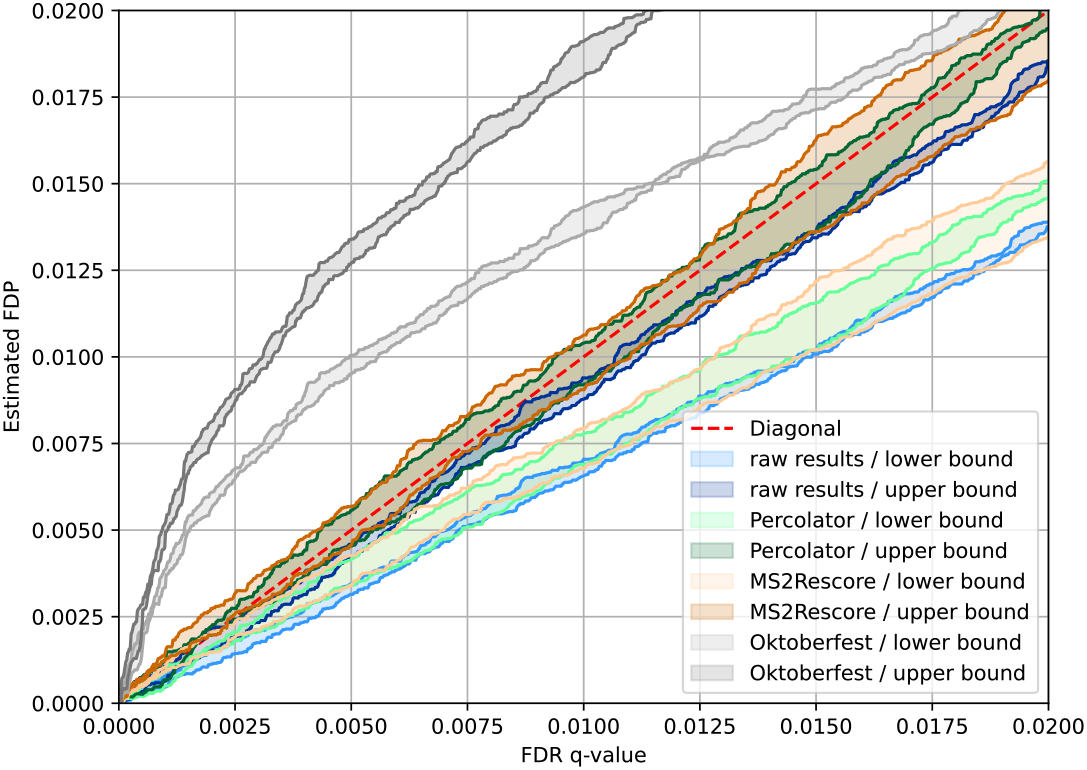
Entrapment-based assessment of FDR estimation for the Orbitrap dataset using MSFragger. Using only raw PSM scoring, both the lower and upper FDP bounds lie below the FDP–FDR diagonal, with the upper bound indicating conservative behavior. Rescoring with Percolator and MS2Rescore shifts the lower bound closer to the diagonal, while the upper bound partially exceeds it, suggesting slight FDR underestimation. Rescoring with Oktoberfest exhibits a distinct pattern, with pronounced FDR underestimation at low FDR thresholds that diminishes at higher thresholds.

### Resources and execution times

While the focus of this study was the comparison of search engine results and the impact of rescoring strategies, an important practical difference between search engines lies in their computational resource consumption and execution time. To illustrate these differences for a representative dataset–database combination, execution times as well as CPU and memory usage are summarized in Table 3.

**Table 3:** Resource usage and execution time for individual pipeline steps during reanalysis of the Orbitrap dataset using the proteome database. For each step, the table reports the average runtime (wall time in seconds), average CPU utilization (in %, where 100% corresponds to full usage of a single CPU core; multithreaded processes may exceed 100%), and average peak memory consumption (in GB) of the specified module in the pipeline. The number of instances required for each respective module for the whole re-analysis is provided in the column “nr. instances”. As the dataset comprised three MS runs, most search engines had to be executed three times. Exceptions were Sage and MS-GF+. Sage was executed once and its results were subsequently split for downstream processing. For MS-GF+, each raw file was divided into 12 parts, each containing up to 10,000 MS2 spectra, prior to analysis to improve computational performance. Percolator and the additional feature generation steps by MS2Rescore and Oktoberfest did not differ for the various search engines, hence they are grouped into single lines and each was executed 21 times for the three runs and seven search engines.

| Pipeline step | duration (sec) | CPU (%) | peak memory (GB) | nr. instances |
| --- | --- | --- | --- | --- |
| Comet | 237 | 1520 | 5.8 | 3 |
| MaxQuant | 4054 | 150 | 745.7 | 3 |
| MS Amanda | 240 | 791 | 251.6 | 3 |
| MSFragger | 276 | 726 | 25.5 | 3 |
| MS-GF+ | 2903 | 482 | 18.1 | 36 |
| Sage | 438 | 1167 | 10.5 | 1 |
| X!Tandem | 257 | 646 | 8.9 | 3 |
| MS2Rescore | 485 | 259 | 100.5 | 21 |
| Oktoberfest | 324 | 110 | 10.7 | 21 |
| Percolator (raw results) | 10 | 336 | 0.5 | 21 |
| Percolator (MS2Rescore) | 195 | 334 | 0.6 | 21 |
| Percolator (Oktoberfest) | 87 | 330 | 0.6 | 21 |

The table reveals substantial variation in runtime between search engines, ranging from 146 seconds (Sage; for this the total runtime for all three runs divided by three, as Sage was executed once and processed all runs jointly) to several hours (MaxQuant and MS-GF+). In addition, scaling behavior differs markedly between algorithms. Multiple configurations per algorithm were evaluated to optimize performance with respect to execution time across the three MS runs. Comet and Sage scale efficiently, utilizing up to 16 CPU cores for most of the runtime while requiring comparatively little memory. MSFragger, MS Amanda, and X!Tandem were restricted to 8 cores, which were generally well utilized, albeit in some cases with high memory consumption. MaxQuant shows limited per-run scalability and mainly operates on a single core. MS-GF+ was restricted to 6 cores, as it did not exceed this level of parallelization in any test, even when additional cores were available. Instead, improved throughput was achieved by splitting raw files and processing them in parallel.

## Conclusion

In this study, we systematically evaluated the impact of different rescoring strategies across multiple publicly available peptide identification algorithms. This assessment was performed on several datasets, including both real-world and benchmarking data acquired on different mass spectrometry platforms, and across protein databases of varying size and composition.

Analysis of the target–decoy distributions demonstrated that, in most cases, the necessary assumption of similar score distributions for decoy PSMs and incorrect target PSMs was visually inspected and could not be ruled out. This remained true for the majority of rescoring approaches. However, in certain combinations, the inclusion of all available features led to overly strong separation between targets and decoys, resulting in decoys no longer modeling incorrect identifications. This issue could be resolved by removing specific problematic features. These observations emphasize that, although rescoring approaches are generally robust, careful inspection of outputs and attention to algorithm warnings remain essential for ensuring reliable results.

Evaluation of FDR control using entrapment represents an additional important validation step. For most combinations of dataset, search engine, and rescoring strategy, FDR control appeared reasonable based on the estimated upper and lower FDP bounds. It should be noted that only these bounds were assessed. The computationally more expensive evaluation using paired target–decoy entrapment strategies was not performed in this study. Some combinations exhibited problematic behavior that require further investigation in practical applications. For example, in the CAMPI dataset, the FDP–FDR relationship for MS Amanda could not be reliably estimated due to high-scoring entrapment identifications. Nevertheless, the subsequent overlap analyses suggested that the resulting peptide identifications were largely consistent with those of other search engines. Overall, MS2Rescore and Oktoberfest showed a higher tendency toward FDR underestimation compared to raw PSM scoring and Percolator.

Inspection of the total number of identified peptidoforms revealed a clear trend. When relying solely on raw search engine results and to a lesser extent on Percolator alone, substantial variability in identification counts was observed between search engines. This variability was markedly reduced when applying prediction-based rescoring methods, resulting in increased consensus across engines, although some outliers remained. MaxQuant generally produced lower identification numbers under the tested conditions. It should be noted, however, that “match between runs” (MBR) was not enabled, which may influence performance for this workflow and is commonly used in quantitative analyses. But also none of the other search engines used anything similar. Sage, the most recently introduced search engine evaluated here, consistently yielded high identification numbers. MS-GF+ though achieved the highest identification rates overall. Although X!Tandem’s numbers of identifications were underwhelming with raw PSM scoring for FDR estimation in some datasets, the identifications could be rescued after rescoring.

The protein database selection had different effects on the datasets. For the CAMPI dataset, the larger protein database DB1 yielded substantially more identifications, most of which overlapped with those obtained using the smaller database DB2. This suggests that the smaller database may not have represented all proteins present in the sample, whereas the larger database resulted in higher identification coverage. Such challenges are common in metaproteomics and highlight the importance of appropriate database selection. In contrast, for the human datasets, identification numbers were relatively stable across databases of different sizes. The Swiss-Prot database with isoforms appeared sufficiently comprehensive for most applications tested here. However, this assessment focuses on identification counts rather than identification quality or biological relevance. In practical applications, databases such as the reference proteome or ProHap may include protein variants not present in Swiss-Prot that are critical for specific studies. Given that identification numbers did not decrease when using the reference proteome,its use is recommended for general applications.

Because rescoring strategies substantially mitigate differences in identification counts between search engines, the choice of algorithm may depend more on workflow integration, computational requirements, and resource availability than on raw identification performance. In terms of computational efficiency, MS-GF+ required the longest processing times, even when database and dataset splitting strategies were applied. Sage and MSFragger were generally the fastest engines, although both require substantial memory when searching large databases. Database splitting can reduce memory demands for these tools, but at the cost of increased computation time. Comet proved robust across different hardware configurations, although its runtime depends on the available CPU and memory resources.

The present results represent a snapshot under specific configurations. In this study, only tryptic peptides were inspected, while the behavior might change for datasets using different or no cleavage agents. Also, only a small subset of MS machines (Orbitrap and TIMS-TOF) was analysed, as were two of the main modifications, oxidation of M and carbamidomethylation of C. To facilitate ongoing benchmarking, we plan to implement entrapment assessment modules and standardized identification workflows within ProteoBench^36^. ProteoBench is an open and collaborative platform for community-curated benchmarks for proteomics data analysis pipelines. It allows the upload of quantification and identification results as datapoints for different modules, which inspect the results under various angles. This enables continuous and updated comparisons of tools and pipelines, not only at the time point of e.g.a tool’s release or publication. The developed Nextflow pipeline for MS/MS identification (mspepid) has been accepted as an nf-core workflow^37^ and the development version is available on the nf-core repository (https://nf-co.re/mspepid/).

In summary, our results demonstrate that substantial differences in identification numbers observed between search engines applying conventional FDR estimation with only the raw search results are largely mitigated by advanced, prediction-driven rescoring strategies. Once additional informative features are incorporated, consensus across engines increases markedly. Nevertheless, rescoring does not diminish the need for careful validation: inspection of target–decoy score distributions and independent assessment of FDR control, for example via entrapment analyses, remain essential to ensure reliable results. Importantly, the convergence of identification counts suggests that the relevant peptide identifications are, in principle, accessible to all evaluated search engines. The results suggest that the discriminatory power of the initial scoring functions, rather than PSM candidate generation, may be a primary limitation in achieving higher consensus across search engines. This highlights the value of improved scoring models and approaches that more directly integrate predicted spectral information, which may further enhance robustness and reliability in future proteomics workflows.

## Acknowledgements

The authors thank the Medical Faculty of the Ruhr University Bochum for the funding of the Core Unit Bioinformatics - CUBiMed.RUB and the possibility of using the hardware provided.

## Data Availability

All data and code are available in public repositories.

The Nextflow workflow^19^ for the peptide identification and rescoring is available at https://github.com/medbioinf/mspepid.

All scripts for the analysis and plotting of the data are available on GitHub (https://github.com/medbioinf/mspepid-analysis/tree/main).

The reanalyzed data are uploaded to PRIDE using the following identifiers:

- CAMPI: PXD075141,
- Cancer Array: PXD075199,
- Orbitrap: PXD075209,
- timsTOF: PXD075206.

## References

(1) Käll, L.; Canterbury, J. D.; Weston, J.; Noble, W. S.; MacCoss, M. J. Semi-supervised learning for peptide identification from shotgun proteomics datasets. Nature Methods 2007, 4, 923–925, DOI: 10.1038/nmeth1113.

(2) Keller, A.; Nesvizhskii, A. I.; Kolker, E.; Aebersold, R. Empirical Statistical Model To Estimate the Accuracy of Peptide Identifications Made by MS/MS and Database Search. Analytical Chemistry 2002, 74, 5383–5392, DOI: 10.1021/ac025747h.

(3) Elias, J. E.; Gygi, S. P. Target-decoy search strategy for increased confidence in large-scale protein identifications by mass spectrometry. Nature Methods 2007, 4, 207–214, DOI: 10.1038/nmeth1019.

(4) Declercq, A.; Bouwmeester, R.; Hirschler, A.; Carapito, C.; Degroeve, S.; Martens, L.; Gabriels, R. MS2Rescore: Data-Driven Rescoring Dramatically Boosts Immunopeptide Identification Rates. Molecular & Cellular Proteomics 2022, 21, 100266, DOI: 10.1016/j.mcpro.2022.100266.

(5) Picciani, M.; Gabriel, W.; Giurcoiu, V.-G.; Shouman, O.; Hamood, F.; Lautenbacher, L.; Jensen, C. B.; Müller, J.; Kalhor, M.; Soleymaniniya, A.; Kuster, B.; The, M.; Wilhelm, M. Oktoberfest: Open-source spectral library generation and rescoring pipeline based on Prosit. PROTEOMICS 2024, 24, 2300112, DOI: 10.1002/pmic.202300112.

(6) Degroeve, S.; Martens, L. MS2PIP: a tool for MS/MS peak intensity prediction. Bioinformatics2013, 29, 3199–3203, DOI: 10.1093/bioinformatics/btt544.

(7) Gessulat, S.; Schmidt, T.; Zolg, D. P.; Samaras, P.; Schnatbaum, K.; Zerweck, J.; Knaute, T.; Rechenberger, J.; Delanghe, B.; Huhmer, A.; Reimer, U.; Ehrlich, H.-C.; Aiche, S.; Kuster, B.; Wilhelm, M. Prosit: proteome-wide prediction of peptide tandem mass spectra by deep learning. Nature Methods 2019, 16, 509–518, DOI: 10.1038/s41592-019-0426-7.

(8) Fondrie, W. E.; Noble, W. S. mokapot: Fast and Flexible Semisupervised Learning for Peptide Detection. Journal of Proteome Research 2021, 20, 1966–1971, DOI: 10.1021/acs.jproteome.0c01010.

(9) Verbruggen, S.; Gessulat, S.; Gabriels, R.; Matsaroki, A.; Van de Voorde, H.; Kuster, B.; Degroeve, S.; Martens, L.; Van Criekinge, W.; Wilhelm, M.; Menschaert, G. Spectral Prediction Features as a Solution for the Search Space Size Problem in Proteogenomics. Molecular & Cellular Proteomics 2021, 20, 100076, DOI: 10.1016/j.mcpro.2021.100076.

(10) Castaño, J. D.; Beaudry, F. Comparative Analysis of Data-Driven Rescoring Platforms for Improved Peptide Identification in HeLa Digest Samples. PROTEOMICS 2025, 25, e202400225, DOI: 10.1002/pmic.202400225.

(11) Cormican, J. A.; Horokhovskyi, Y.; Soh, W. T.; Mishto, M.; Liepe, J. inSPIRE: An Open-Source Tool for Increased Mass Spectrometry Identification Rates Using Prosit Spectral Prediction. Molecular & Cellular Proteomics 2022, 21, 100432, DOI: 10.1016/j.mcpro.2022.100432.

(12) Craig, R.; Beavis, R. C. TANDEM: matching proteins with tandem mass spectra. Bioinformatics 2004, 20, 1466–1467, DOI: 10.1093/bioinformatics/bth092.

(13) Cox, J.; Neuhauser, N.; Michalski, A.; Scheltema, R. A.; Olsen, J. V.; Mann, M. Andromeda: A Peptide Search Engine Integrated into the MaxQuant Environment. Journal of Proteome Research 2011, 10, 1794–1805, DOI: 10.1021/pr101065j.

(14) Eng, J. K.; Jahan, T. A.; Hoopmann, M. R. Comet: An open-source MS/MS sequence database search tool. PROTEOMICS 2013, 13, 22–24, DOI: 10.1002/pmic.201200439.

(15) Dorfer, V.; Pichler, P.; Stranzl, T.; Stadlmann, J.; Taus, T.; Winkler, S.; Mechtler, K. MS Amanda, a Universal Identification Algorithm Optimized for High Accuracy Tandem Mass Spectra. Journal of Proteome Research 2014, 13, 3679–3684, DOI: 10.1021/pr500202e.

(16) Kim, S.; Pevzner, P. A. MS-GF+ makes progress towards a universal database search tool for proteomics. Nature Communications 2014, 5, 5277, DOI: 10.1038/ncomms6277.

(17) Kong, A. T.; Leprevost, F. V.; Avtonomov, D. M.; Mellacheruvu, D.; Nesvizhskii, A. I. MSFragger: ultrafast and comprehensive peptide identification in mass spectrometry–based proteomics. Nature Methods 2017, 14, 513–520, DOI: 10.1038/nmeth.4256.

(18) Lazear, M. R. Sage: An Open-Source Tool for Fast Proteomics Searching and Quantification at Scale. Journal of Proteome Research 2023, 22, 3652–3659, DOI: 10.1021/acs.jproteome.3c00486.

(19) Di Tommaso, P.; Chatzou, M.; Floden, E. W.; Barja, P. P.; Palumbo, E.; Notredame, C. Nextflow enables reproducible computational workflows. Nature Biotechnology 2017, 35, 316–319, DOI: 10.1038/nbt.3820.

(20) Chambers, M. C. et al. A cross-platform toolkit for mass spectrometry and proteomics. Nature Biotechnology 2012, 30, 918–920, DOI: 10.1038/nbt.2377.

(21) Pfeuffer, J. et al. OpenMS 3 enables reproducible analysis of large-scale mass spectrometry data. Nature Methods 2024, 21, 365–367, DOI: 10.1038/s41592-024-02197-7.

(22) He, K.; Fu, Y.; Zeng, W.-F.; Luo, L.; Chi, H.; Liu, C.; Qing, L.-Y.; Sun, R.-X.; He, S.-M. A theoretical foundation of the target-decoy search strategy for false discovery rate control in proteomics. arXiv (Statistics) 2015, DOI: 10.48550/arXiv.1501.00537.

(23) Wen, B.; Freestone, J.; Riffle, M.; MacCoss, M. J.; Noble, W. S.; Keich, U. Assessment of false discovery rate control in tandem mass spectrometry analysis using entrapment. Nature Methods 2025, 22, 1454–1463, DOI: 10.1038/s41592-025-02719-x.

(24) Gabriels, R.; Declercq, A.; Bouwmeester, R.; Degroeve, S.; Martens, L. psm_utils: A High-Level Python API for Parsing and Handling Peptide-Spectrum Matches and Proteomics Search Results. Journal of Proteome Research 2023, 22, 557–560, DOI: 10.1021/acs.jproteome.2c00609.

(25) Declercq, A. et al. TIMS^2^ Rescore: A Data Dependent Acquisition-Parallel Accumulation and Serial Fragmentation-Optimized Data-Driven Rescoring Pipeline Based on MS^2^ Rescore. Journal of Proteome Research 2025, 24, 1067–1076, DOI: 10.1021/acs.jproteome.4c00609.

(26) Vašíček, J.; Kuznetsova, K. G.; Skiadopoulou, D.; Unger, L.; Chera, S.; Ghila, L. M.; Bandeira, N.; Njølstad, P. R.; Johansson, S.; Bruckner, S.; Käll, L.; Vaudel, M. ProHap enables human proteomic database generation accounting for population diversity. Nature Methods 2025, 22, 273–277, DOI: 10.1038/s41592-024-02506-0.

(27) Deutsch, E. W. et al. The ProteomeXchange consortium in 2017: supporting the cultural change in proteomics public data deposition. Nucleic Acids Research 2017, 45, D1100–D1106, DOI: 10.1093/nar/gkw936.

(28) Perez-Riverol, Y.; Bandla, C.; Kundu, D. J.; Kamatchinathan, S.; Bai, J.; Hewapathirana, S.; John, N. S.; Prakash, A.; Walzer, M.; Wang, S.; Vizcaíno, J. A. The PRIDE database at 20 years: 2025 update. Nucleic Acids Research 2025, 53, D543–D553, DOI: 10.1093/nar/gkae1011.

(29) Van Puyvelde, B. et al. A comprehensive LFQ benchmark dataset on modern day acquisition strategies in proteomics. Scientific Data 2022, 9, 126, DOI: 10.1038/s41597-022-01216-6.

(30) Tüshaus, J. et al. Towards routine proteome profiling of FFPE tissue: insights from a 1,220-case pan-cancer study. The EMBO Journal 2024, 44, 304–329, DOI: 10.1038/s44318-024-00289-w.

(31) The UniProt Consortium et al. UniProt: the Universal Protein Knowledgebase in 2025. Nucleic Acids Research 2025, 53, D609–D617, DOI: 10.1093/nar/gkae1010.

(32) Frankenfield, A. M.; Ni, J.; Ahmed, M.; Hao, L. Protein Contaminants Matter: Building Universal Protein Contaminant Libraries for DDA and DIA Proteomics. Journal of Proteome Research 2022, 21, 2104–2113, DOI: 10.1021/acs.jproteome.2c00145.

(33) Van Den Bossche, T. et al. Critical Assessment of MetaProteome Investigation (CAMPI): a multilaboratory comparison of established workflows. Nature Communications 2021, 12, 7305, DOI: 10.1038/s41467-021-27542-8.

(34) Li, J. et al. An integrated catalog of reference genes in the human gut microbiome. Nature Biotechnology 2014, 32, 834–841, DOI: 10.1038/nbt.2942.

(35) Debrie, E.; Malfait, M.; Gabriels, R.; Declerq, A.; Sticker, A.; Martens, L.; Clement, L. Quality Control for the Target Decoy Approach for Peptide Identification. Journal of Proteome Research 2023, 22, 350–358, DOI: 10.1021/acs.jproteome.2c00423.

(36) Devreese, R. et al. ProteoBench: the community-curated platform for comparing proteomics data analysis workflows. bioRxiv 2025, ISSN: 2692-8205 Pages: 2025.12.09.692895 Section: New Results, DOI: 10.64898/2025.12.09.692895.

(37) Ewels, P. A.; Peltzer, A.; Fillinger, S.; Patel, H.; Alneberg, J.; Wilm, A.; Garcia, M. U.; Di Tommaso, P.; Nahnsen, S. The nf-core framework for community-curated bioinformatics pipelines. Nature Biotechnology 2020, 38, 276–278, DOI: 10.1038/s41587-020-0439-x.

