## supplemental data for "From variability to consensus: rescoring harmonizes peptide identification across diverse search engines and datasets"

This file contains supplemental plots for all analyzed combinations of search engines, databases, and datasets. Except for the target-decoy distribution plots, all figures present aggregated results across runs within each dataset.

For the pseudo-ROC and entrapment analyses, the graphs display the maximum and minimum values observed across all runs. The bar charts report the mean number of peptidoform identifications across runs, with whiskers indicating the corresponding standard deviation. UpSet plots show the mean number of peptidoforms per intersection group across runs.

As only a single run was assessed for the CAMPI dataset, the reported values correspond directly to that run.

The color scheme in the UpSet plots is intended solely to improve readability and reflects the degree of the respective intersection sets. Intersections comprising all sets are shown in red, while degree-one intersections ("present in only one set") are displayed in blue. In Chapter 4, all remaining intersection degrees are colored violet. In Chapter 5, degree-six intersections ("all search engines except one") are shown in violet, whereas intersections of degrees two to five are displayed in green.

**6 Entrapment Analyses** 45

**7 Software tools** 50

### 1 Target-decoy distribution plots

These plots illustrate the distribution of target and decoy PSMs used for FDR estimation. Each column corresponds to an individual run (the number of runs varies by dataset), while each row represents a specific search engine. Within each plot, search engines are ordered as follows: Comet, MaxQuant, MS Amanda, MSFragger, MS-GF+, Sage, and X!Tandem.

#### 1.1 Cancer Array Dataset

##### 1.1.1 Swiss-Prot

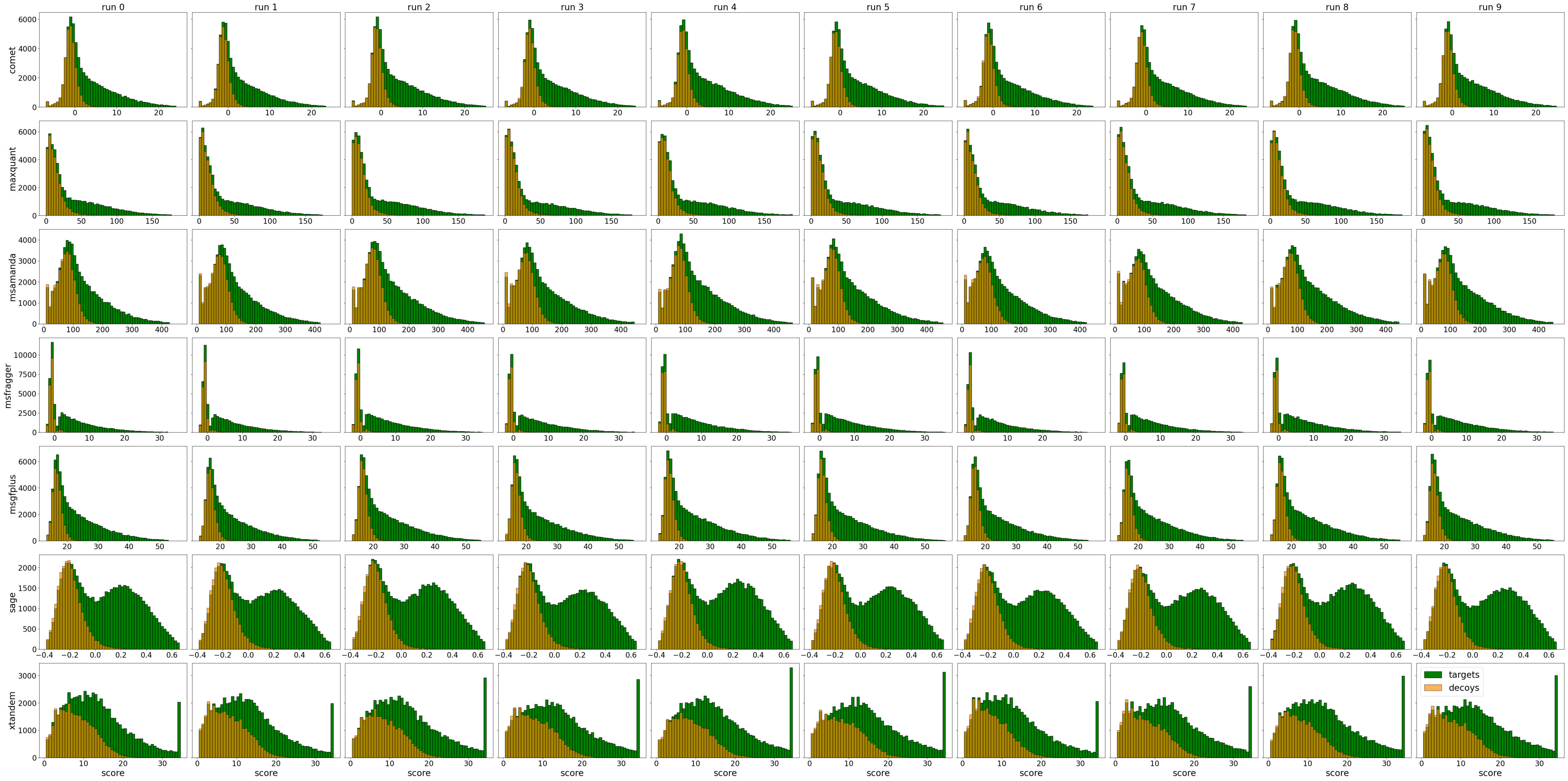

Figure S1: Target-decoy distribution, Cancer Array, Swiss-Prot, raw search results

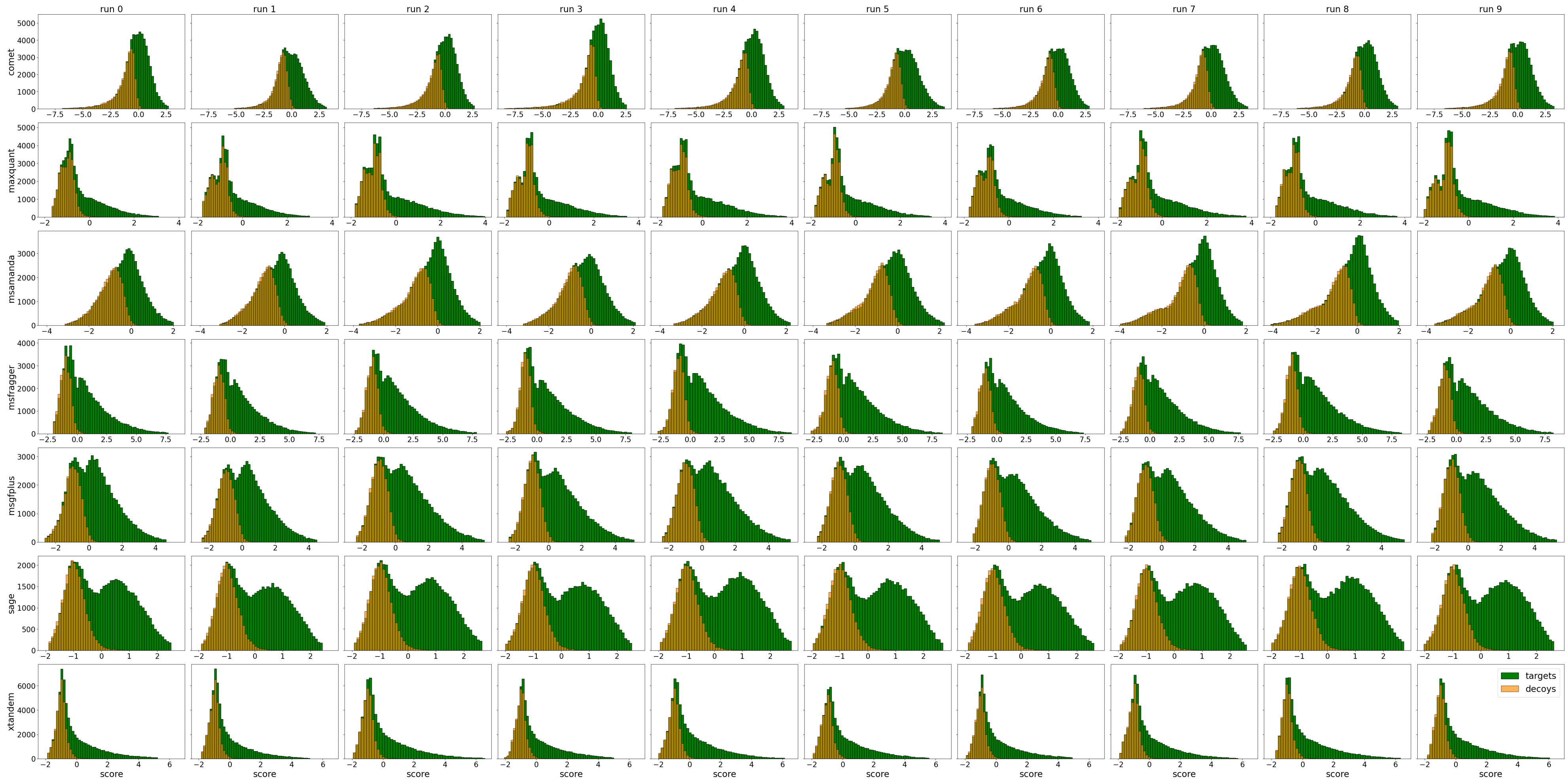

Figure S2: Target-decoy distribution, Cancer Array, Swiss-Prot, Percolator

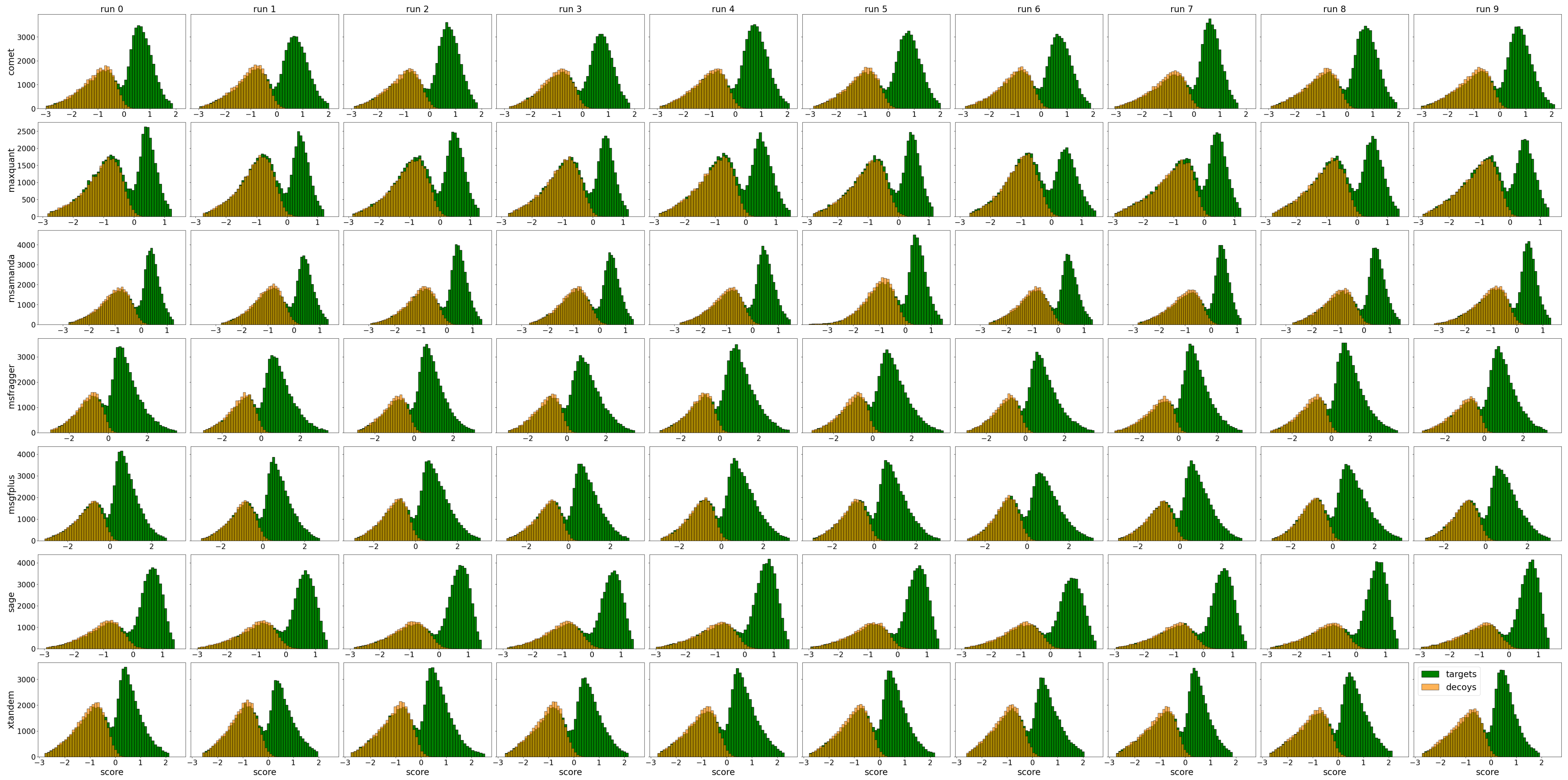

Figure S3: Target-decoy distribution, Cancer Array, Swiss-Prot, MS2Rescore

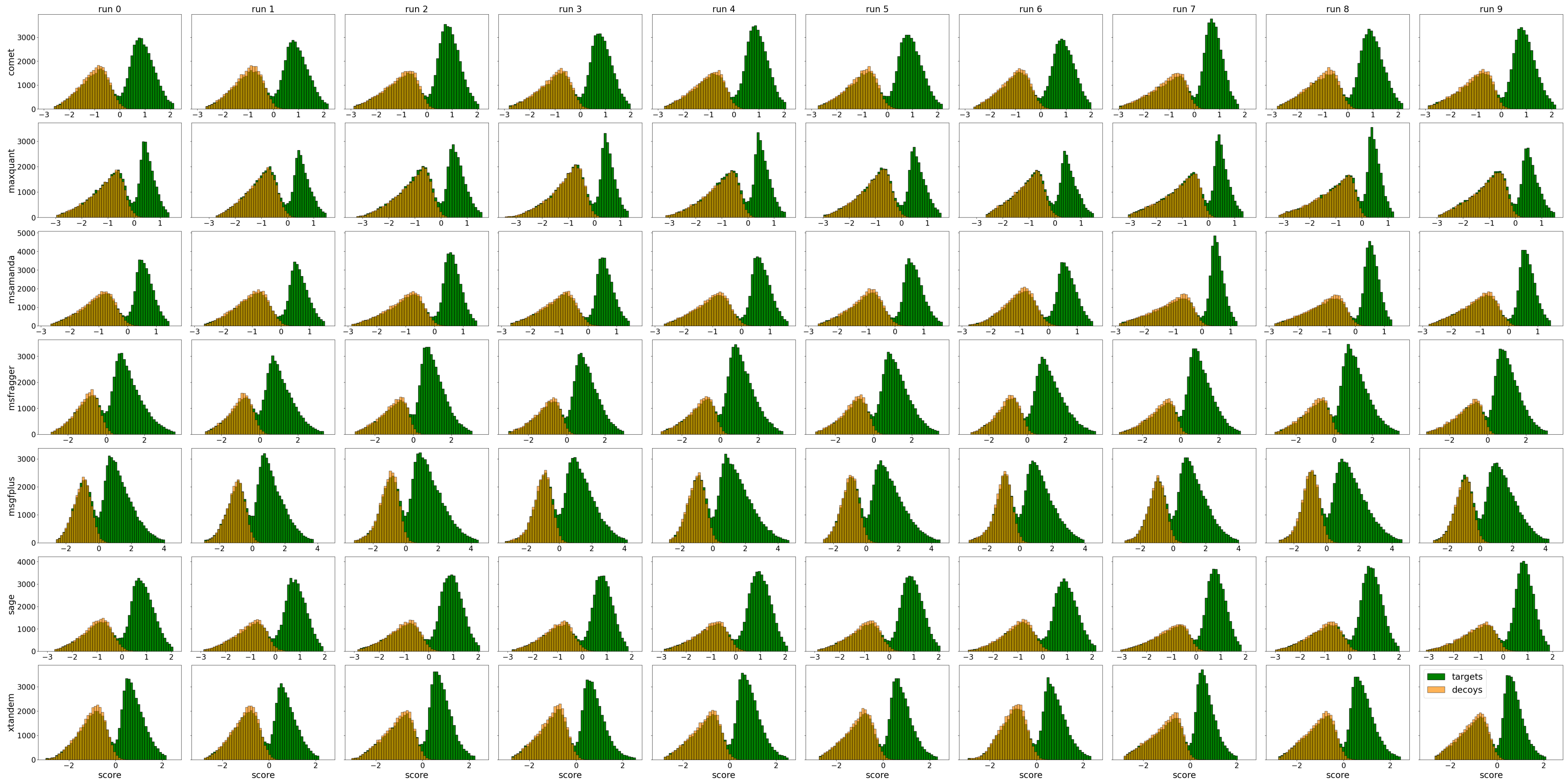

Figure S4: Target-decoy distribution, Cancer Array, Swiss-Prot, Oktoberfest

##### 1.1.2 Proteome

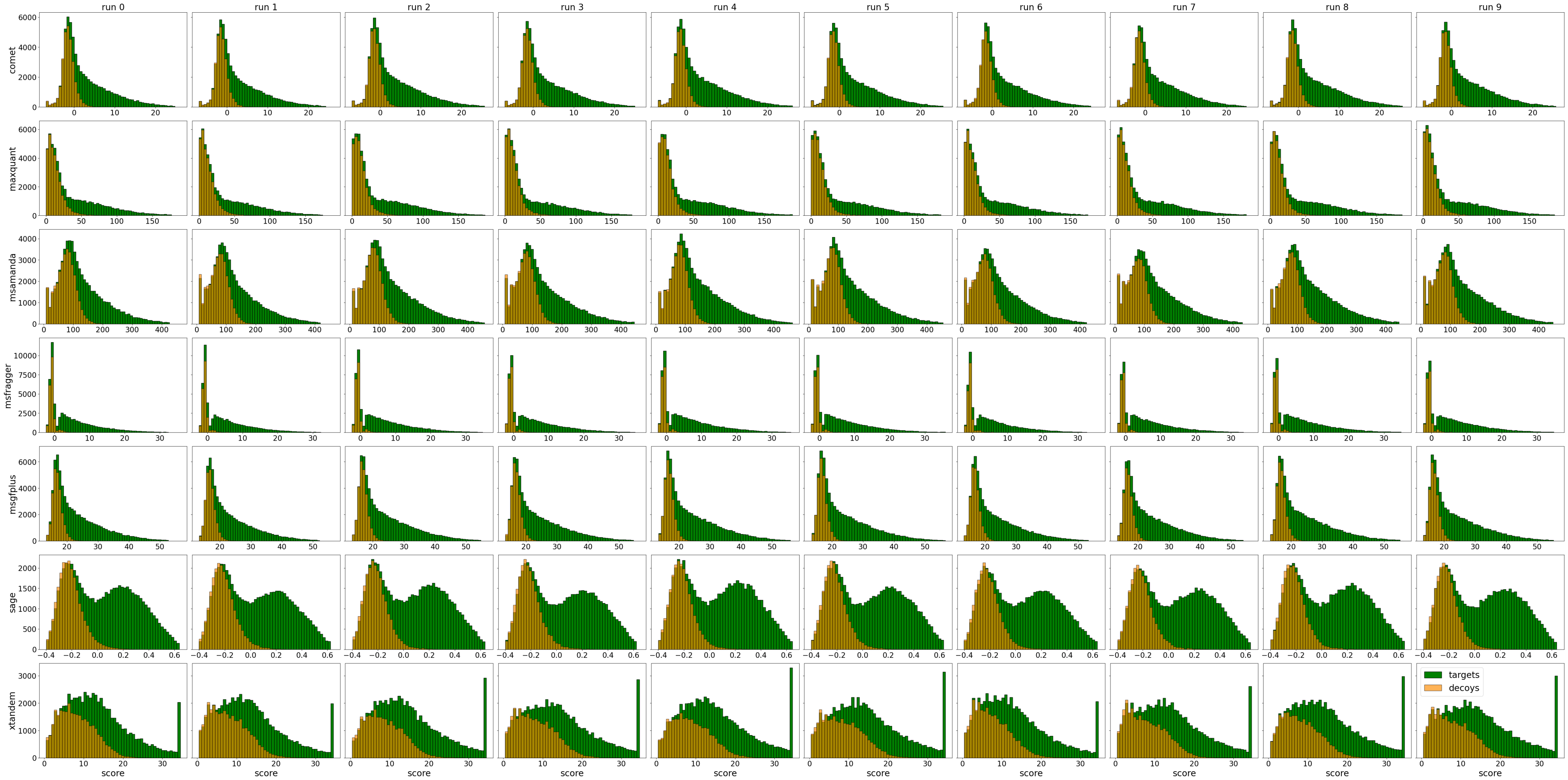

Figure S5: Target-decoy distribution, Cancer Array, Proteome, raw search results

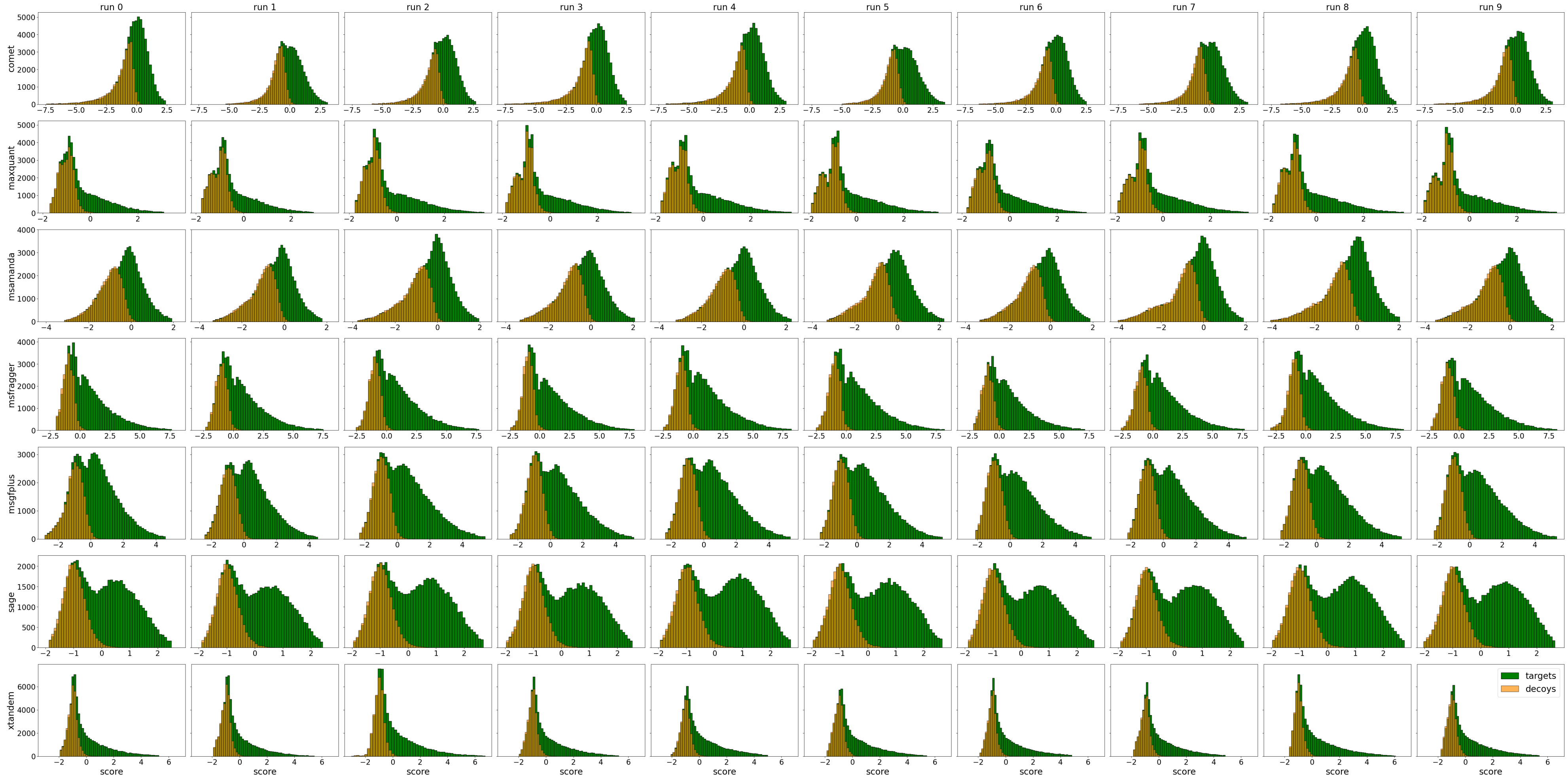

Figure S6: Target-decoy distribution, Cancer Array, Proteome, Percolator

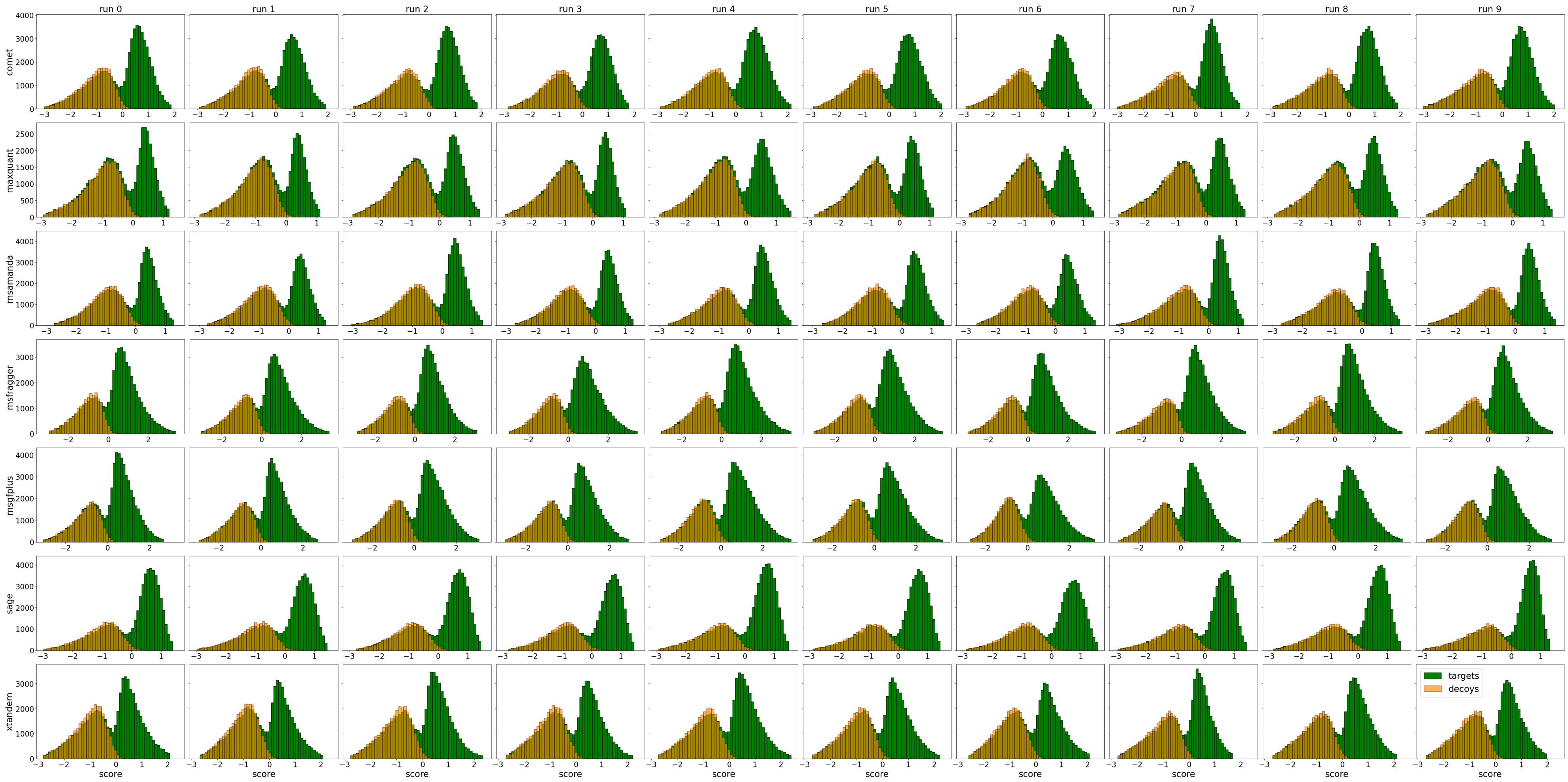

Figure S7: Target-decoy distribution, Cancer Array, Proteome, MS2Rescore

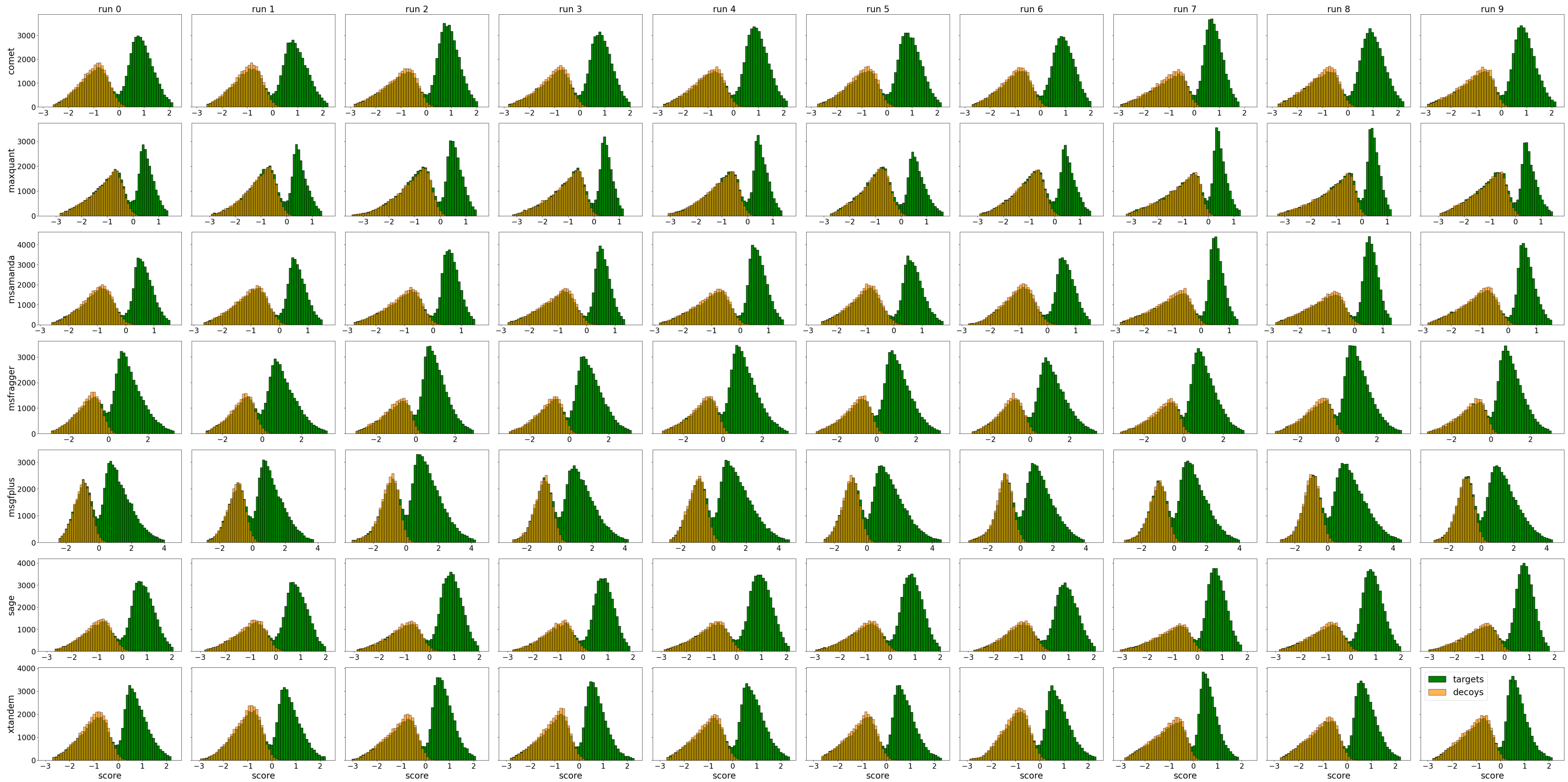

Figure S8: Target-decoy distribution, Cancer Array, Proteome, Oktoberfest

##### 1.1.3 ProHap

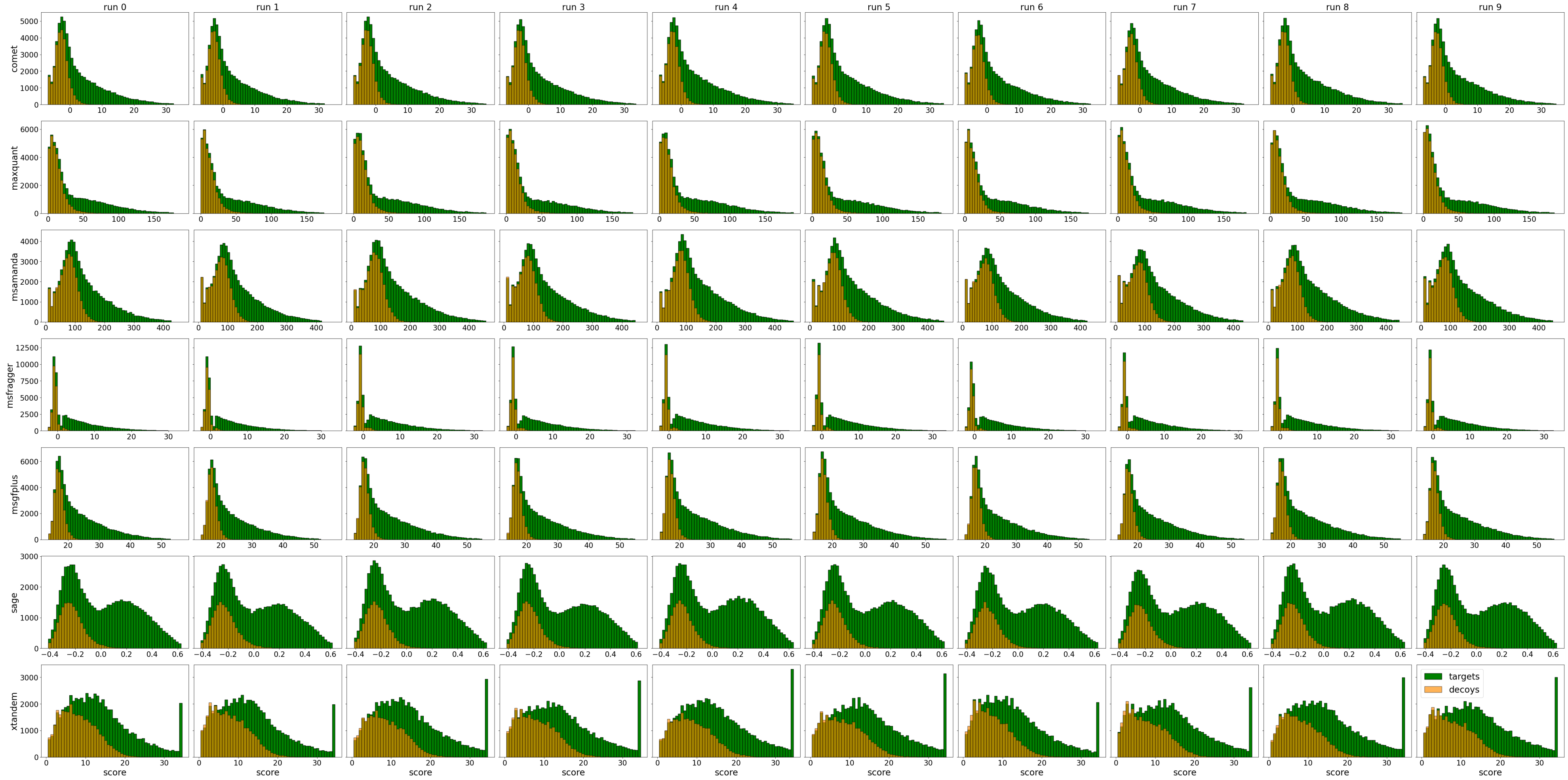

Figure S9: Target-decoy distribution, Cancer Array, ProHap, raw search results

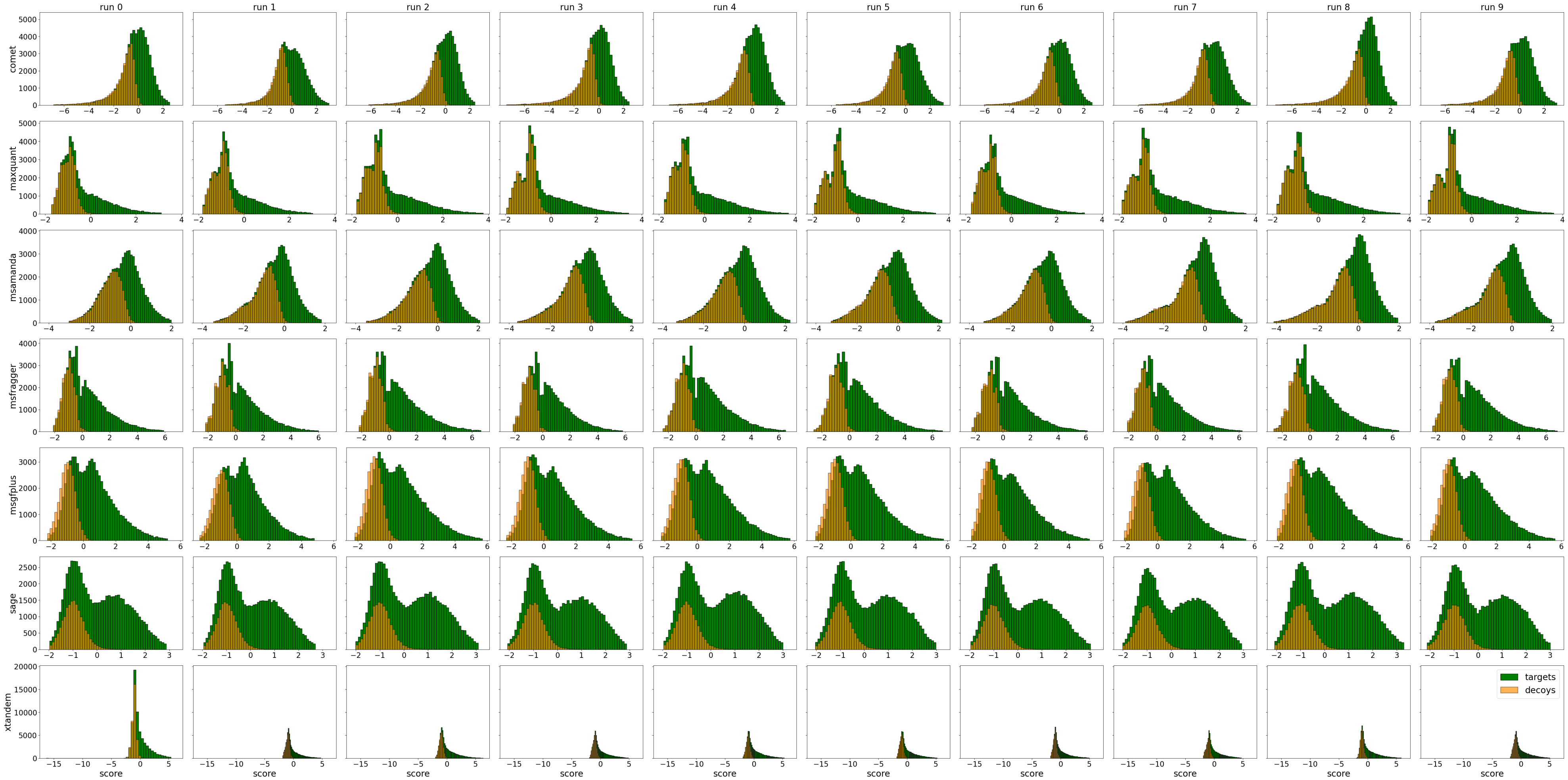

Figure S10: Target-decoy distribution, Cancer Array, ProHap, Percolator

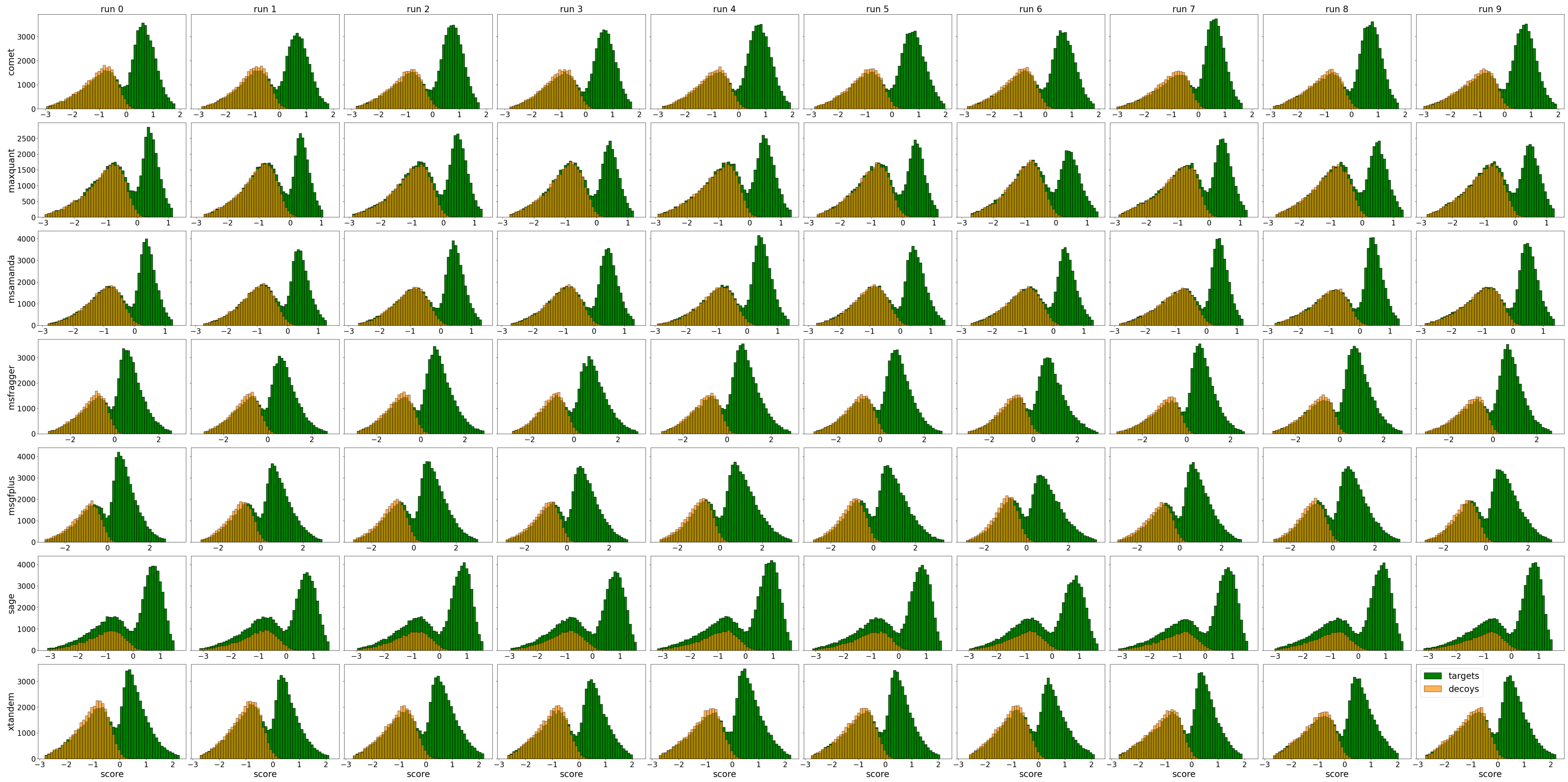

Figure S11: Target-decoy distribution, Cancer Array, ProHap, MS2Rescore

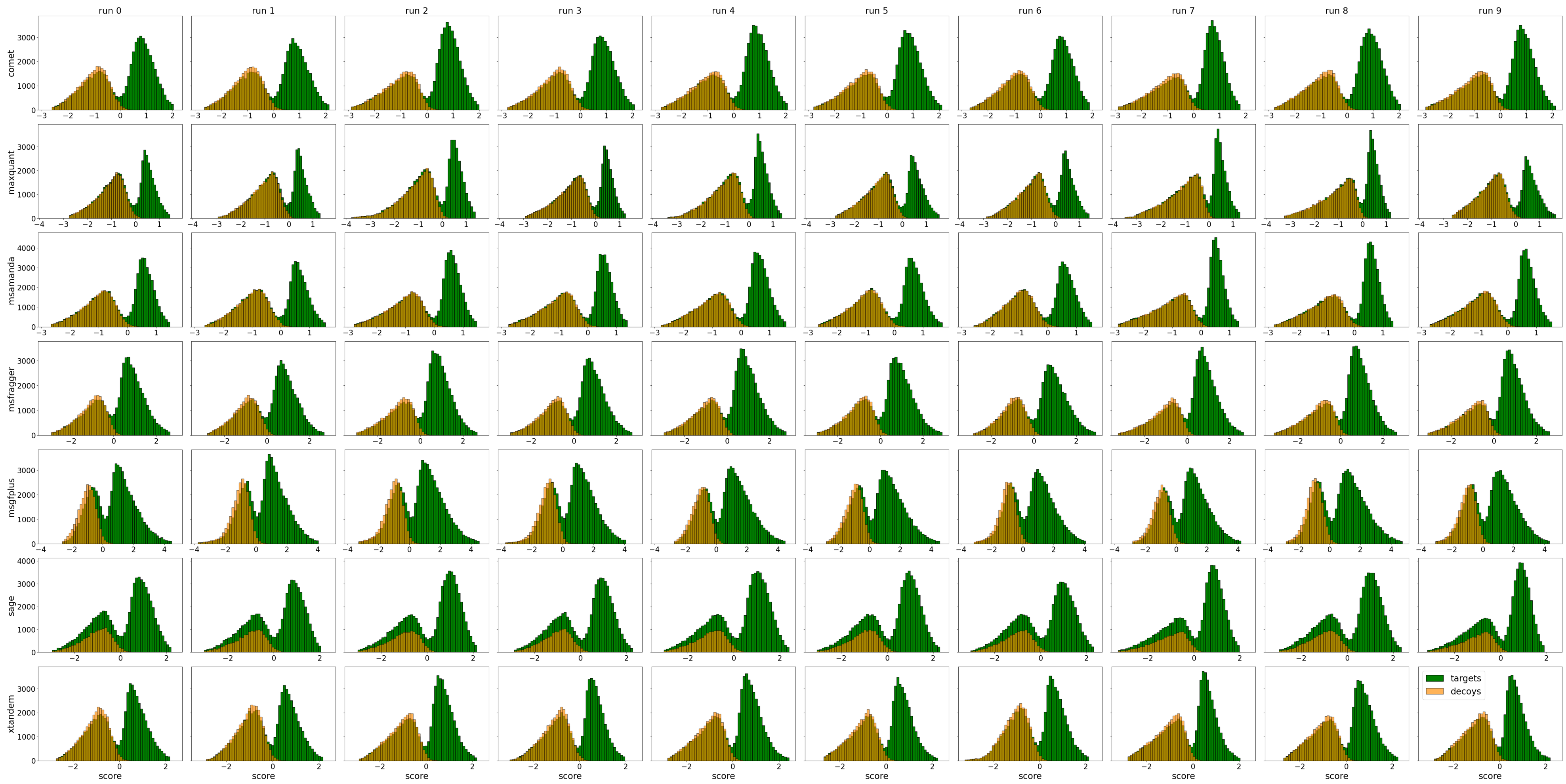

Figure S12: Target-decoy distribution, Cancer Array, ProHap, Oktoberfest

#### 1.2 CAMPI Dataset

### 1.2.1 DB1

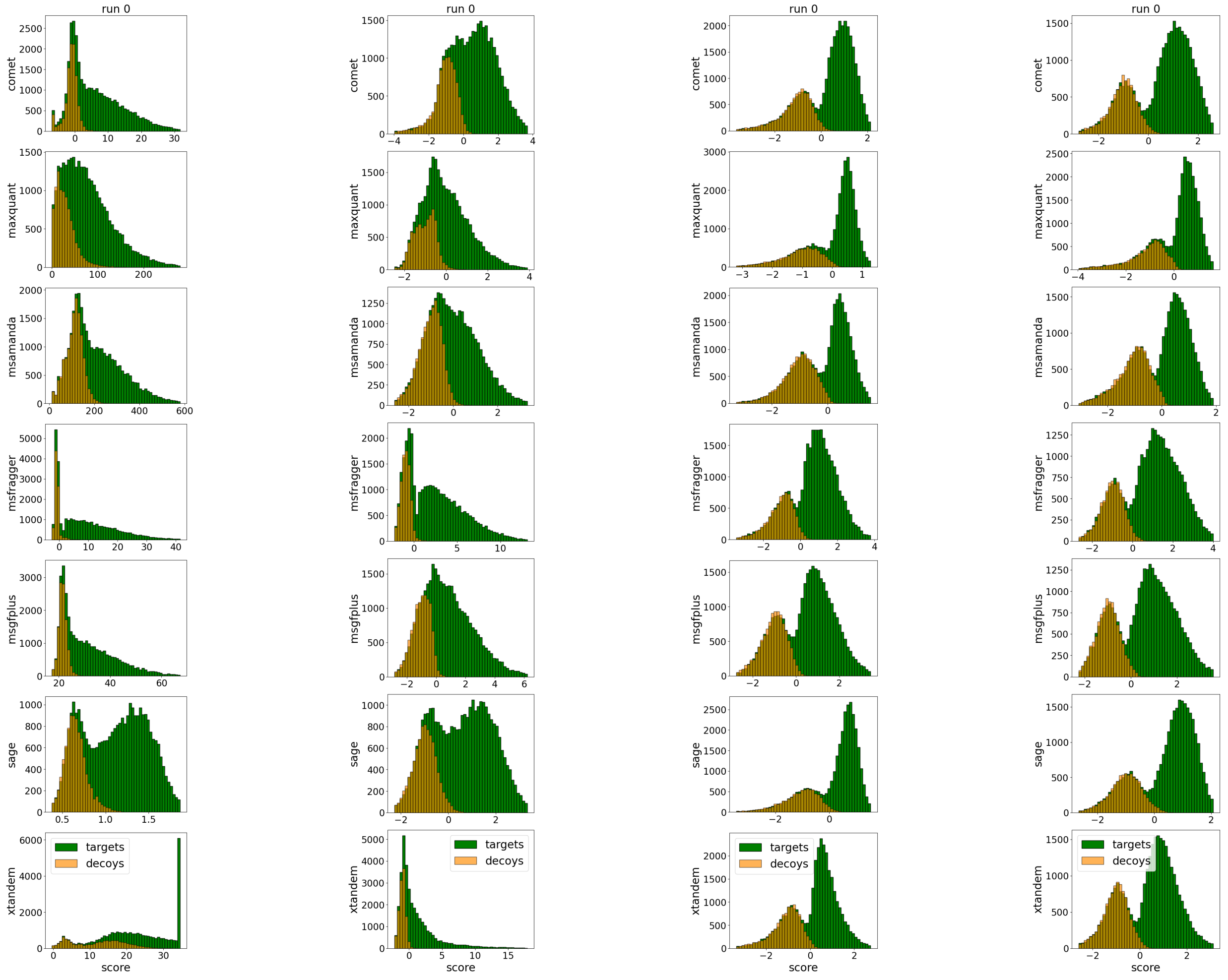

(a) raw search results (b) Percolator (c) MS2Rescore (d) Oktoberfest

Figure S13: Target-decoy distribution, CAMPI, DB1

### 1.2.2 DB2

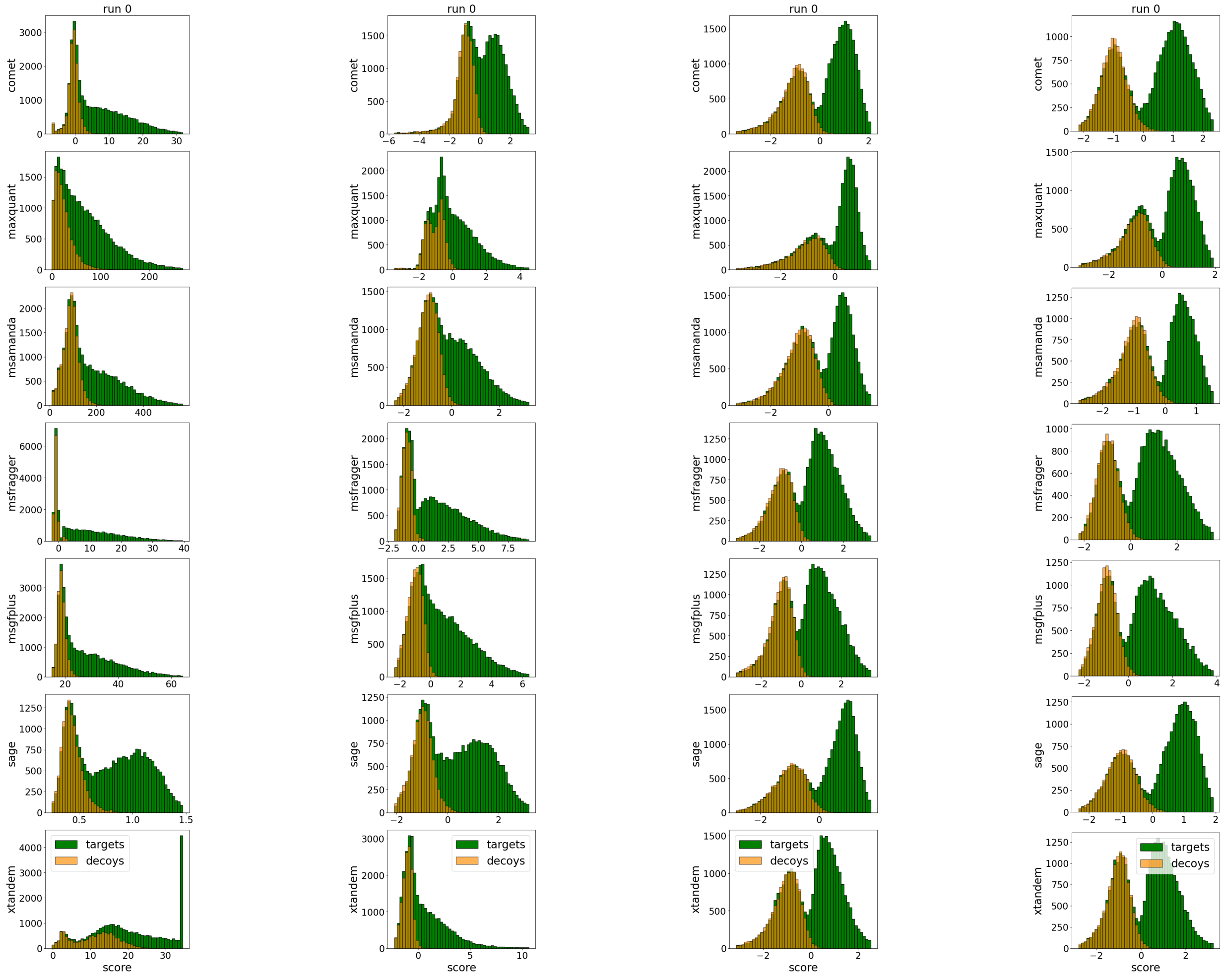

(a) raw search results (b) Percolator (c) MS2Rescore (d) Oktoberfest

Figure S14: Target-decoy distribution, CAMPI, DB2

#### 1.3 Orbitrap Dataset

##### 1.3.1 Swiss-Prot

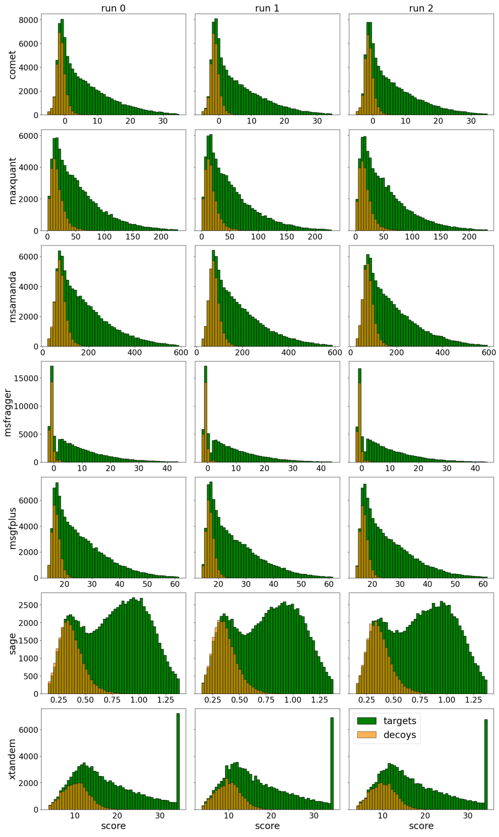

(a) raw search results

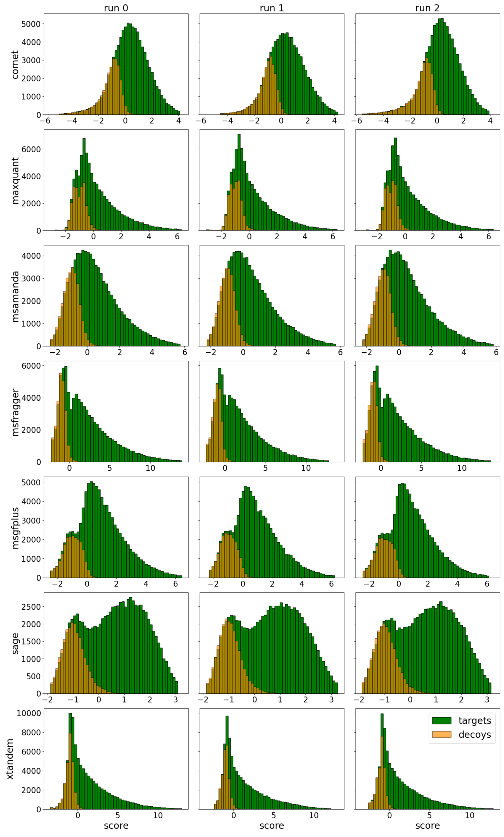

(b) Percolator

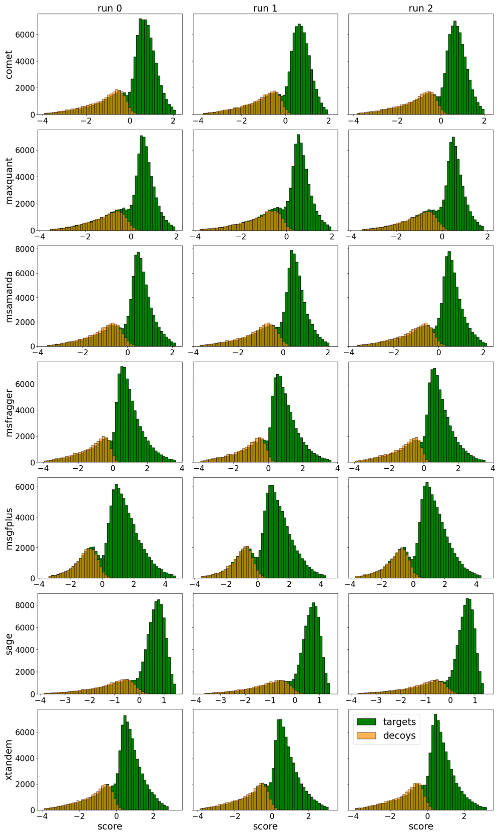

(c) MS2Rescore

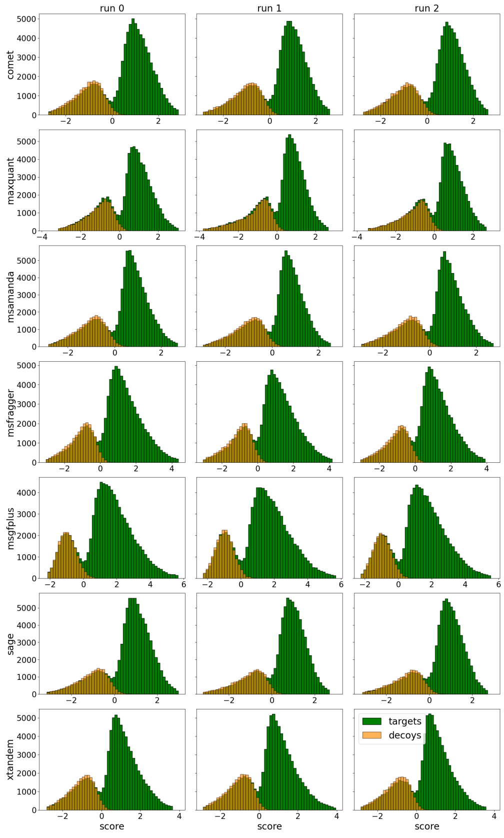

(d) Oktoberfest

Figure S15: Target-decoy distribution, Orbitrap dataset, Swiss-Prot

##### 1.3.2 Proteome

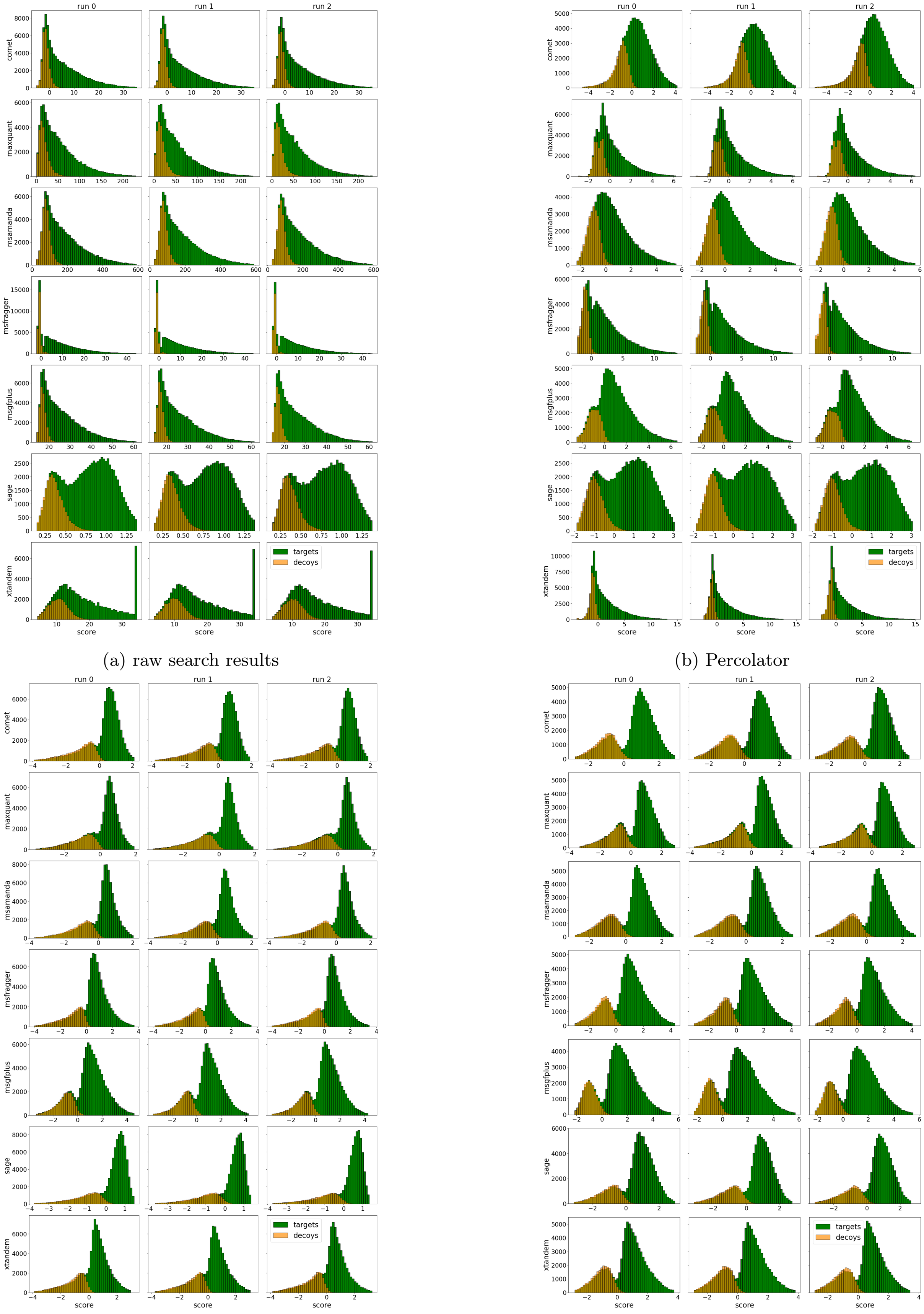

(a) raw search results (b) Percolator

(c) MS2Rescore (d) Oktoberfest

Figure S16: Target-decoy distribution, Orbitrap dataset, Proteome

##### 1.3.3 ProHap

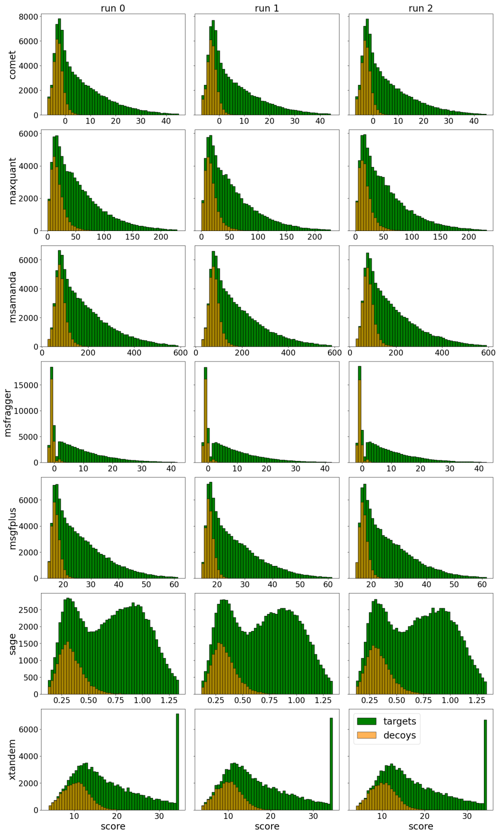

(a) raw search results

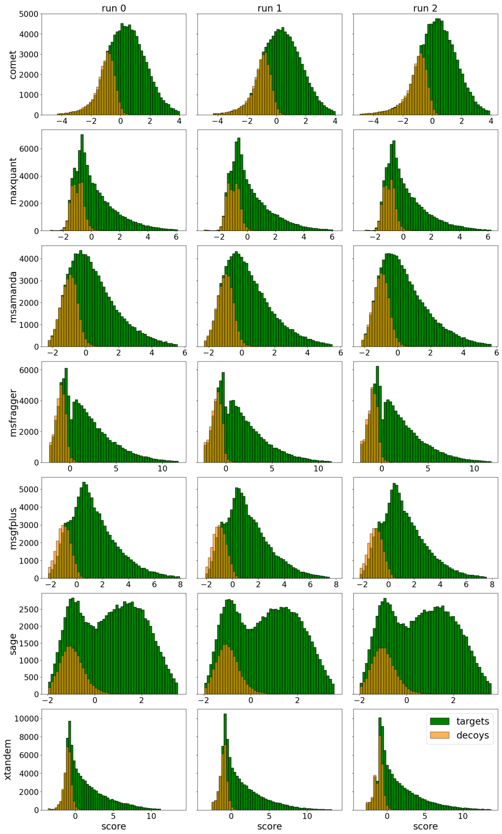

(b) Percolator

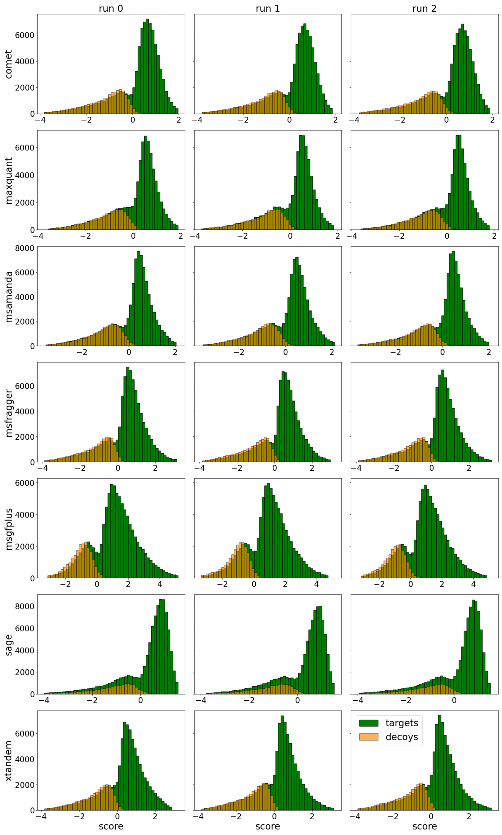

(c) MS2Rescore

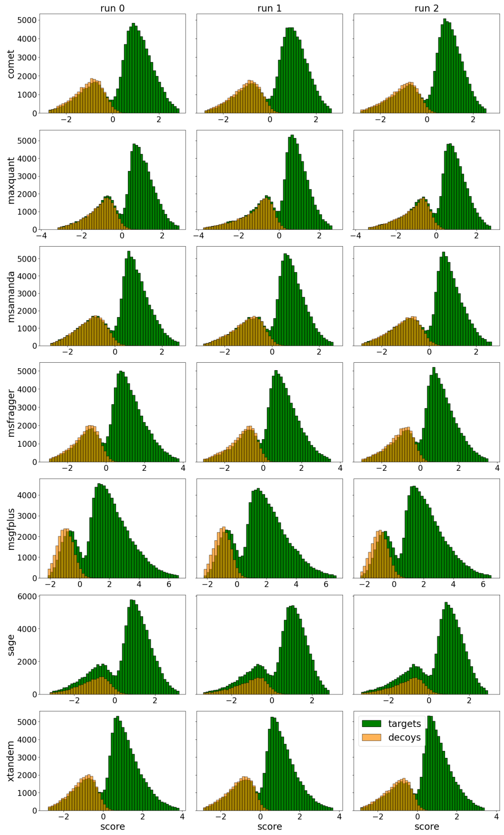

(d) Oktoberfest

Figure S17: Target-decoy distribution, Orbitrap dataset, ProHap

#### 1.4 timsTOF Dataset

##### 1.4.1 Swiss-Prot

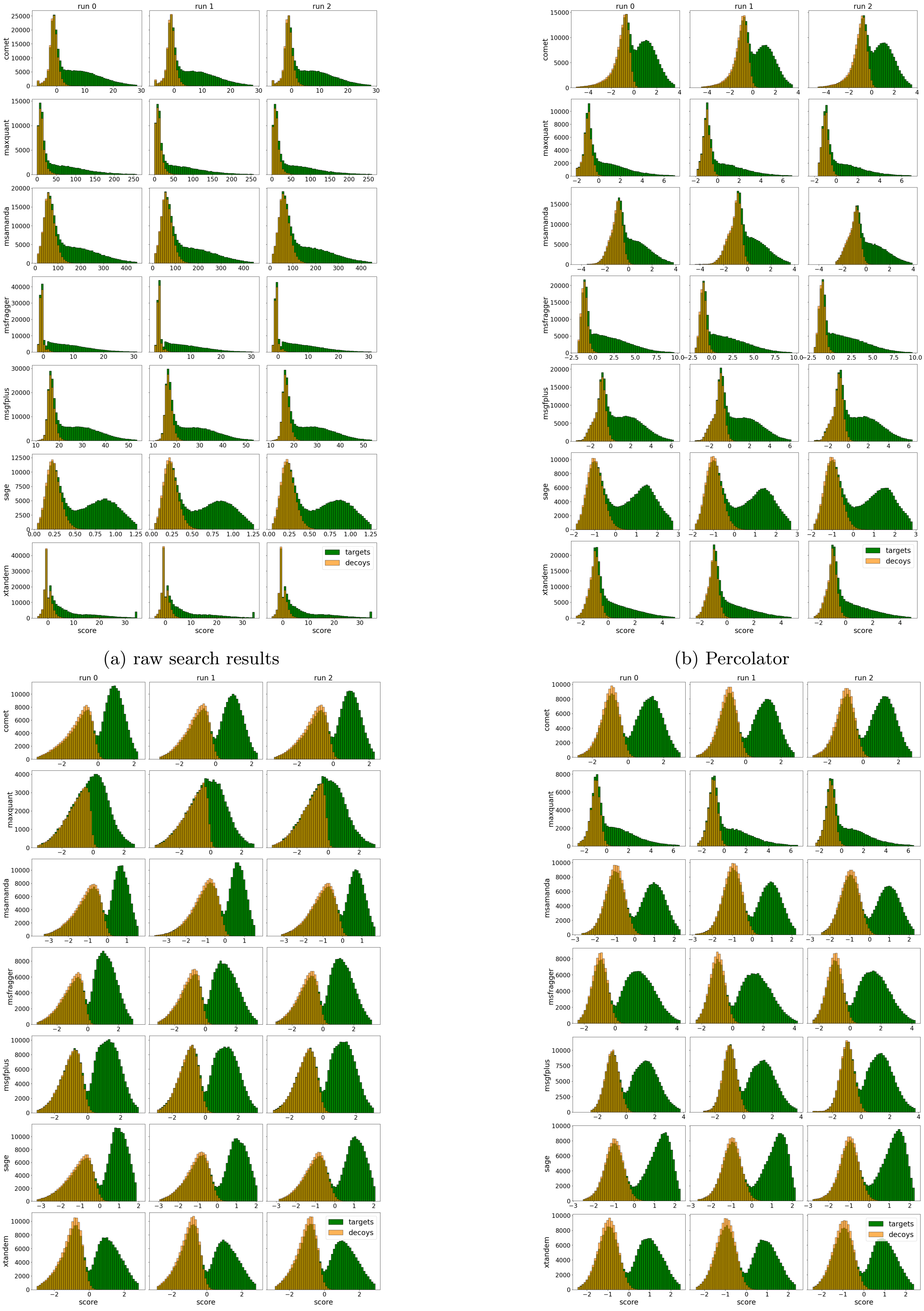

(a) raw search results

(b) Percolator

(c) MS2Rescore

(d) Oktoberfest

Figure S18: Target-decoy distribution, timsTOF dataset, Swiss-Prot

##### 1.4.2 Proteome

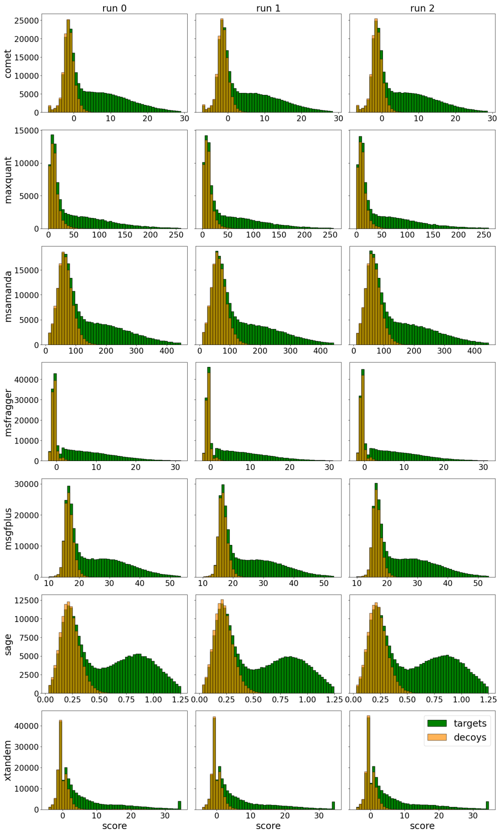

(a) raw search results

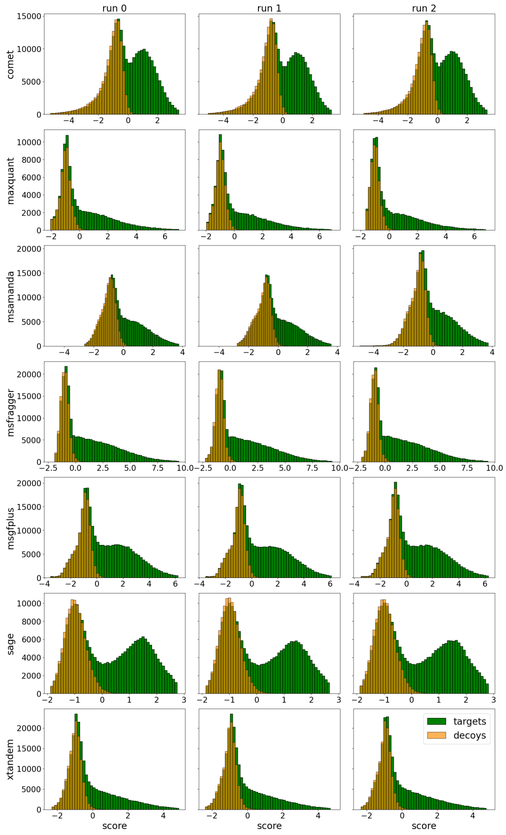

(b) Percolator

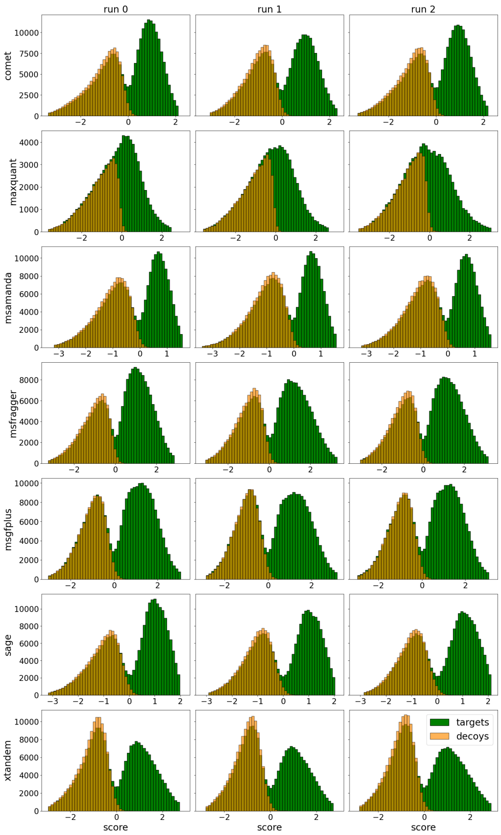

(c) MS2Rescore

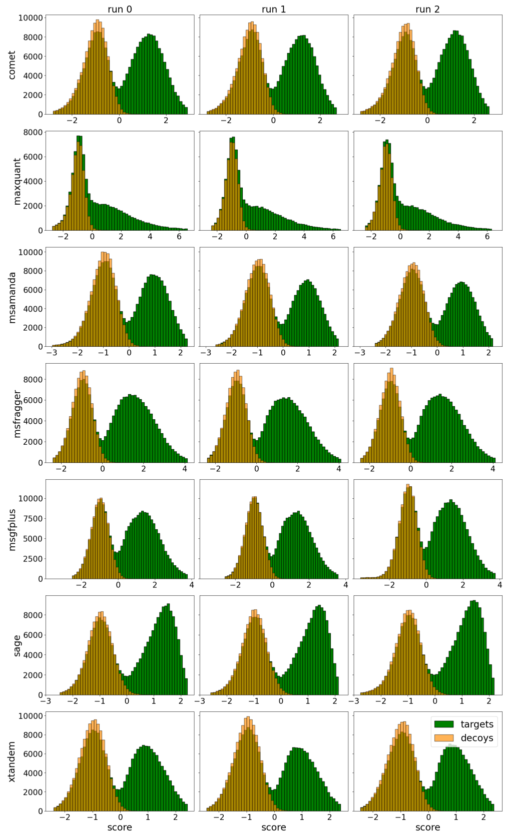

(d) Oktoberfest

Figure S19: Target-decoy distribution, timsTOF dataset, Proteome

##### 1.4.3 ProHap

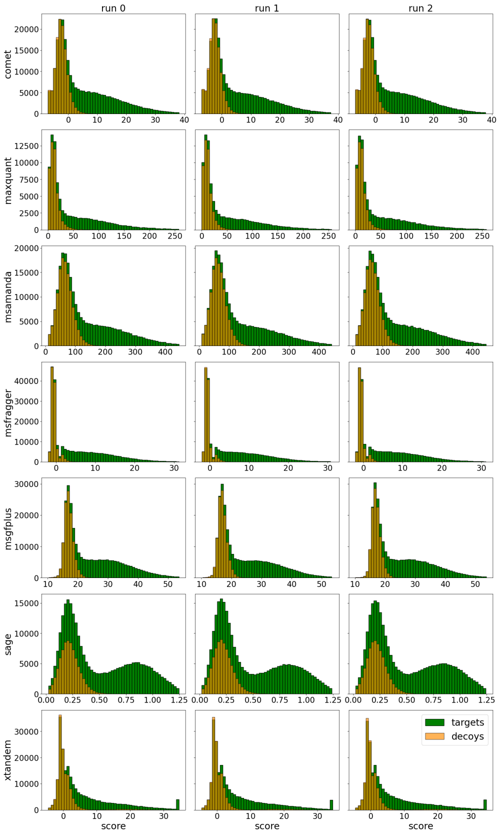

(a) raw search results

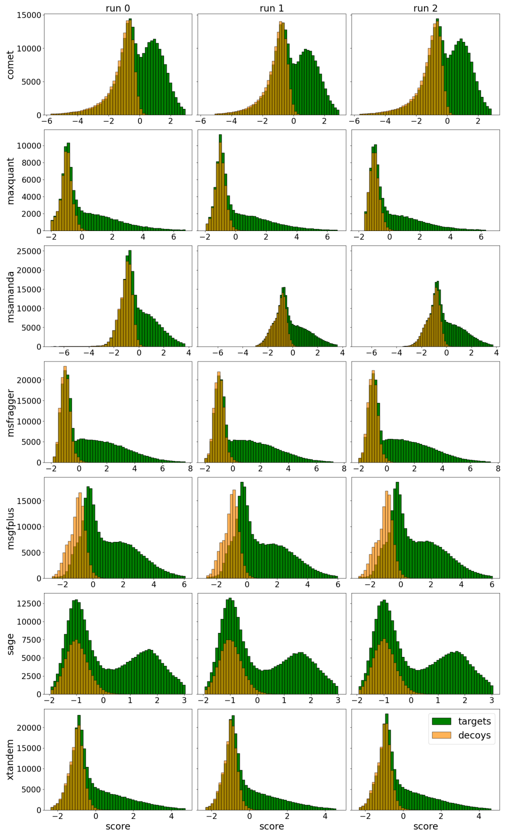

(b) Percolator

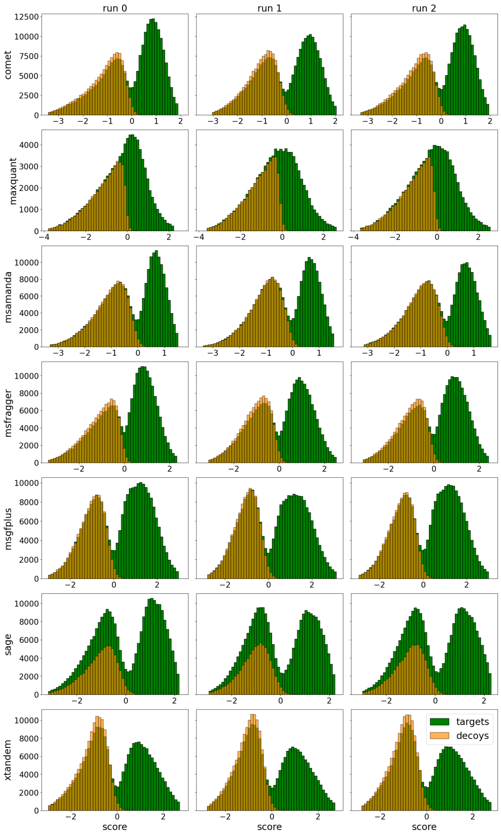

(c) MS2Rescore

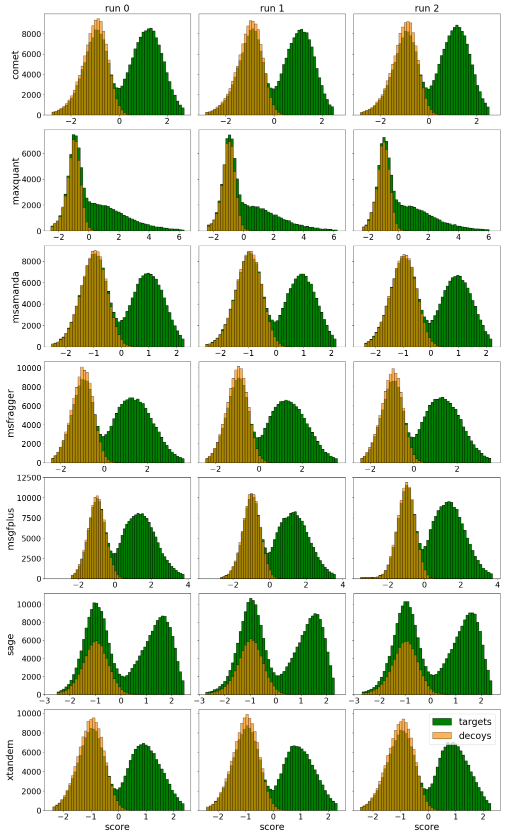

(d) Oktoberfest

Figure S20: Target-decoy distribution, timsTOF dataset, ProHap

##### 1.4.4 ProHap - clear target-decoy-separation

The plots in this section show only the MS-GF+ results of the dataset using the ProHap database, where the target-decoy separation performed too well using all available features to estimate the FDR correctly. Compare the main article for more information.

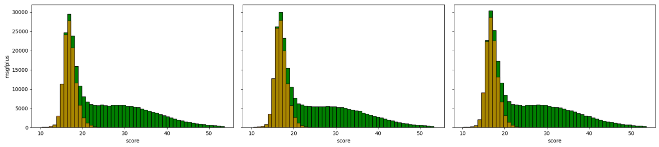

(a) raw search results

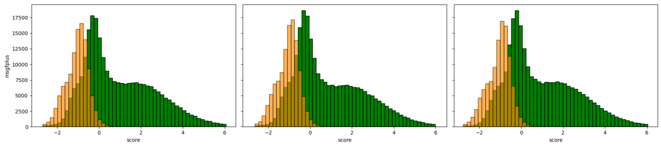

(b) Percolator

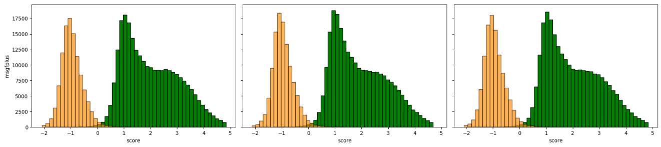

(c) MS2Rescore

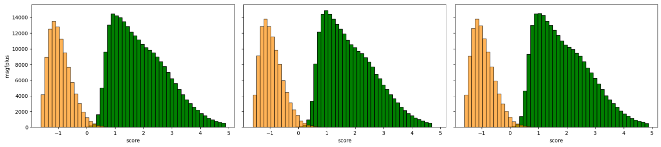

(d) Oktoberfest

Figure S21: Target-decoy distribution with clear separation, timsTOF dataset, ProHap

### 2 Identified PSMs as a function of q-value thresholds

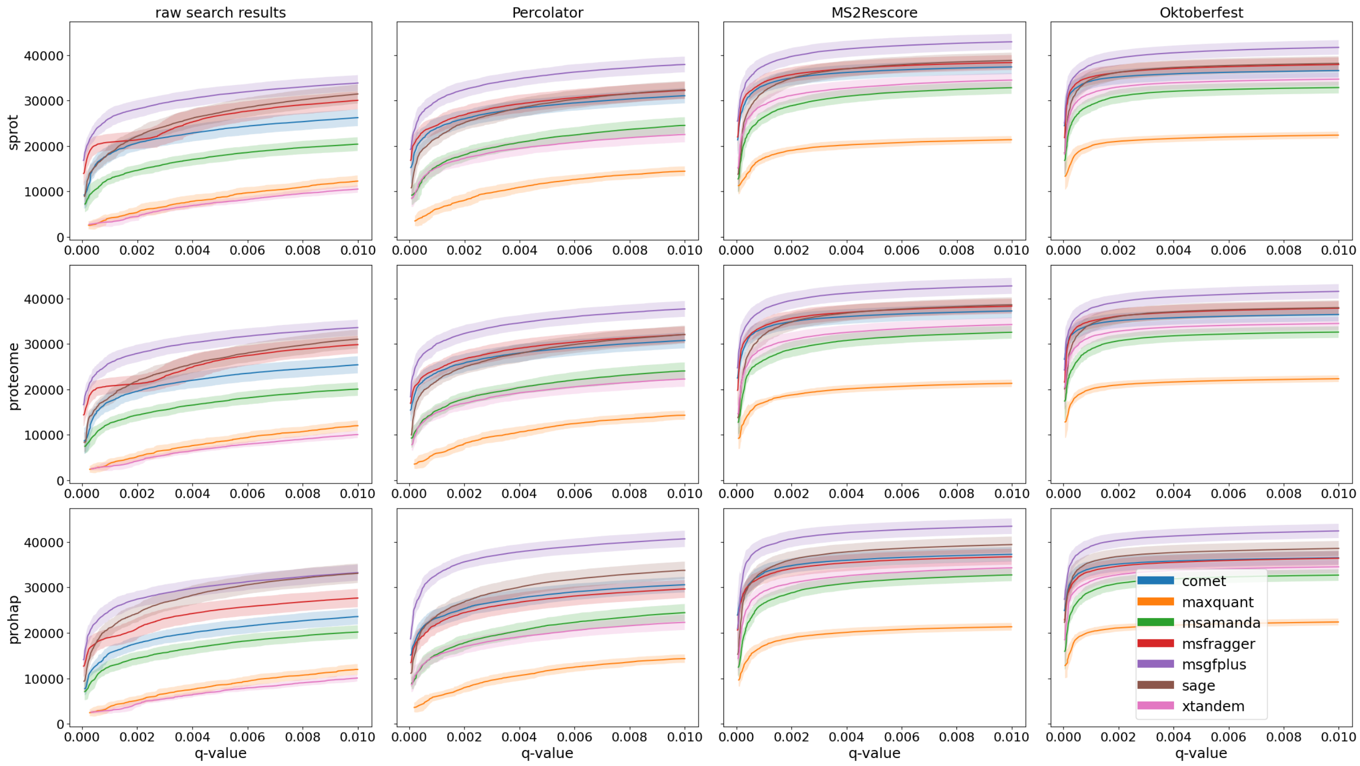

Figure S22: PSMs pseudo ROC for the Cancer Array Dataset

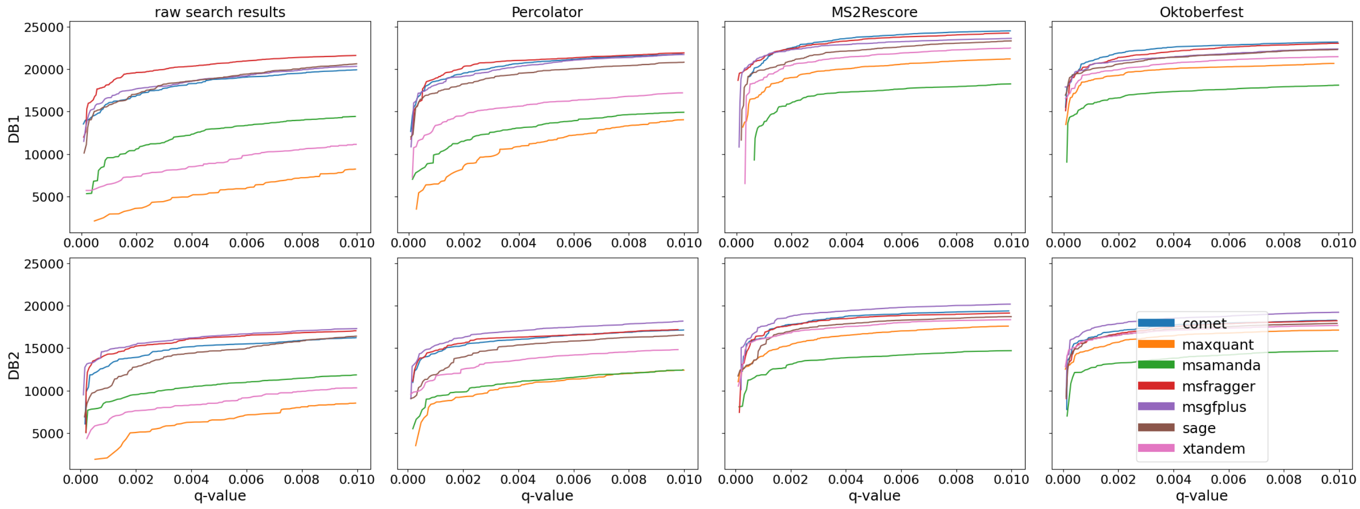

Figure S23: PSMs pseudo ROC for the CAMPI Dataset

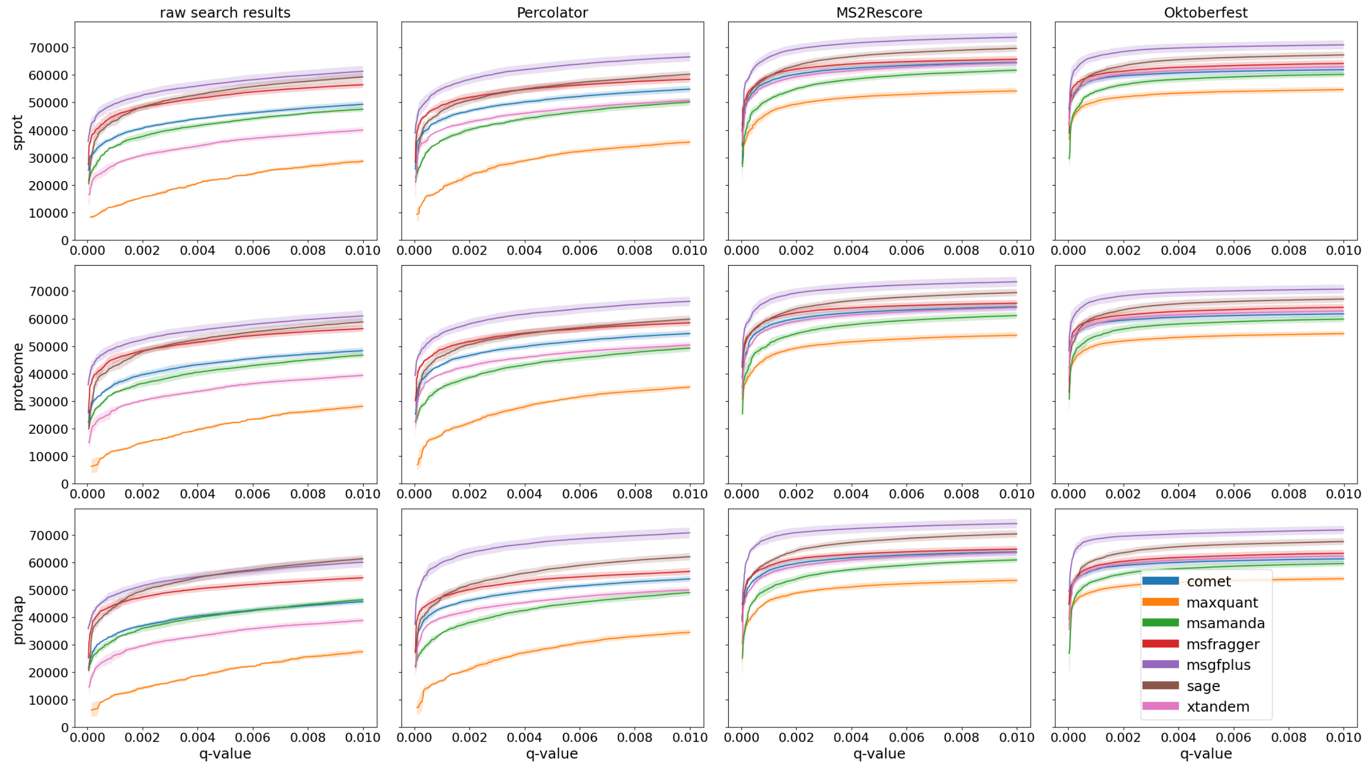

Figure S24: PSMs pseudo ROC for the Orbitrap dataset

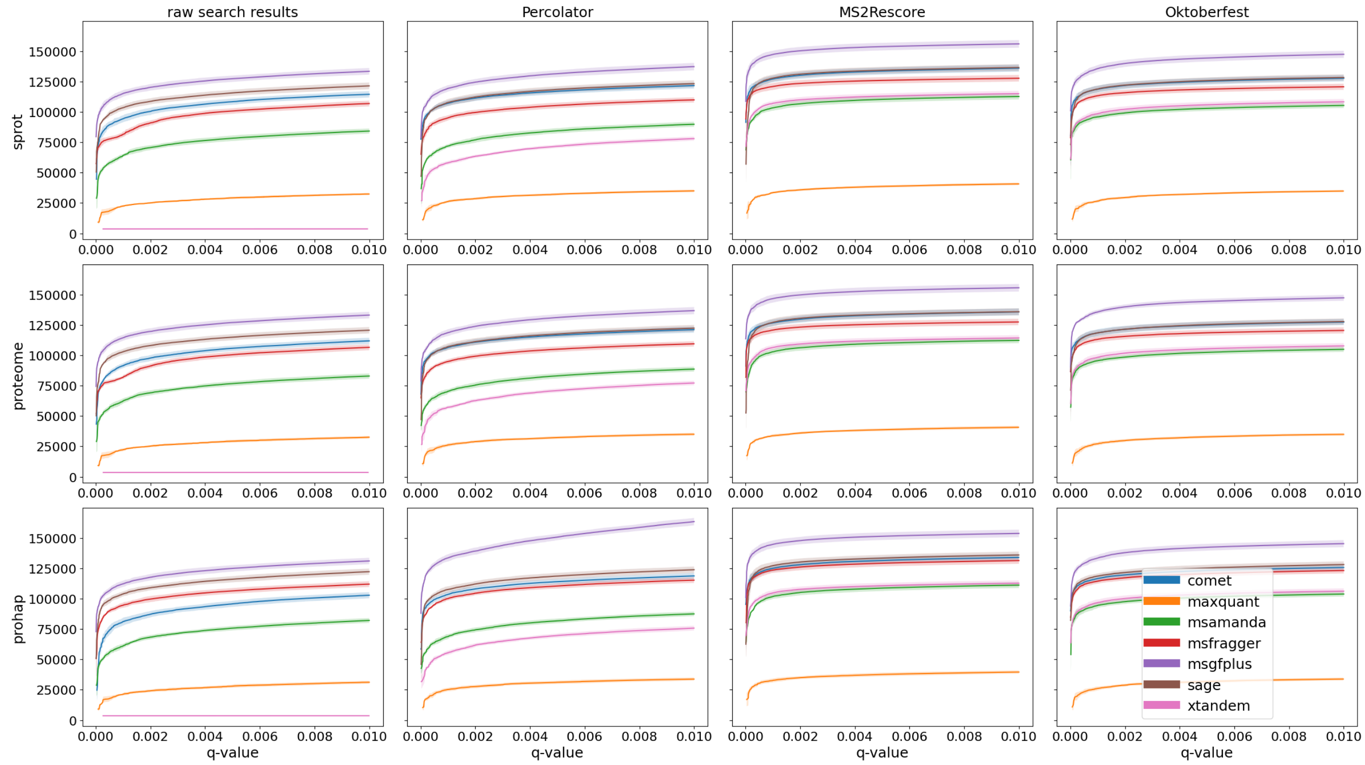

Figure S25: PSMs pseudo ROC for the timsTOF dataset

### 3 Number of identifications per search engine and rescoring method

#### 3.1 Cancer Array Dataset

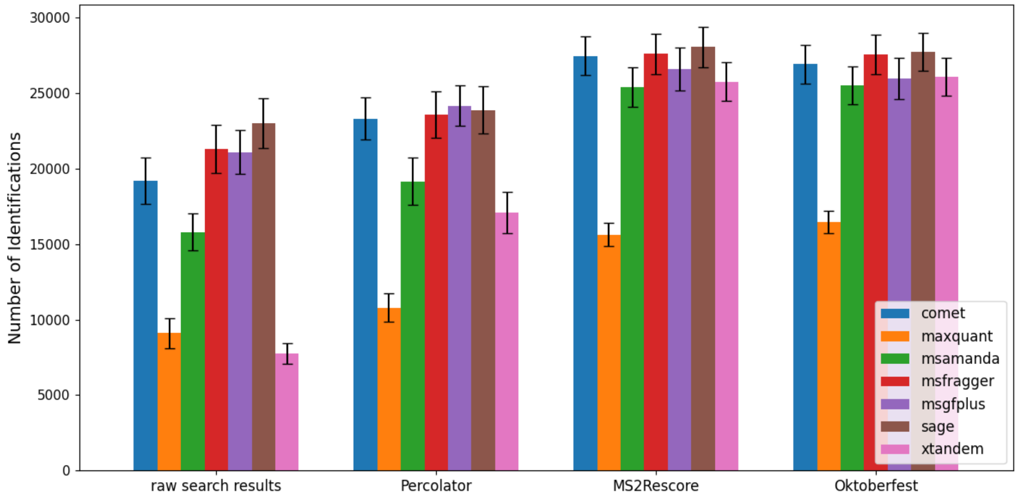

Figure S26: Numbers of peptidoforms for the Cancer Array Dataset and Swiss-Prot database

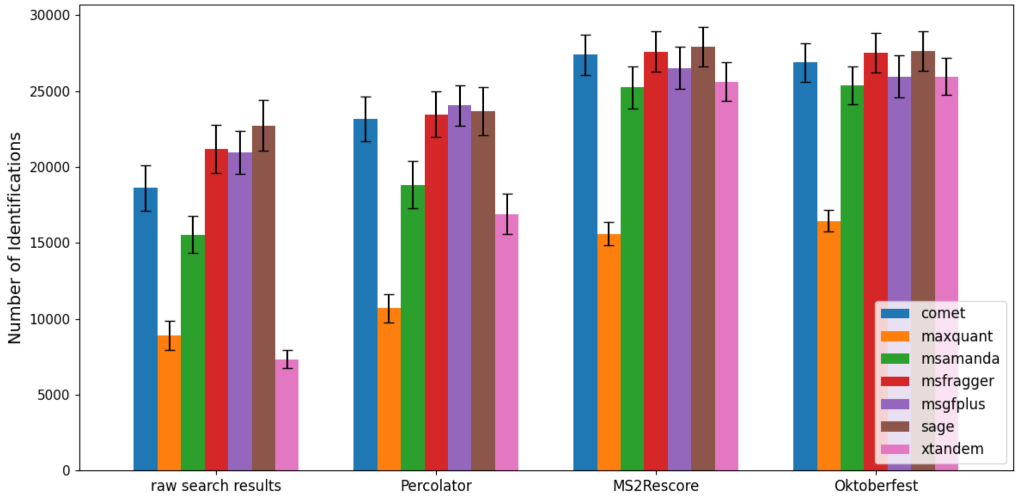

Figure S27: Numbers of peptidoforms for the Cancer Array Dataset and Proteome database

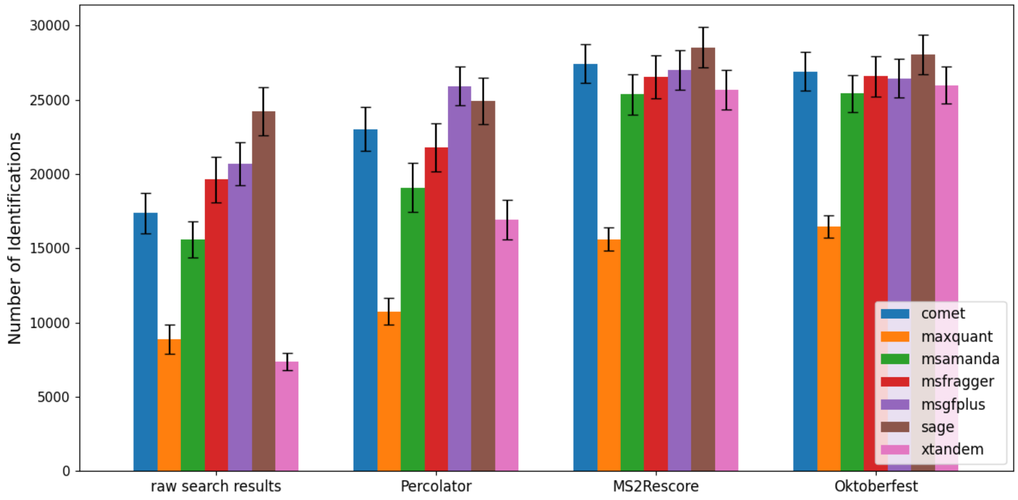

Figure S28: Numbers of peptidoforms for the Cancer Array Dataset and ProHap database

#### 3.2 CAMPI Dataset

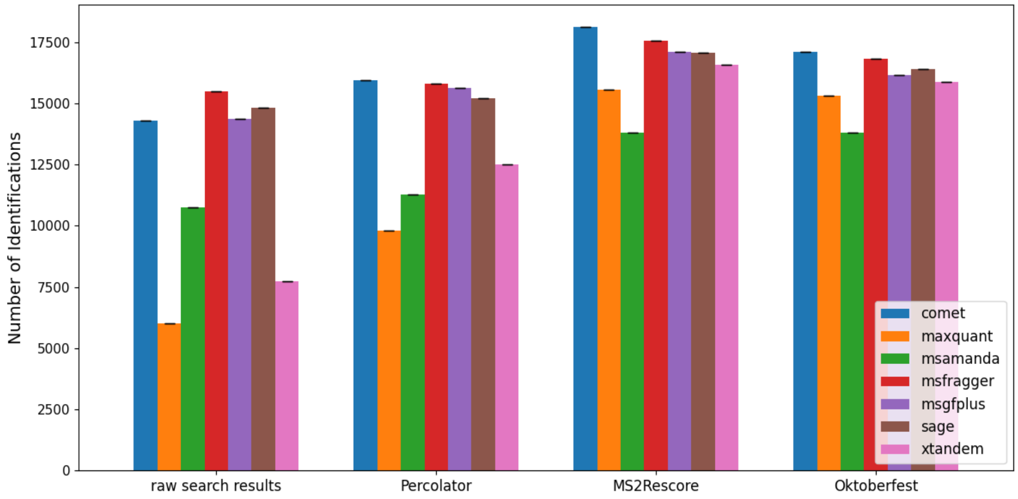

Figure S29: Numbers of peptidoforms for the CAMPI Dataset and DB1

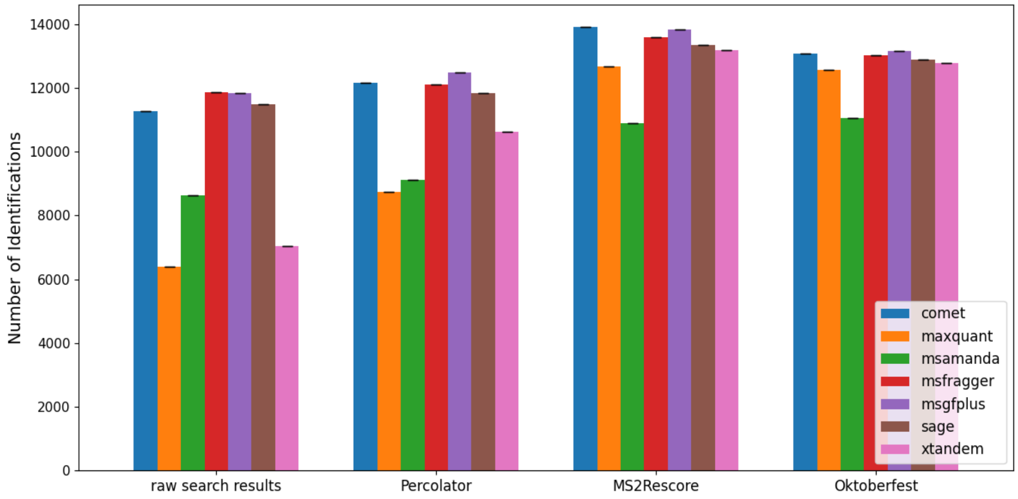

Figure S30: Numbers of peptidoforms for the CAMPI Dataset and DB2

#### 3.3 Orbitrap Dataset

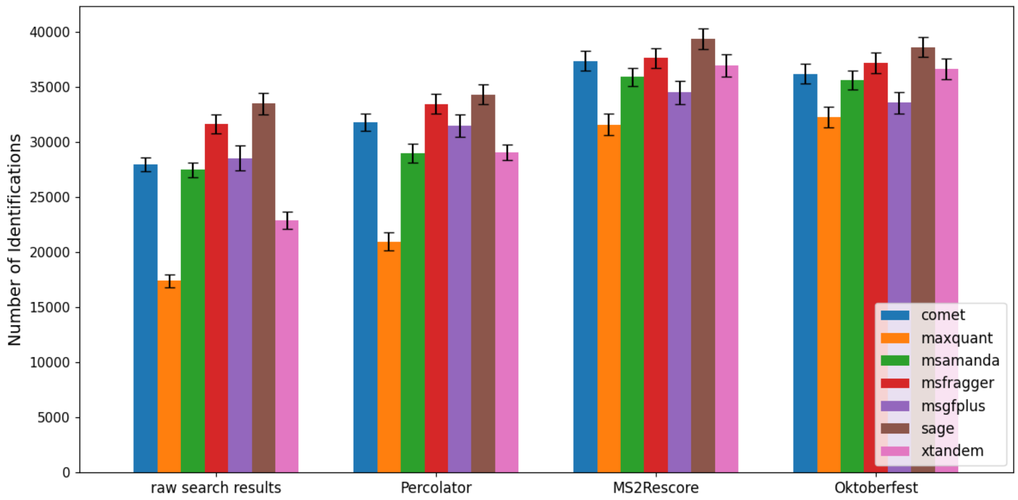

Figure S31: Numbers of peptidoforms for the Orbitrap dataset and Swiss-Prot database

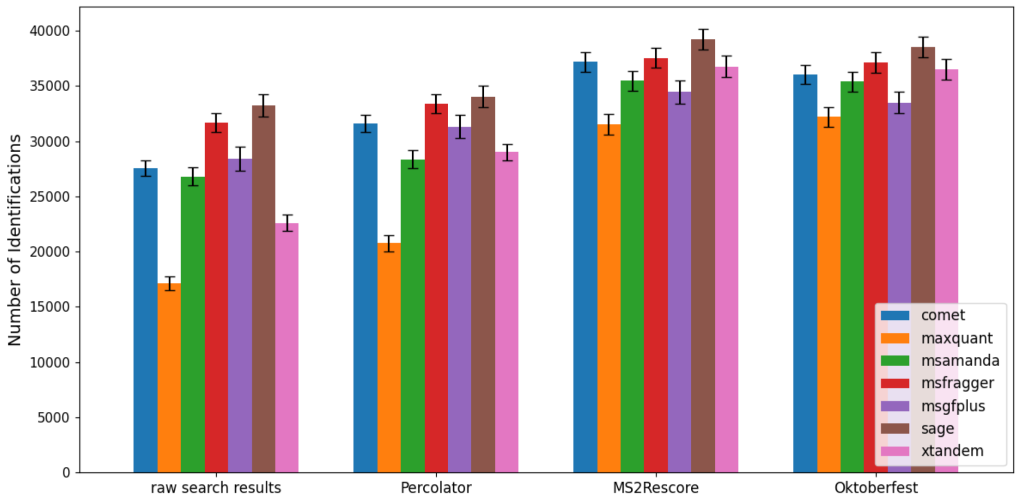

Figure S32: Numbers of peptidoforms for the Orbitrap dataset and Proteome database

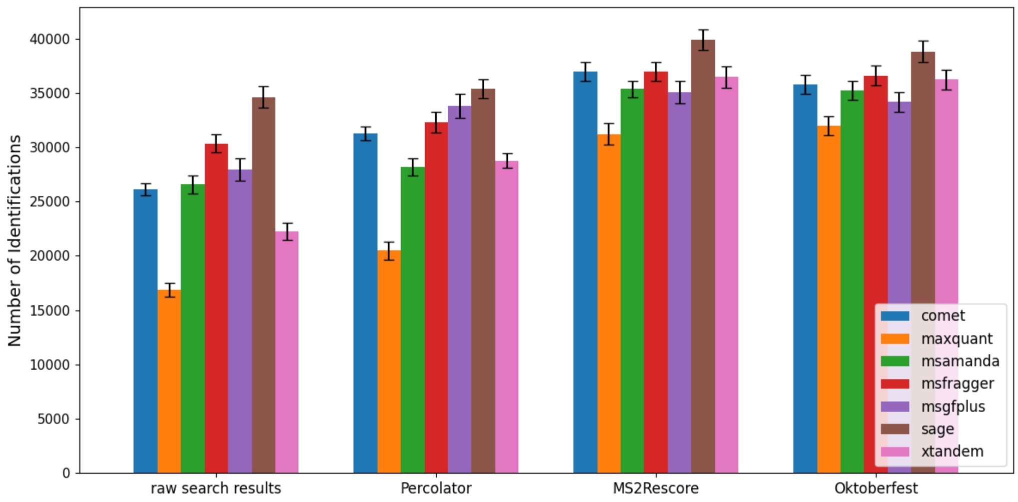

Figure S33: Numbers of peptidoforms for the Orbitrap dataset and ProHap database

#### 3.4 timsTOF Dataset

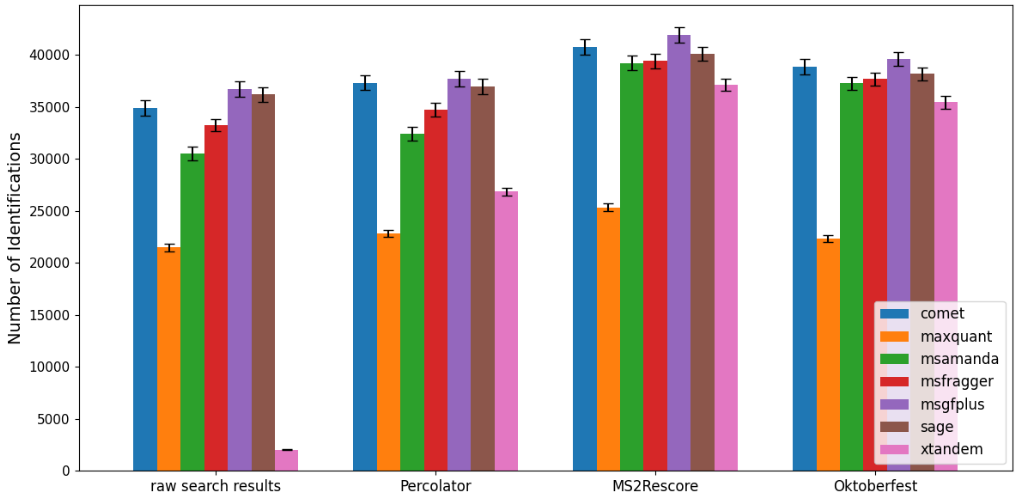

Figure S34: Numbers of peptidoforms for the timsTOF dataset and Swiss-Prot database

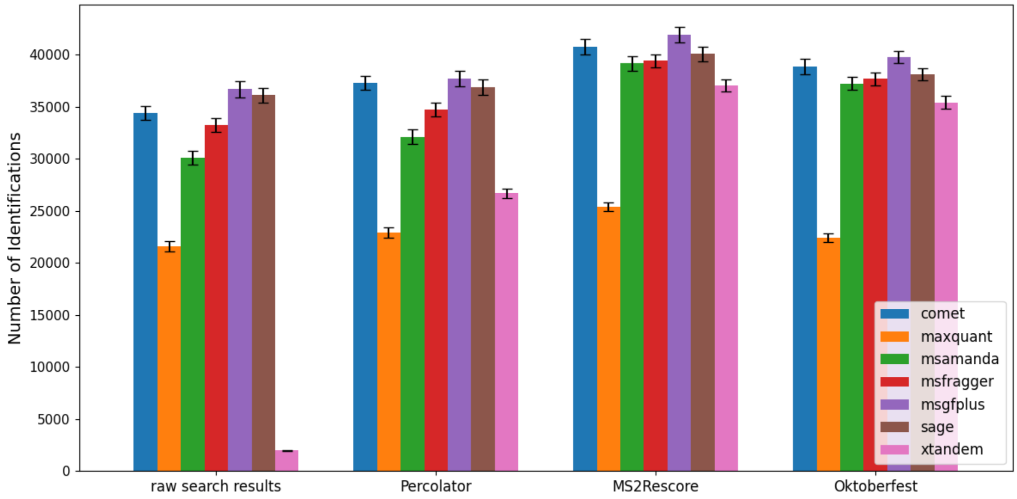

Figure S35: Numbers of peptidoforms for the timsTOF dataset and Proteome database

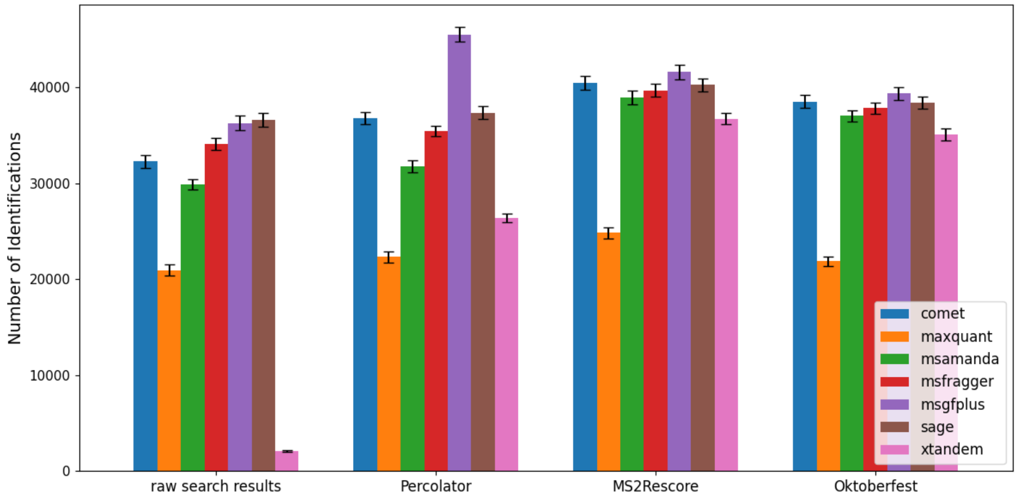

Figure S36: Numbers of peptidoforms for the timsTOF dataset and ProHap database

### 4 Overlap of databases per search engine

Only groups comprising at least 1% of the number of peptidoform identifications in the largest group are shown.

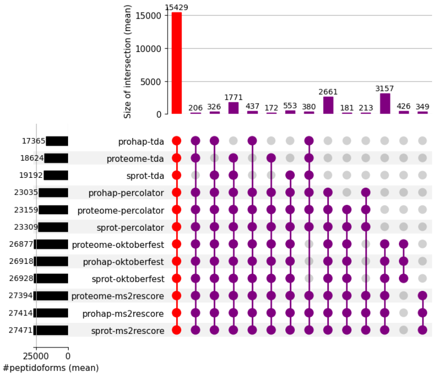

(a) Comet

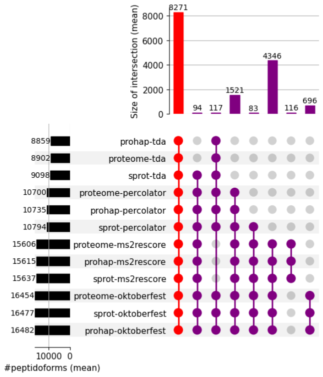

(b) MaxQuant

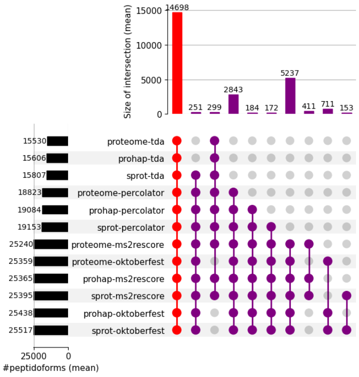

(c) MS Amanda

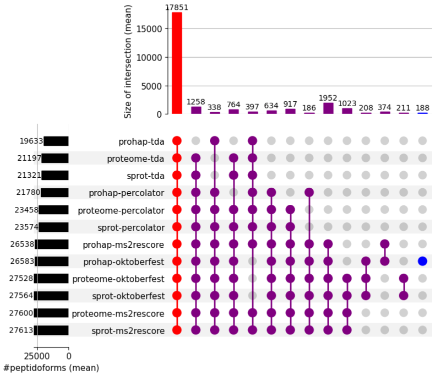

(d) MSFragger

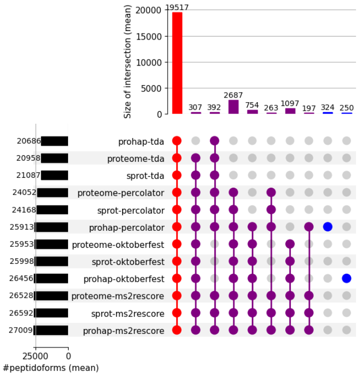

(e) MS-GF+

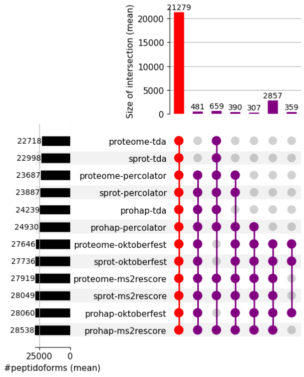

(f) Sage

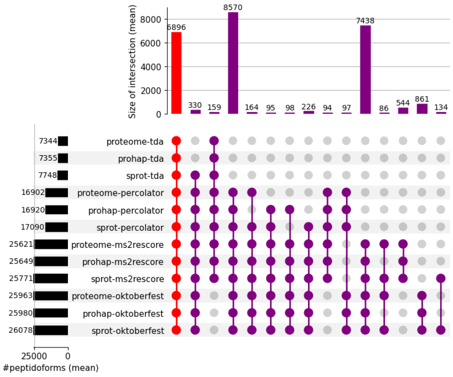

(g) X!Tandem

Figure S37: Overlap per search engine, database, and rescoring, Cancer Array Dataset

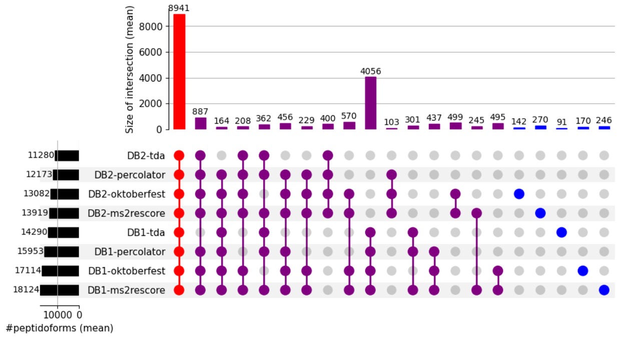

(a) Comet

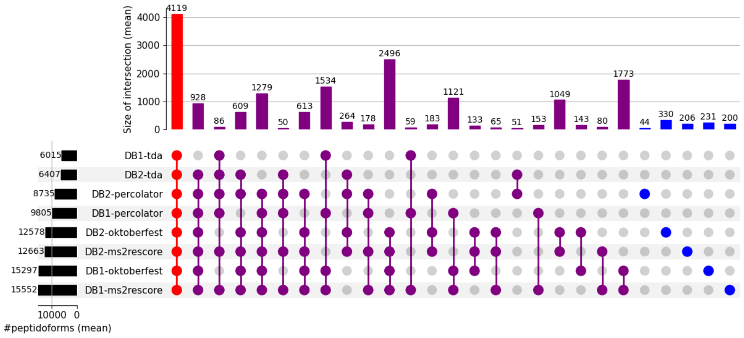

(b) MaxQuant

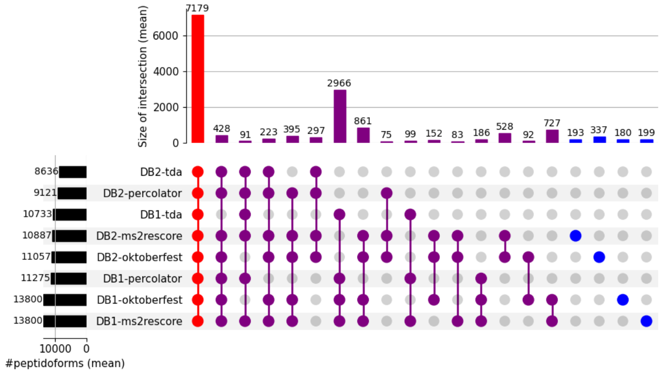

(c) MS Amanda

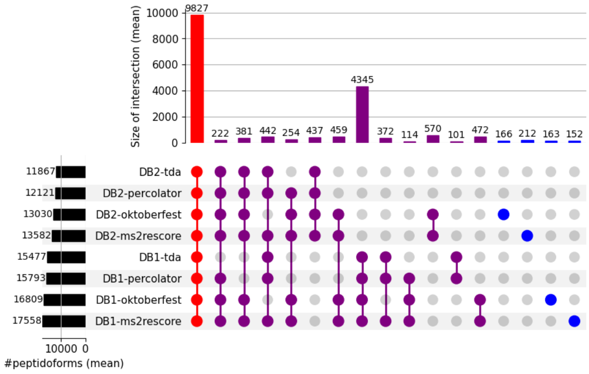

(d) MSFragger

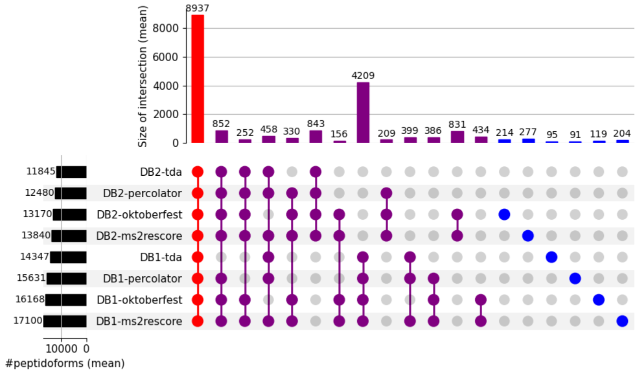

(e) MS-GF+

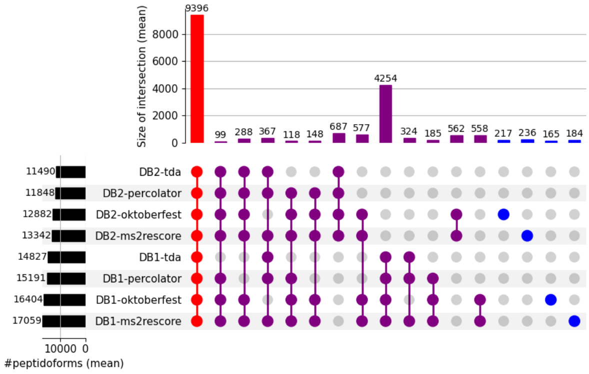

(f) Sage

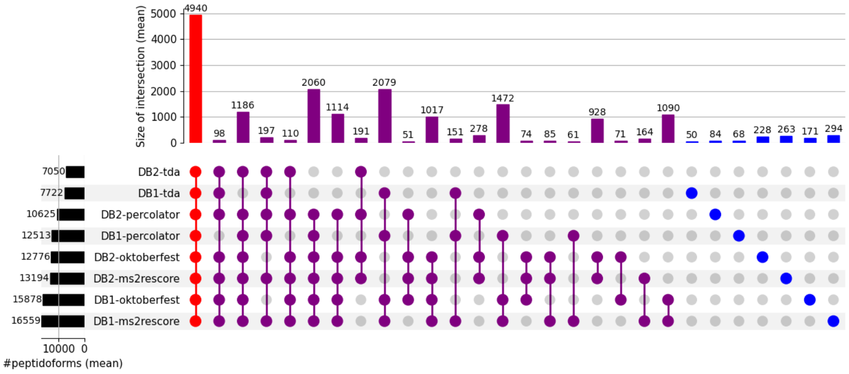

(g) X!Tandem

Figure S38: Overlap per search engine, database, and rescoring, CAMPI Dataset

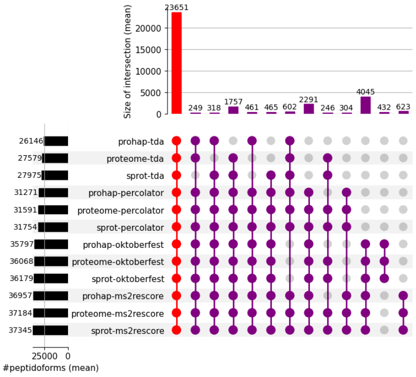

(a) Comet

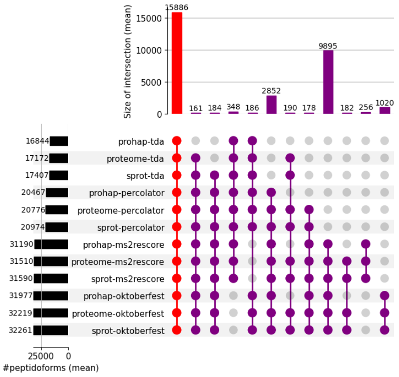

(b) MaxQuant

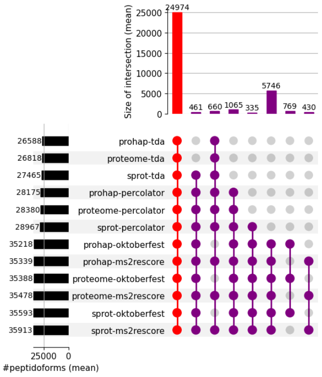

(c) MS Amanda

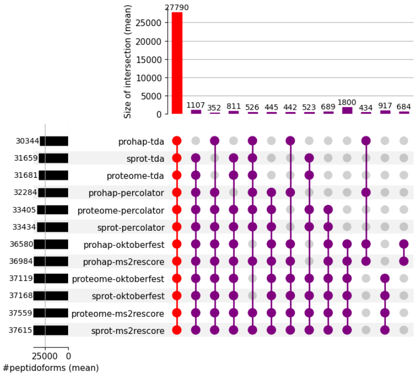

(d) MSFragger

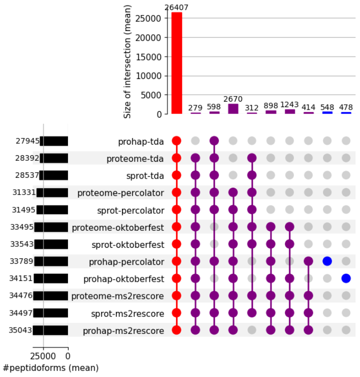

(e) MS-GF+

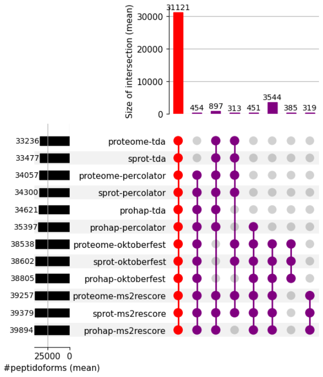

(f) Sage

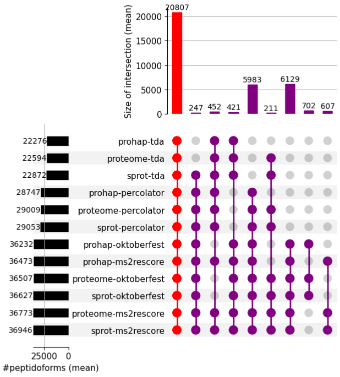

(g) X!Tandem

Figure S39: Overlap per search engine, database, and rescoring, Orbitrap dataset

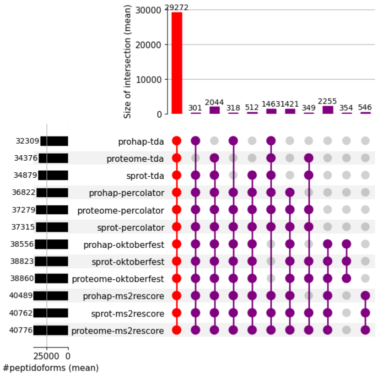

(a) Comet

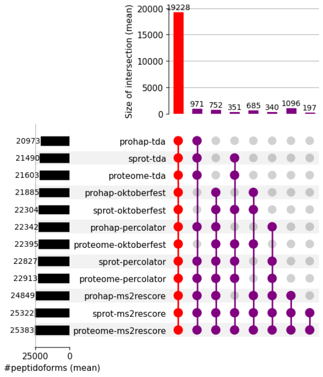

(b) MaxQuant

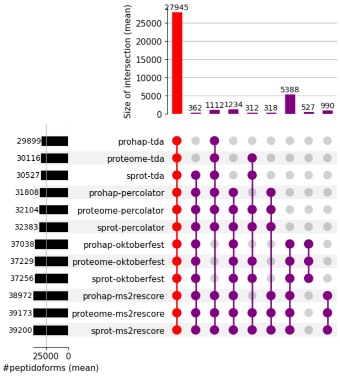

(c) MS Amanda

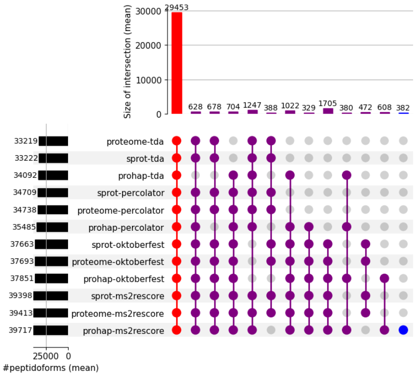

(d) MSFragger

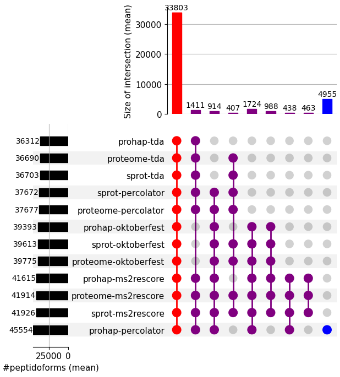

(e) MS-GF+

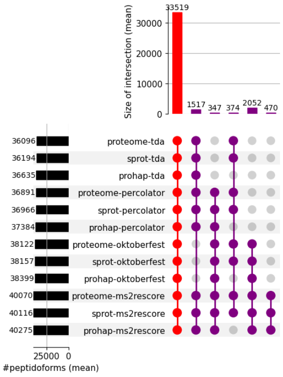

(f) Sage

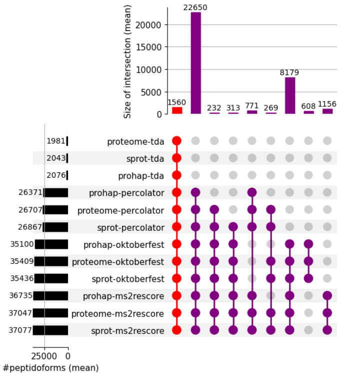

(g) X!Tandem

Figure S40: Overlap per search engine, database, and rescoring, timsTOF dataset

### 5 Overlap of all search engines per rescoring method

For the groups representing the overlap of all, all but one and the identifications unique to one search engine all results are shown, while for all other groups only groups comprising at least 1% of the number of identifications in the largest group are shown.

#### 5.1 Cancer Array Dataset

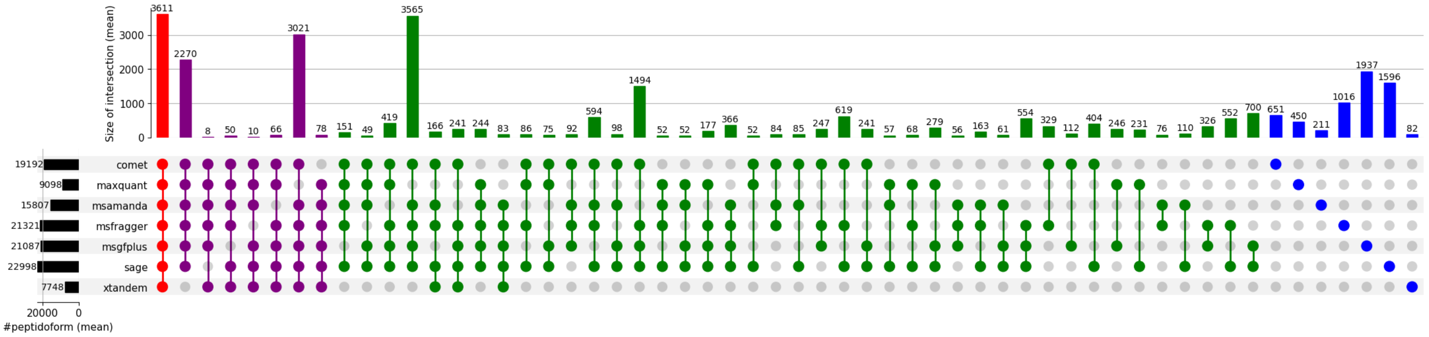

(a) raw search results

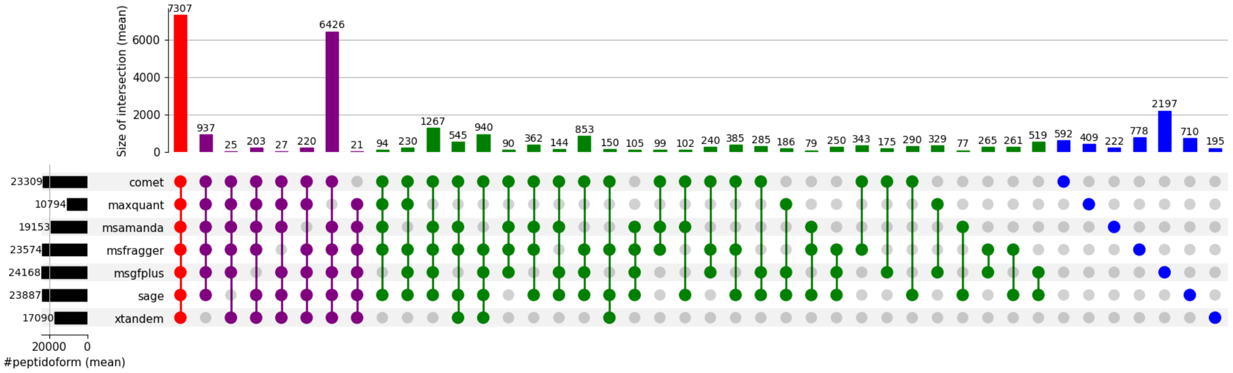

(b) Percolator

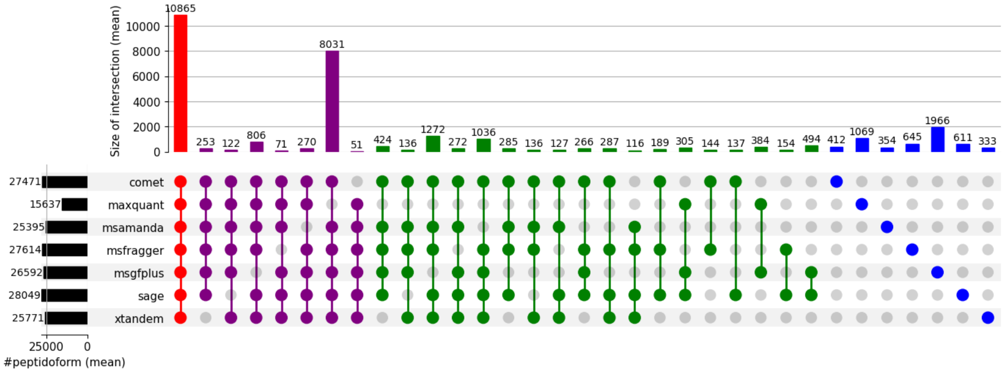

(c) MS2Rescore

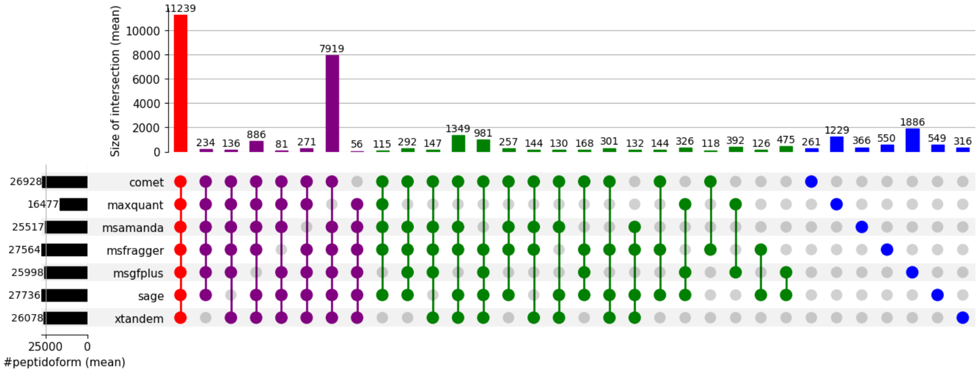

(d) Oktoberfest

Figure S41: Overlap of search engines per rescoring method, Cancer Array Dataset, Swiss-Prot

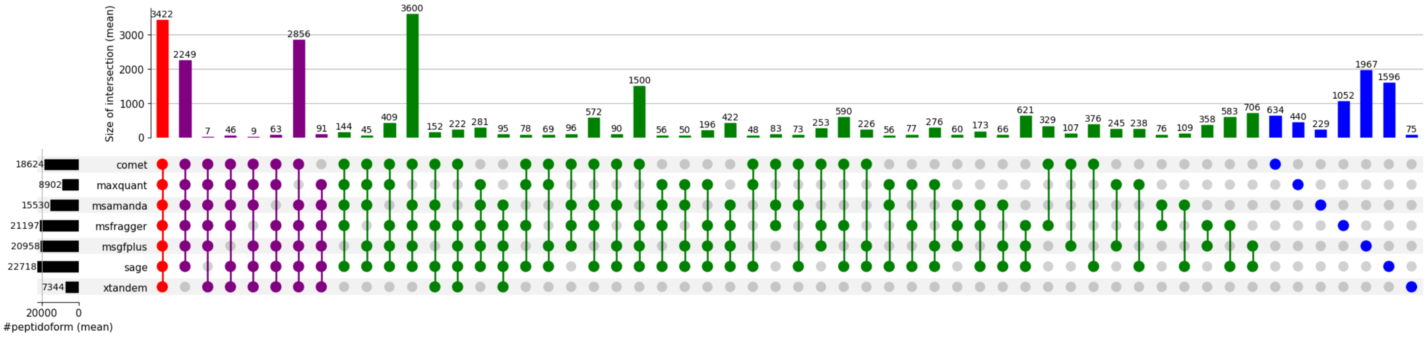

(a) raw search results

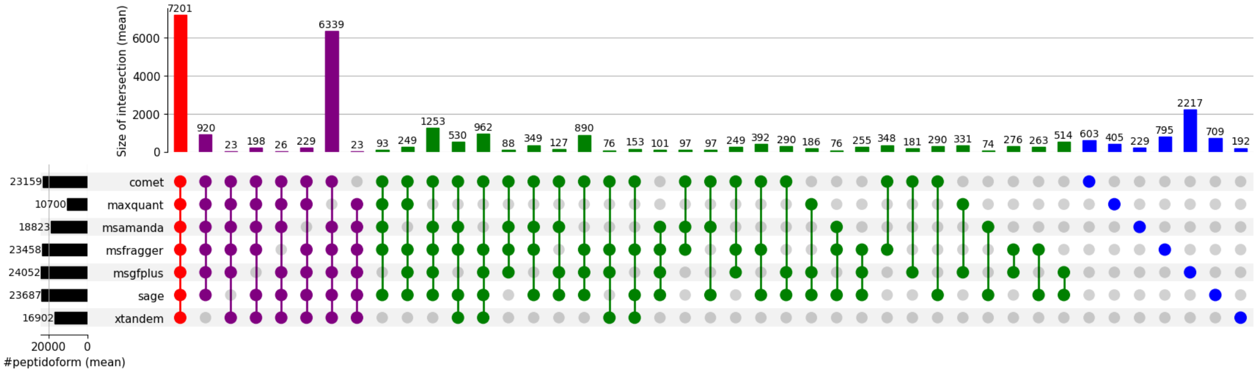

(b) Percolator

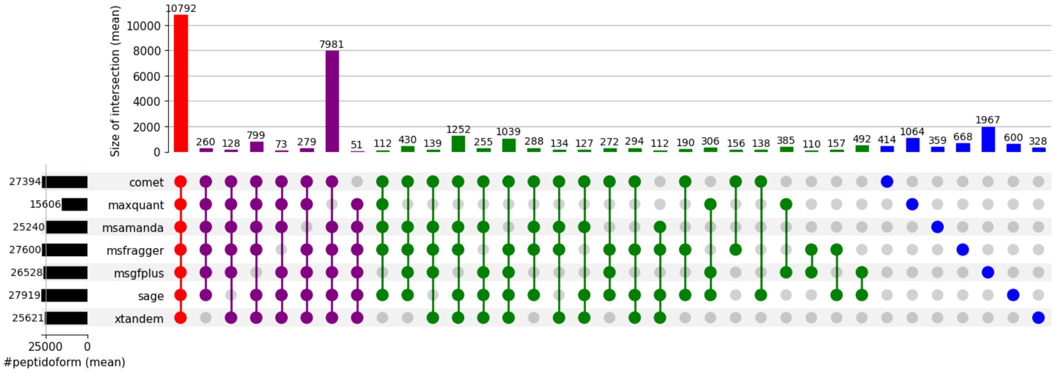

(c) MS2Rescore

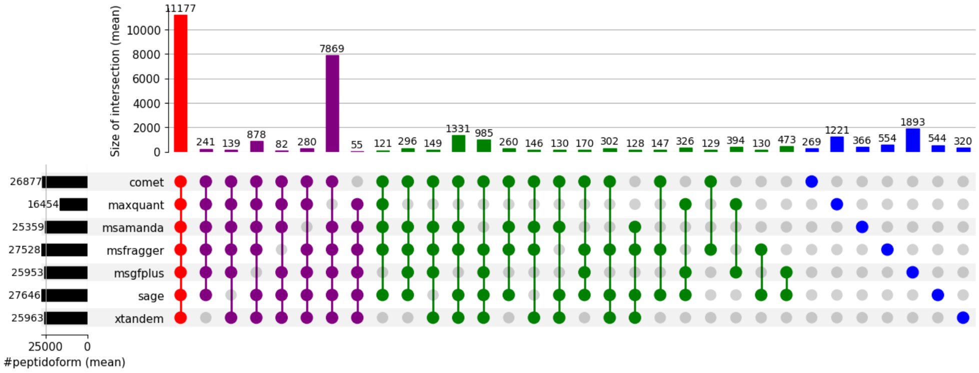

(d) Oktoberfest

Figure S42: Overlap of search engines per rescoring method, Cancer Array Dataset, Proteome

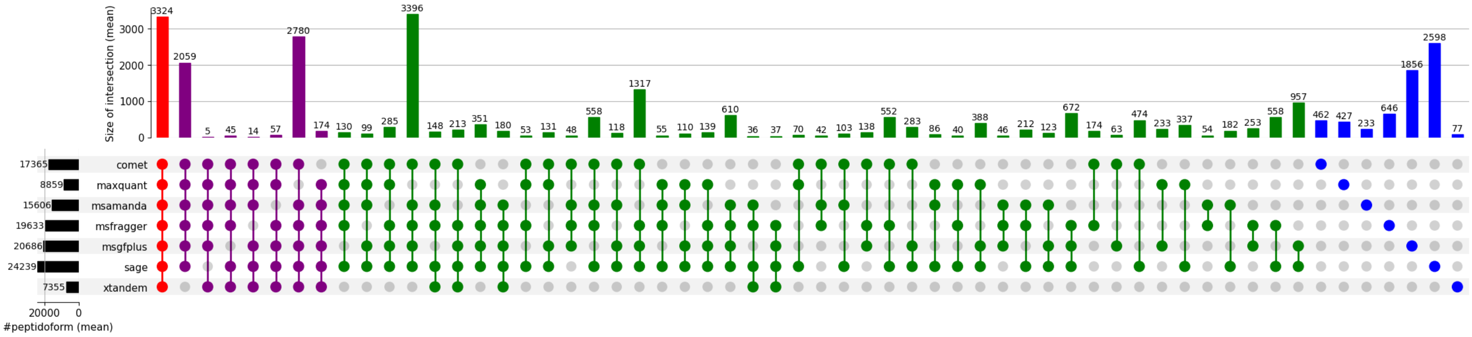

(a) raw search results

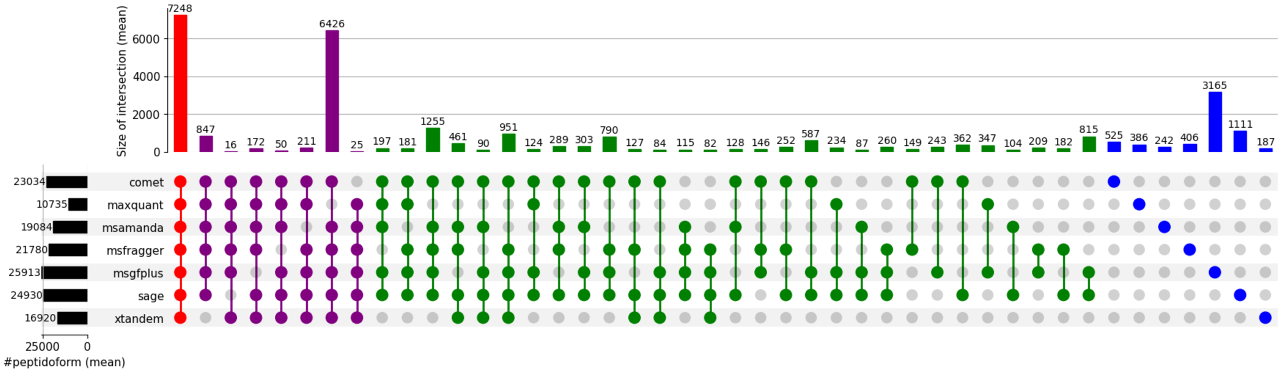

(b) Percolator

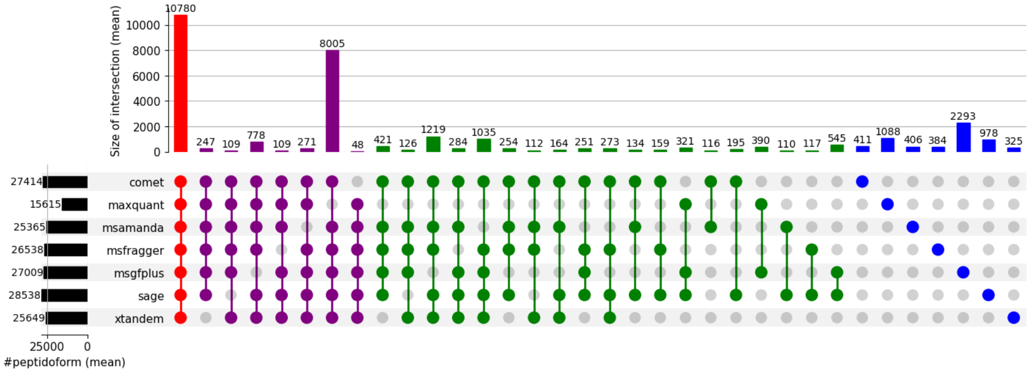

(c) MS2Rescore

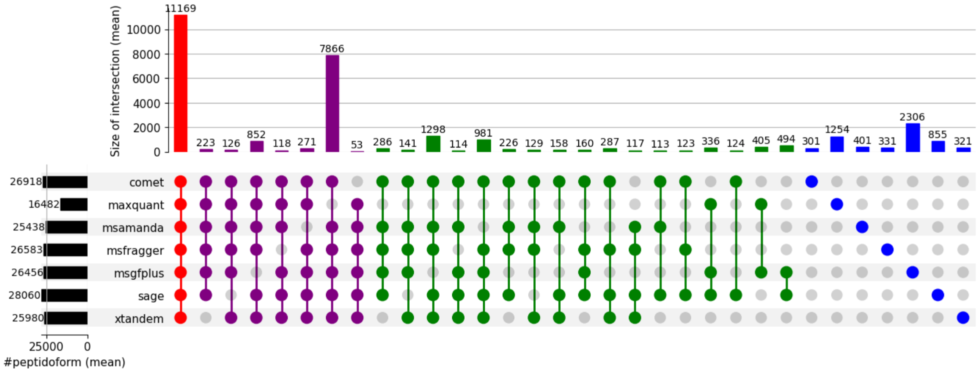

(d) Oktoberfest

Figure S43: Overlap of search engines per rescoring method, Cancer Array Dataset, ProHap

#### 5.2 CAMPI Dataset

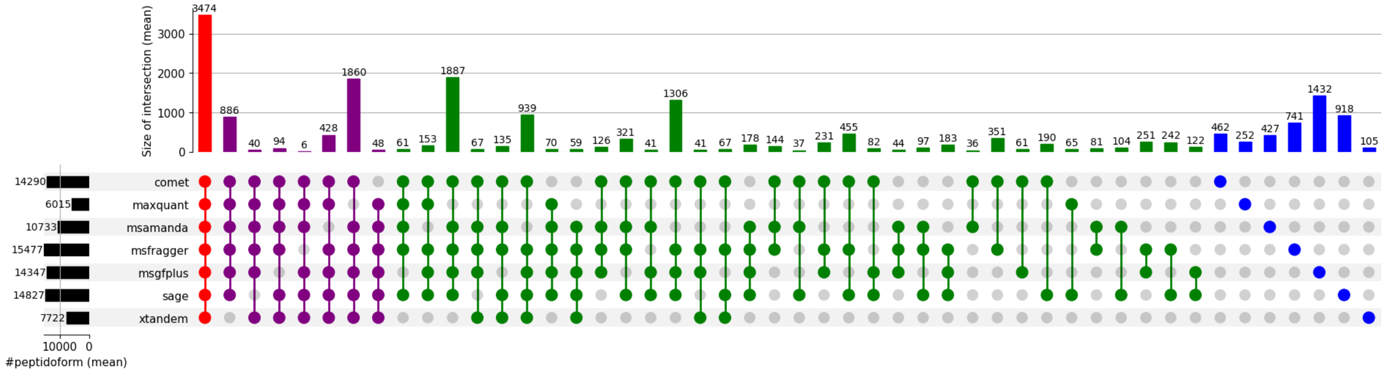

(a) raw search results

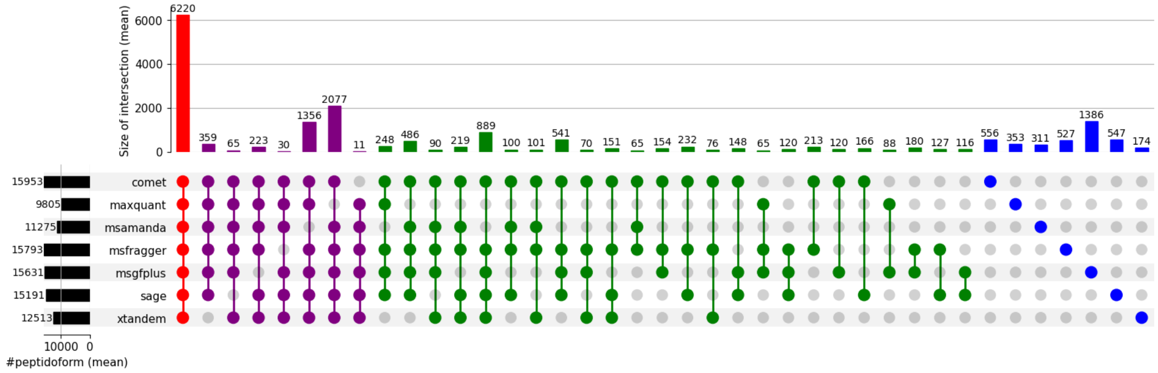

(b) Percolator

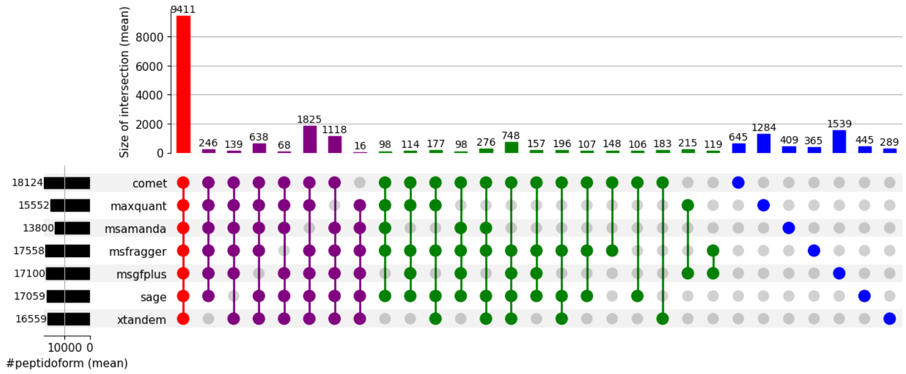

(c) MS2Rescore

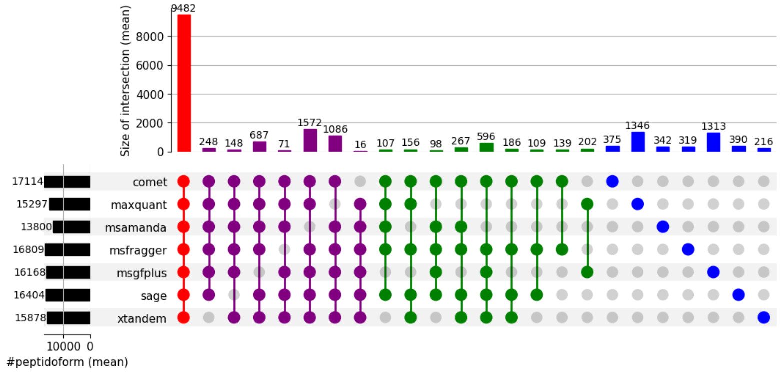

(d) Oktoberfest

Figure S44: Overlap of search engines per rescoring method, CAMPI Dataset, DB1

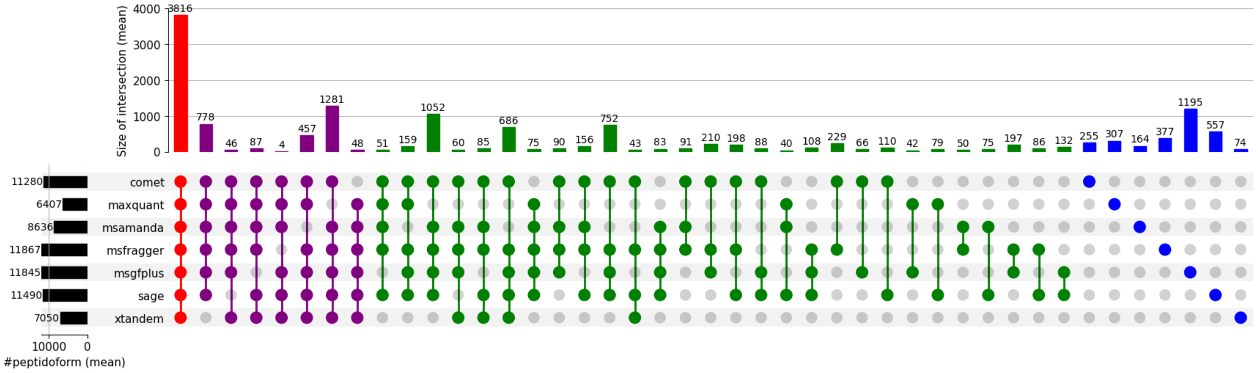

(a) raw search results

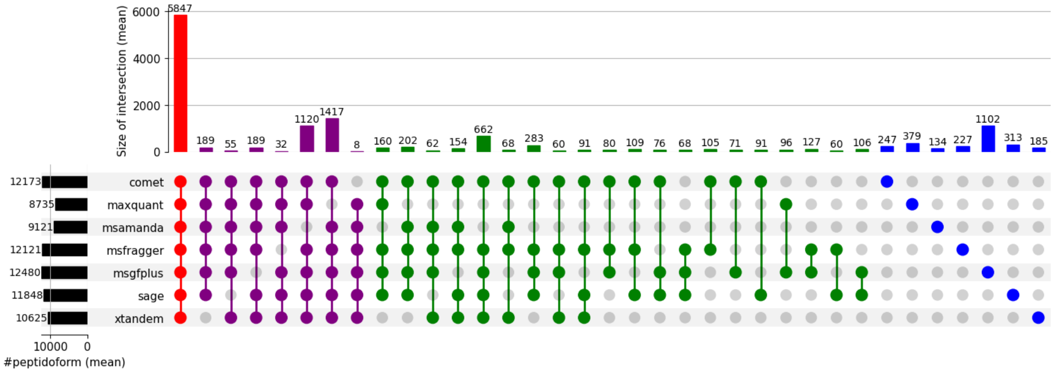

(b) Percolator

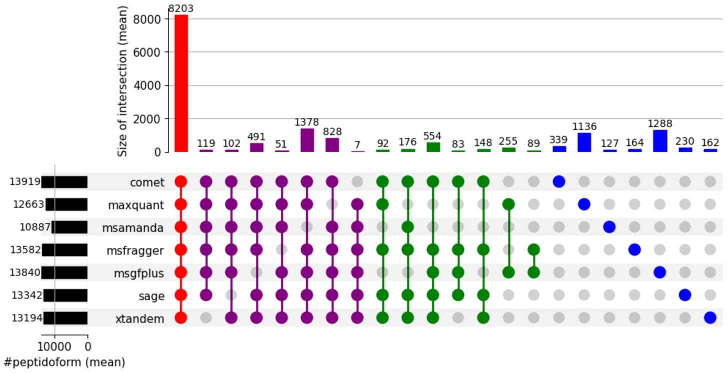

(c) MS2Rescore

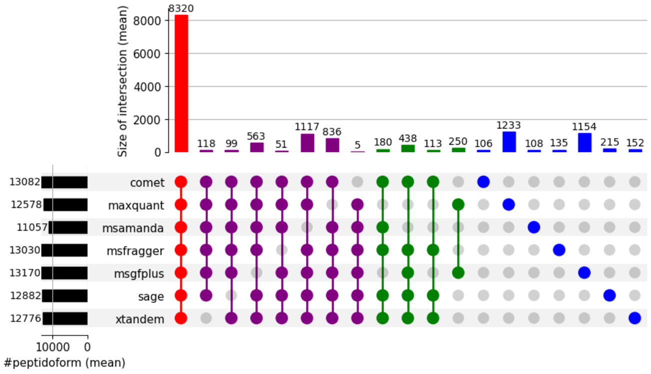

(d) Oktoberfest

Figure S45: Overlap of search engines per rescoring method, CAMPI Dataset, DB2

#### 5.3 Orbitrap Dataset

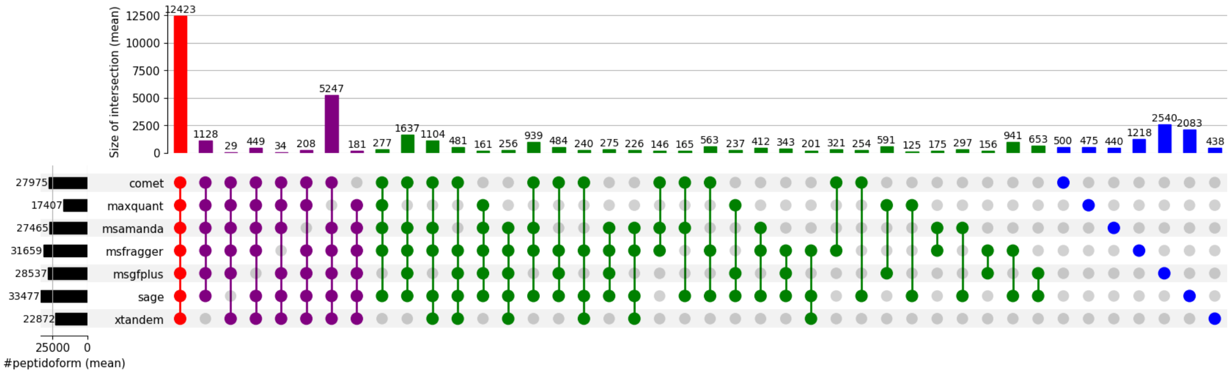

(a) raw search results

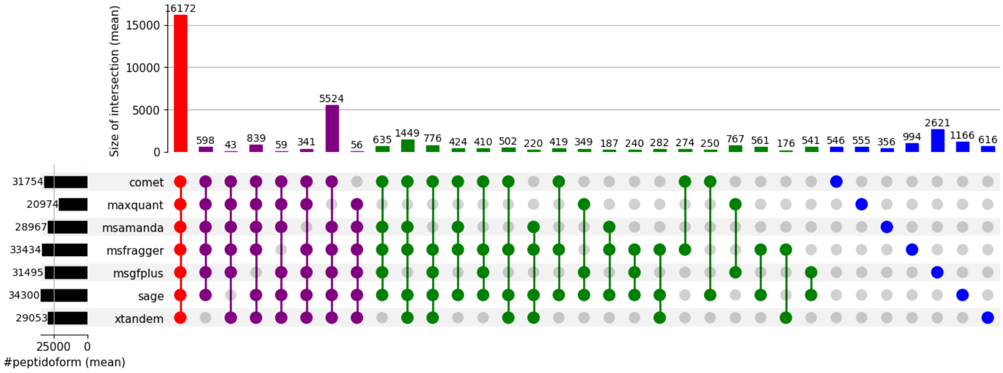

(b) Percolator

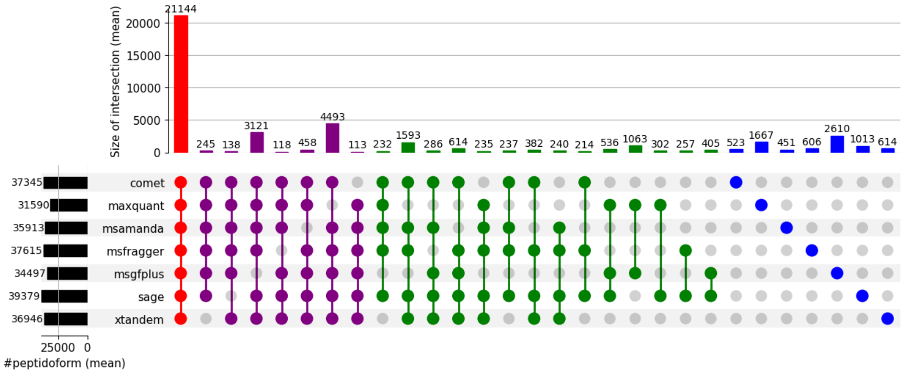

(c) MS2Rescore

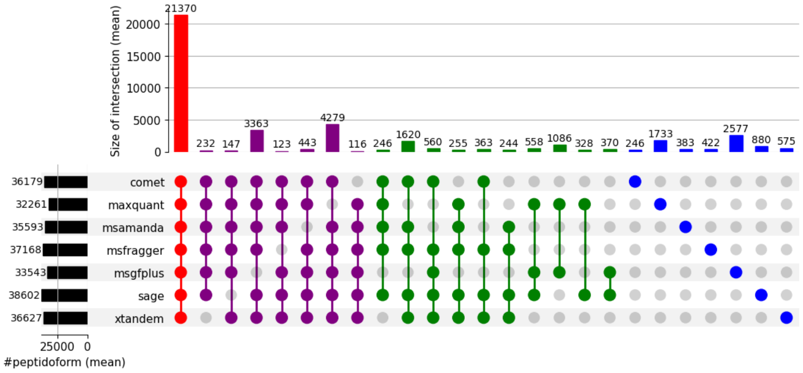

(d) Oktoberfest

Figure S46: Overlap of search engines per rescoring method, Orbitrap dataset, Swiss-Prot

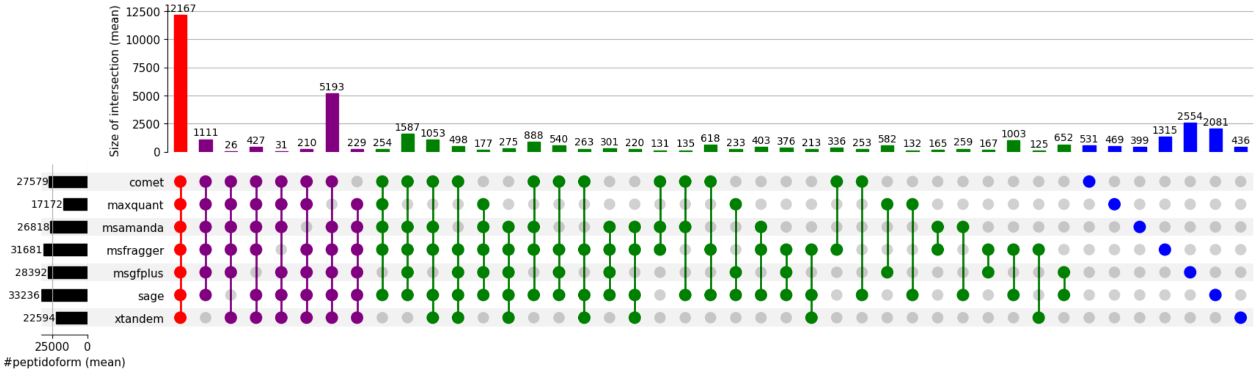

(a) raw search results

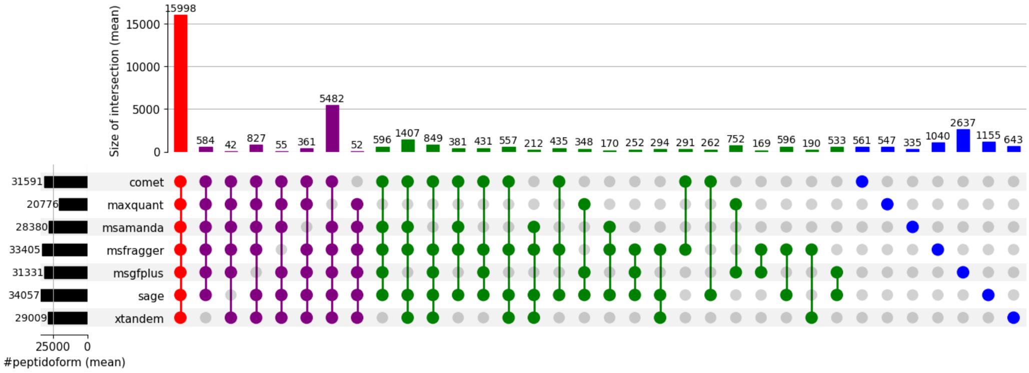

(b) Percolator

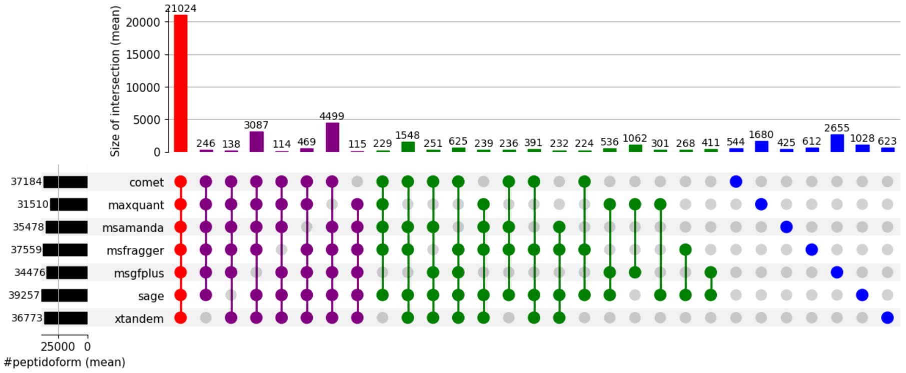

(c) MS2Rescore

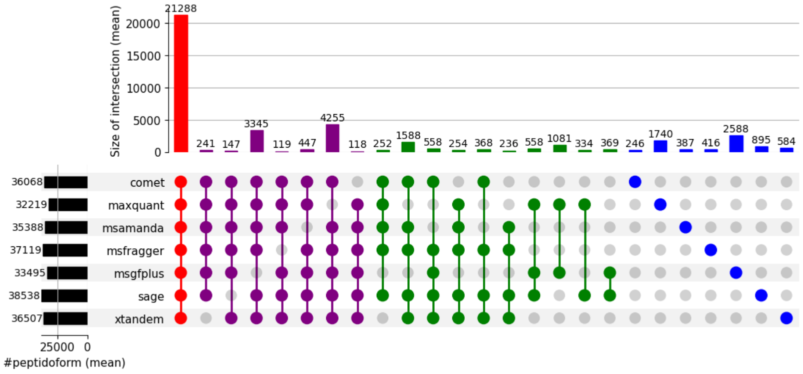

(d) Oktoberfest

Figure S47: Overlap of search engines per rescoring method, Orbitrap dataset, Proteome

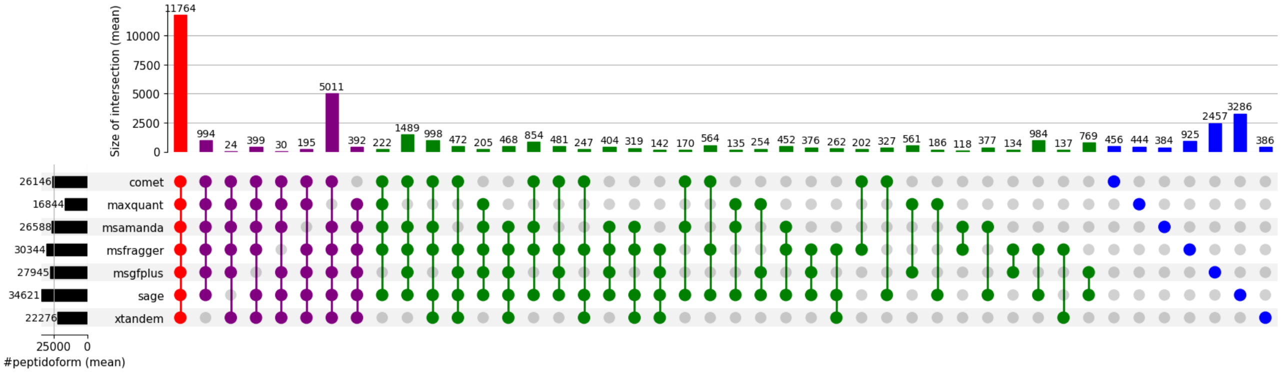

(a) raw search results

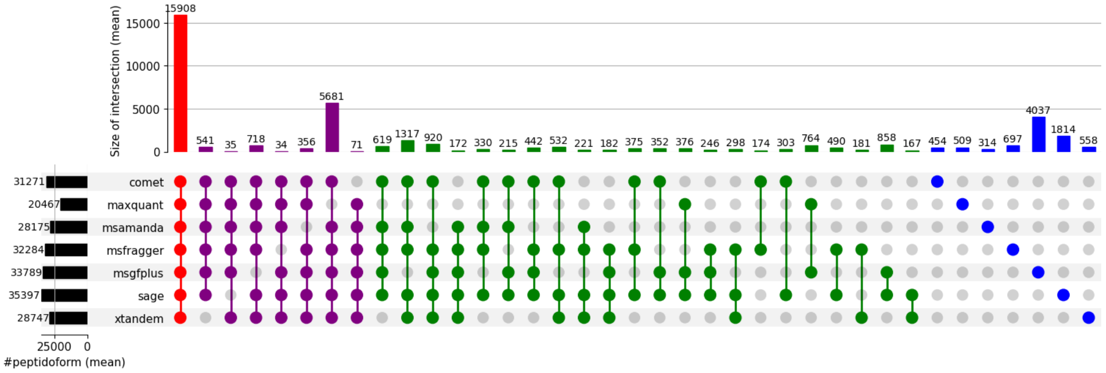

(b) Percolator

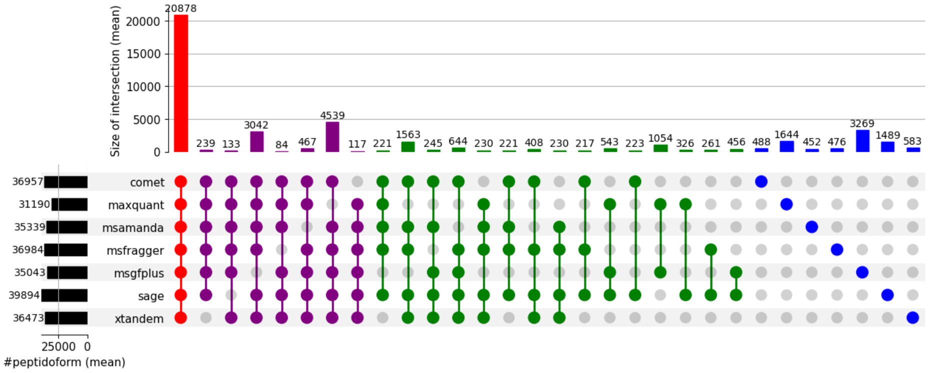

(c) MS2Rescore

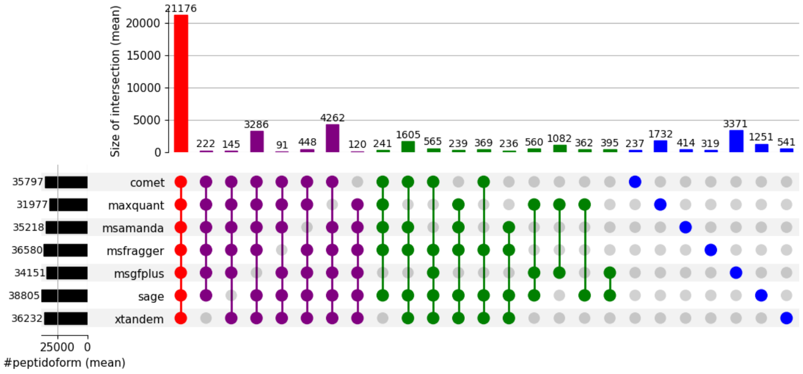

(d) Oktoberfest

Figure S48: Overlap of search engines per rescoring method, Orbitrap dataset, ProHap

#### 5.4 timsTOF Dataset

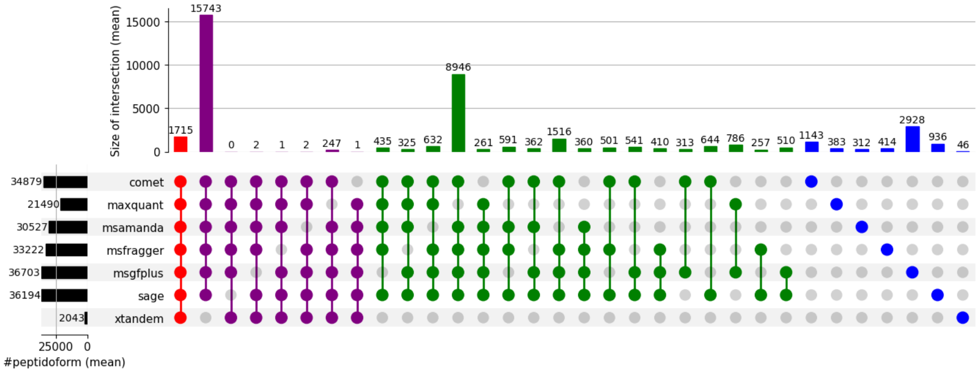

(a) raw search results

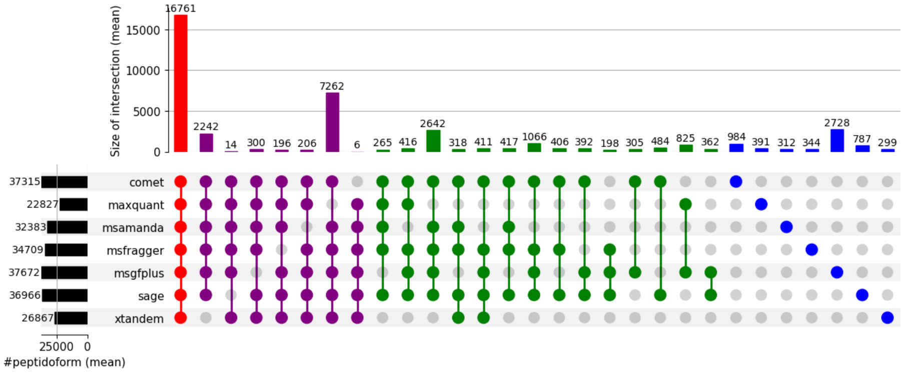

(b) Percolator

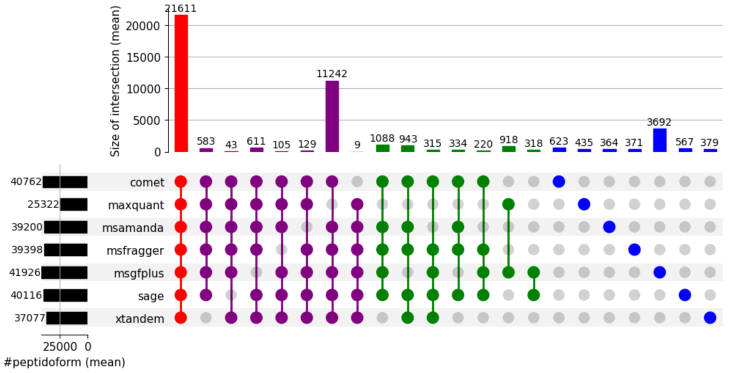

(c) MS2Rescore

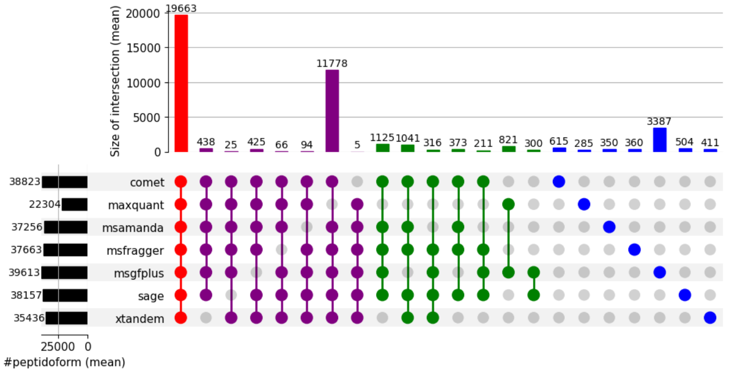

(d) Oktoberfest

Figure S49: Overlap of search engines per rescoring method, timsTOF Dataset, Swiss-Prot

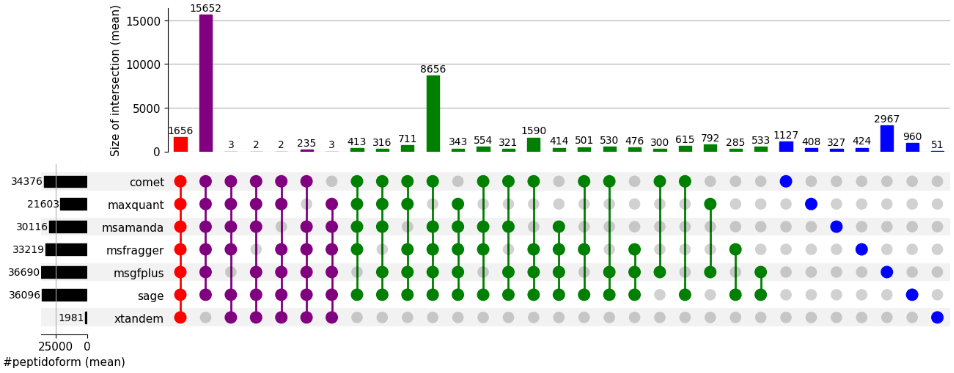

(a) raw search results

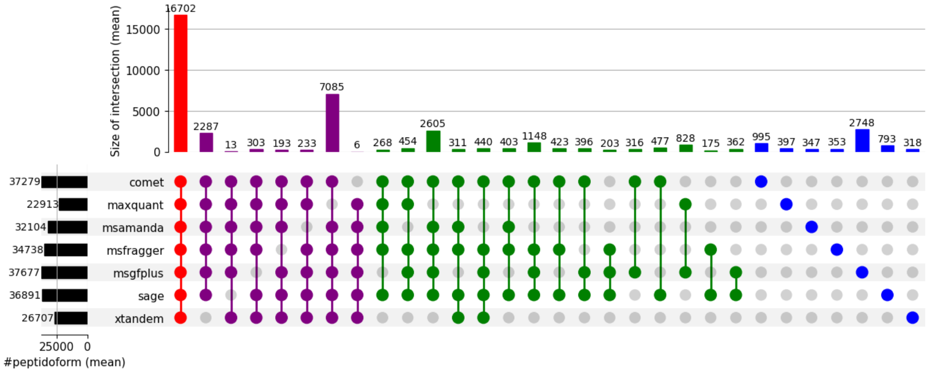

(b) Percolator

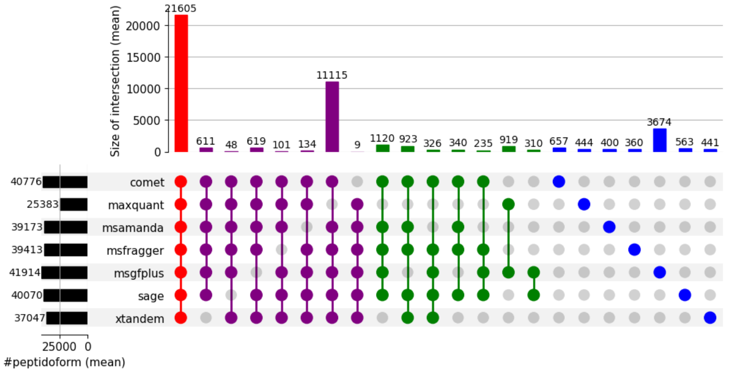

(c) MS2Rescore

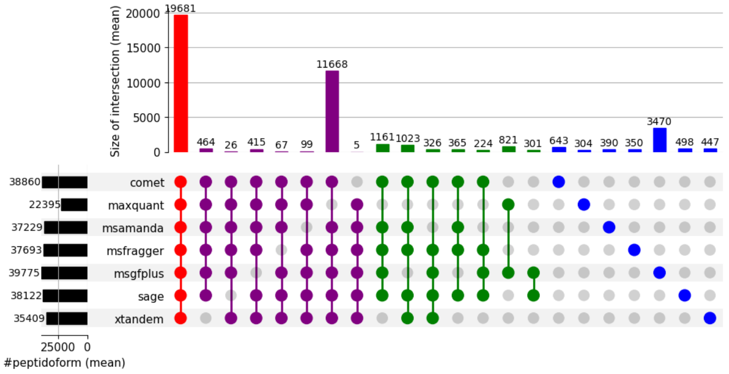

(d) Oktoberfest

Figure S50: Overlap of search engines per rescoring method, timsTOF dataset, Proteome

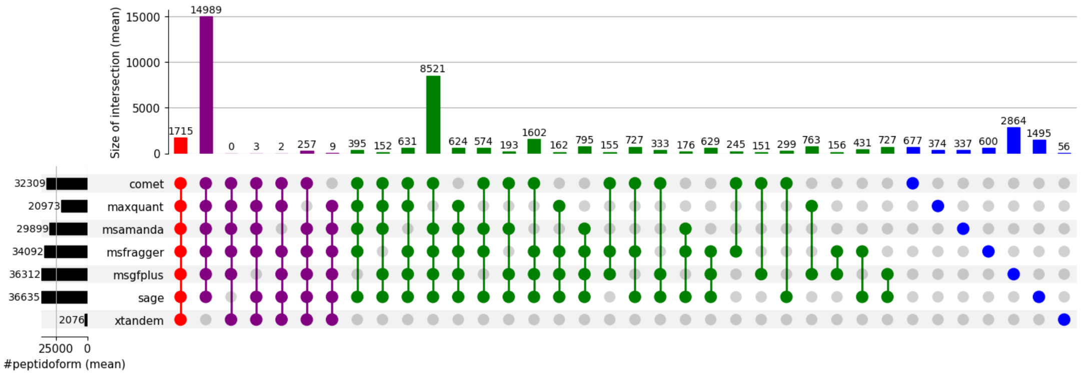

(a) raw search results

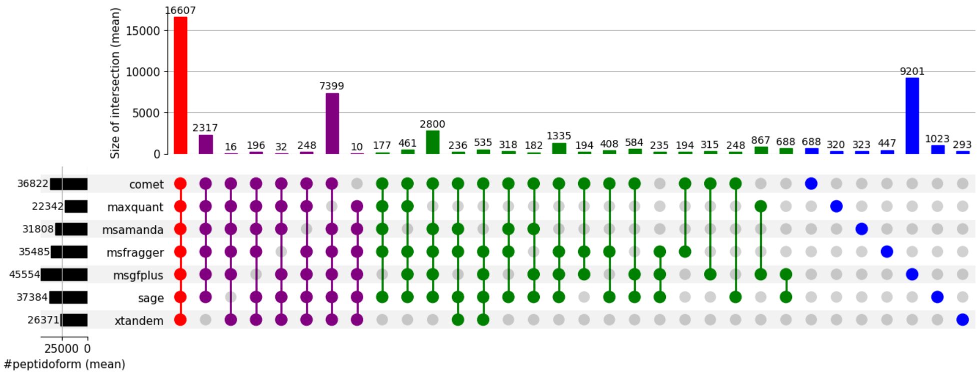

(b) Percolator

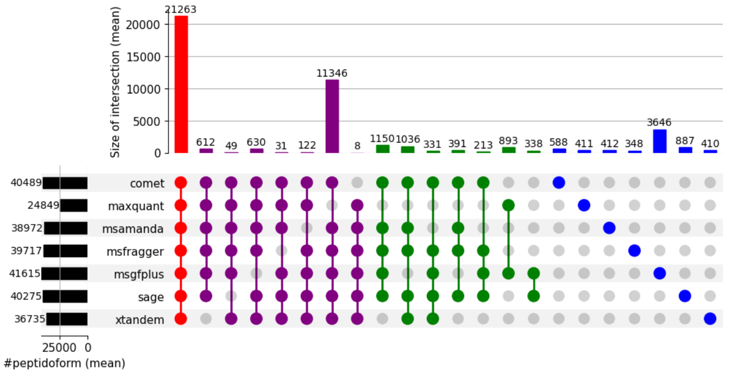

(c) MS2Rescore

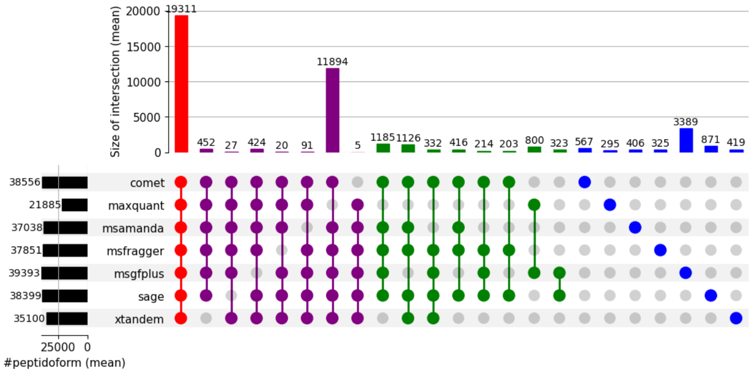

(d) Oktoberfest

Figure S51: Overlap of search engines per rescoring method, timsTOF dataset, ProHap

### 6 Entrapment Analyses

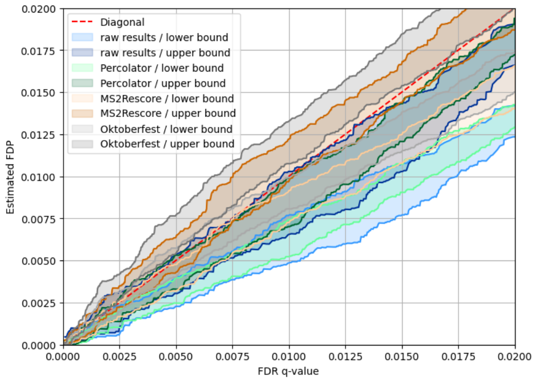

(a) Comet

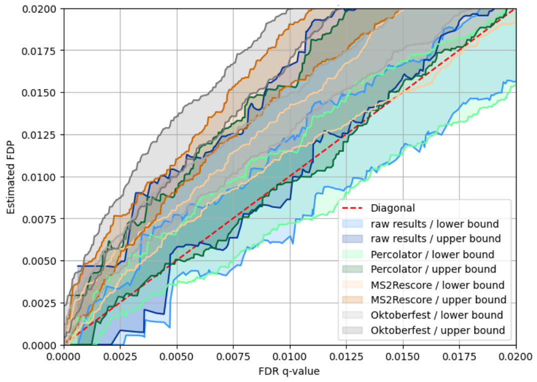

(b) MaxQuant

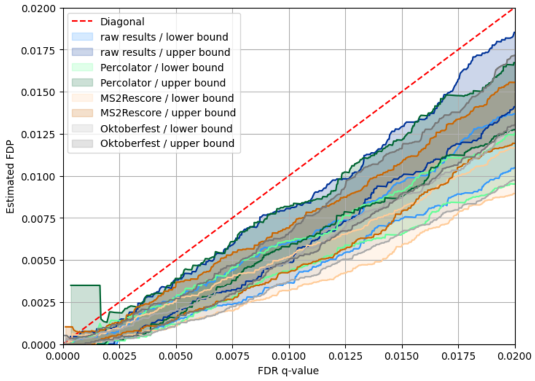

(c) MS Amanda

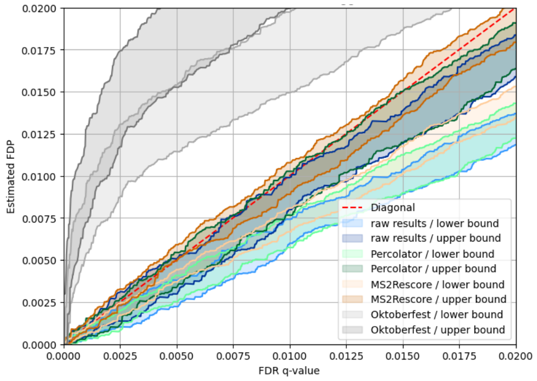

(d) MSFragger

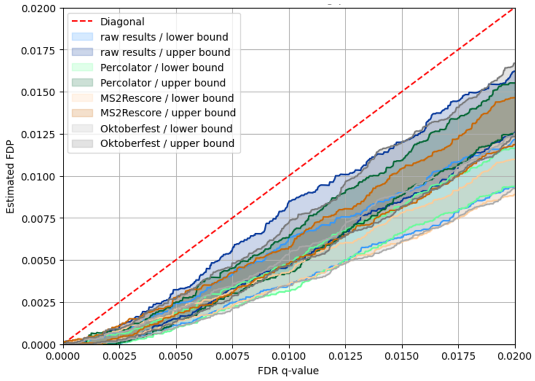

(e) MS-GF+

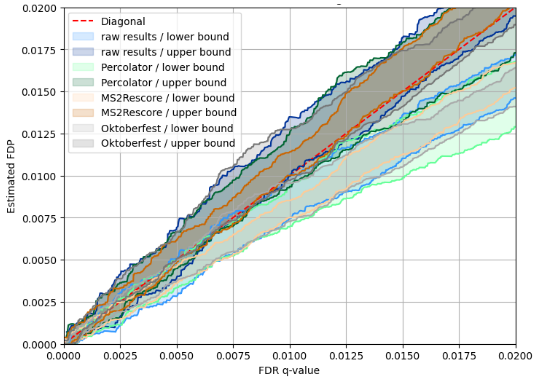

(f) Sage

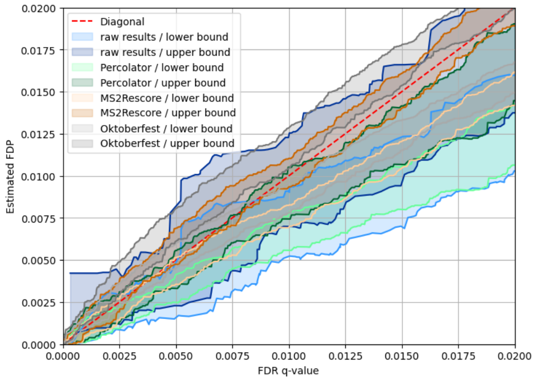

(g) X!Tandem

Figure S52: Entrapment analyses using the proteome database for the Cancer Array Dataset

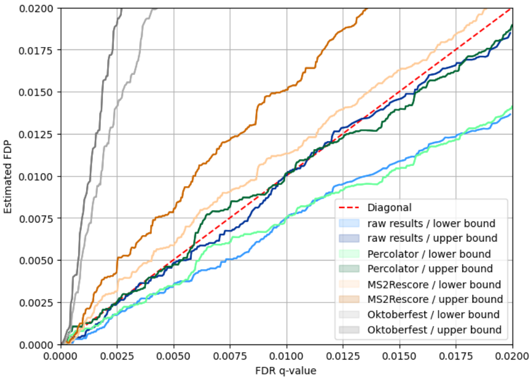

(a) Comet

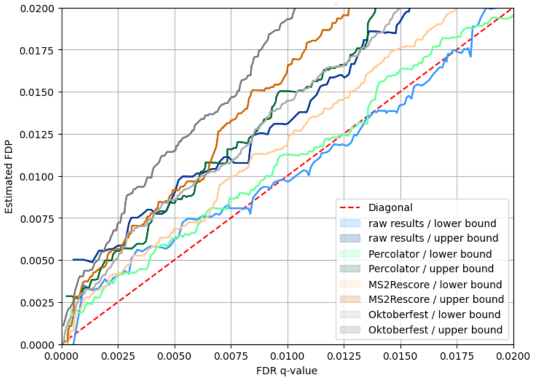

(b) MaxQuant

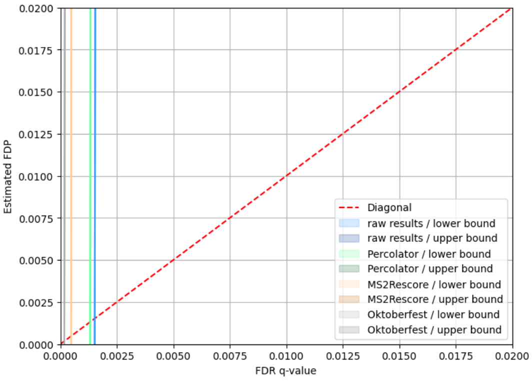

(c) MS Amanda

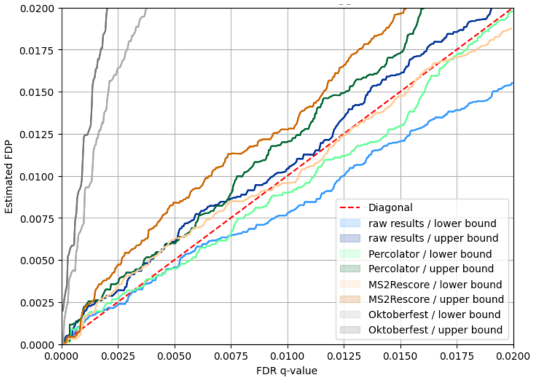

(d) MSFragger

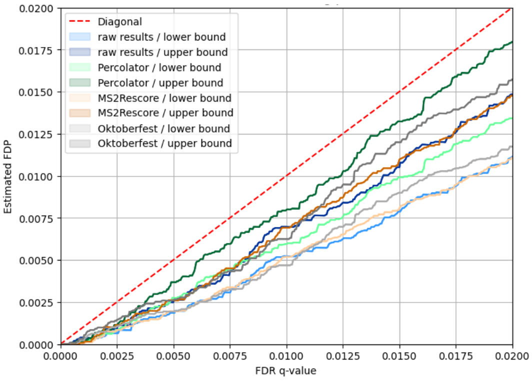

(e) MS-GF+

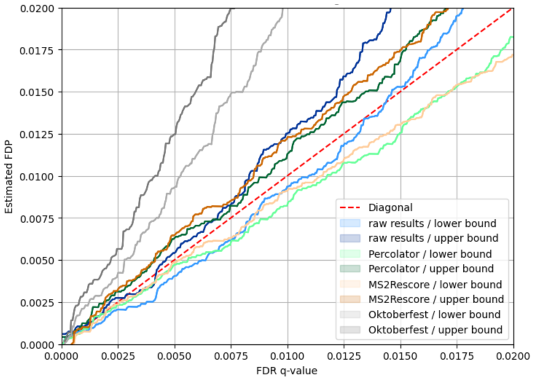

(f) Sage

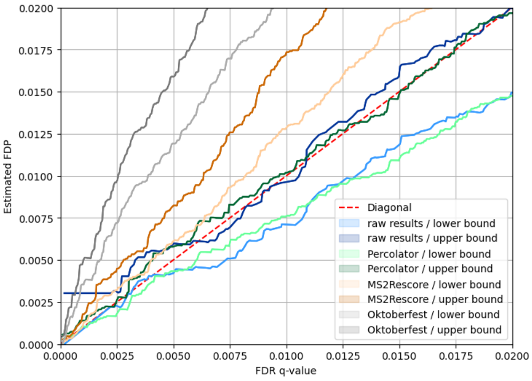

(g) X!Tandem

Figure S53: Entrapment analyses using the DB1 for the CAMPI Dataset

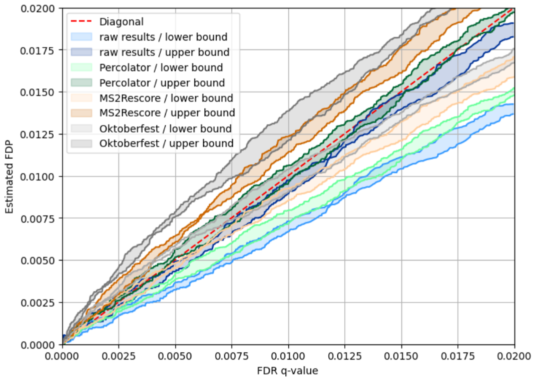

(a) Comet

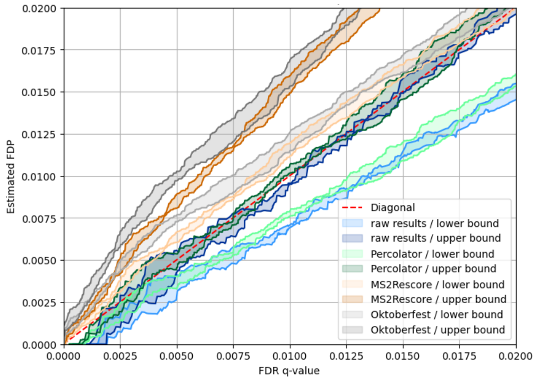

(b) MaxQuant

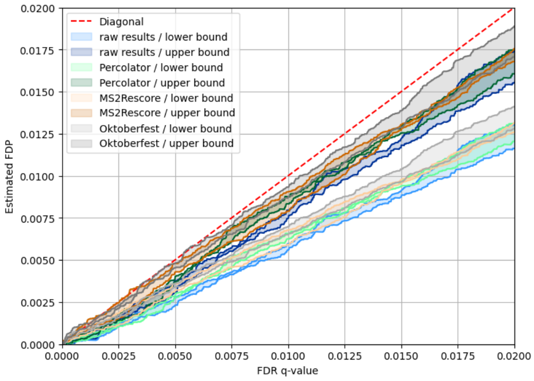

(c) MS Amanda

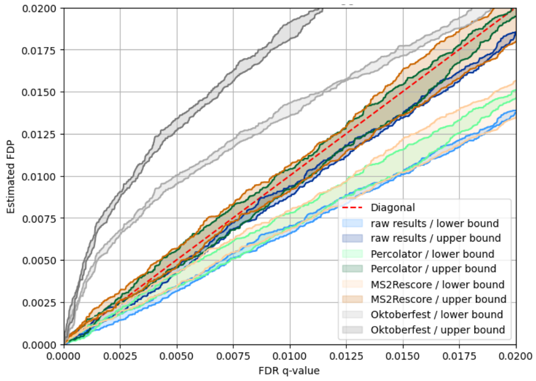

(d) MSFragger

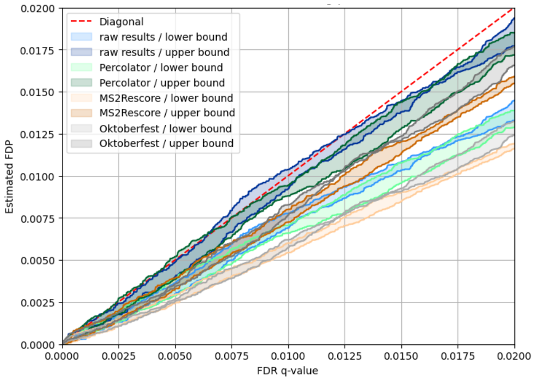

(e) MS-GF+

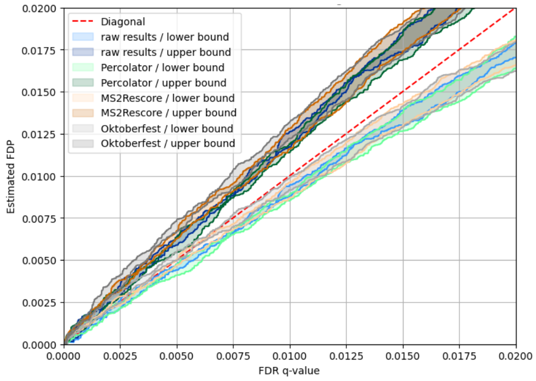

(f) Sage

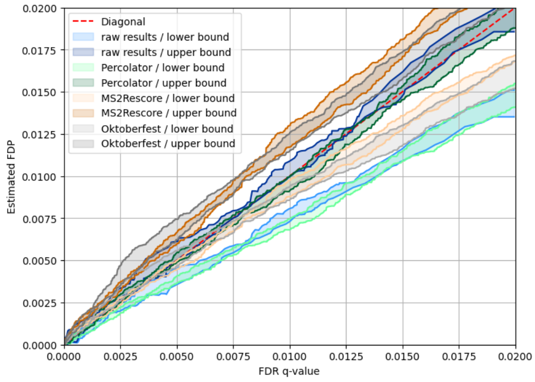

(g) X!Tandem

Figure S54: Entrapment analyses using the proteome database for the Orbitrap dataset

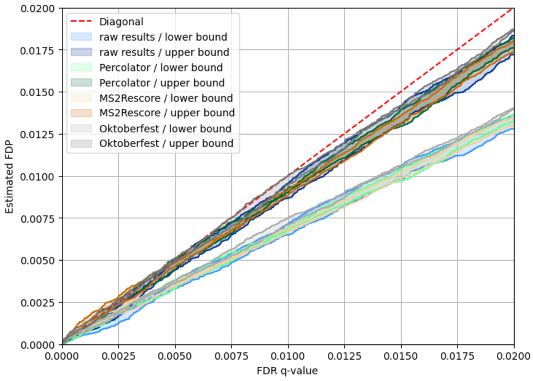

(a) Comet

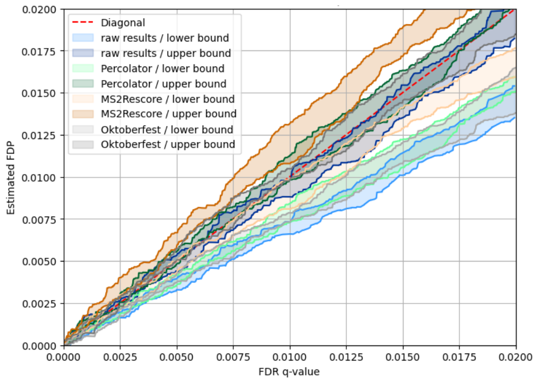

(b) MaxQuant

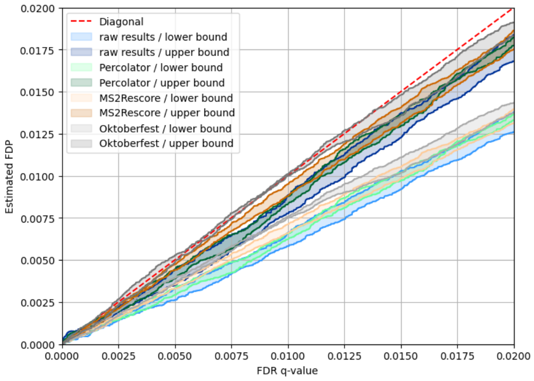

(c) MS Amanda

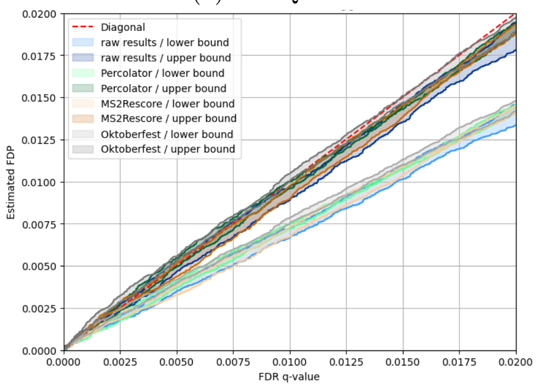

(d) MSFragger

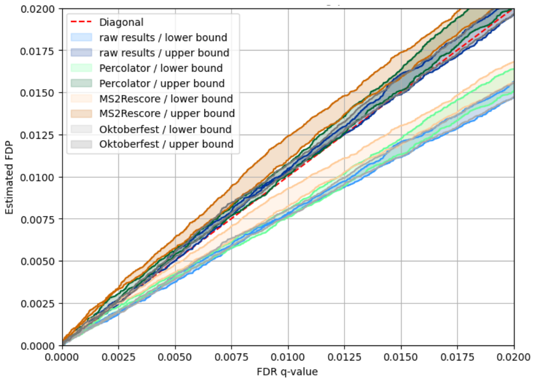

(e) MS-GF+

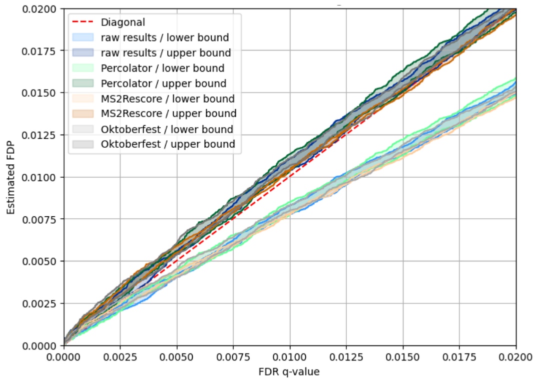

(f) Sage

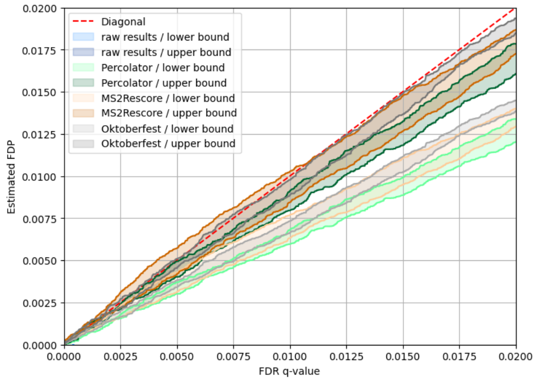

(g) X!Tandem

Figure S55: Entrapment analyses using the proteome database for the timsTOF dataset

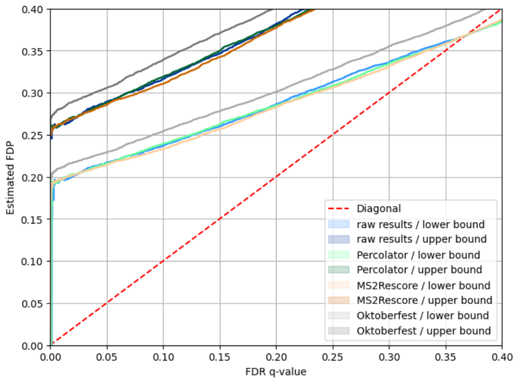

(a) MS Amanda, CAMPI Dataset, DB1

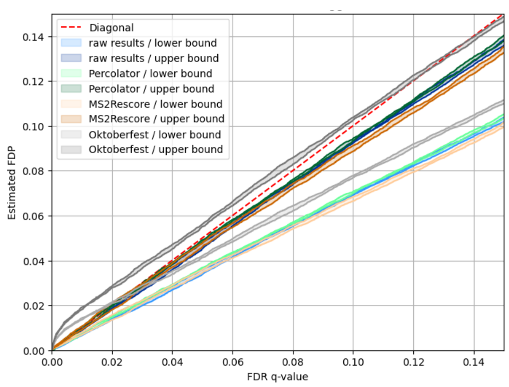

(b) MS Fragger, Orbitrap Dataset, proteome database

Figure S56: Entrapment analyses of selected search engines and datasets for higher ranges of estimated FDP and FDR values

### 7 Software tools

| Pipeline step | Software | Version | Container | Repository |
|---|---|---|---|---|
| all | mspepid | tag "rescore_comparison" | ghcr.io/medbioinf/mspepid:latest | https://github.com/medbioinf/mspepid |
| preprocessing | msconvert | 3.0.25073 | proteowizard/pwiz-skyline-i-agree-to-the-vendor-licenses:3.0.25073-842baef | https://github.com/ProteoWizard/container |
| preprocessing | tdf2mzml | 0.4 | quay.io/medbioinf/tdf2mzml:0.4 | https://github.com/medbioinf/muir |
| preprocessing | openms | 3.4.1 | quay.io/medbioinf/openms:3.4.1 | https://github.com/medbioinf/muir |
| preprocessing | FDR Bench | Commit #143f77 | quay.io/medbioinf/fdrbench-nightly:146f77 | https://github.com/medbioinf/muir |
| identification | Comet | v2024.01.0 | quay.io/medbioinf/comet-ms:v2024.01.0 | https://github.com/medbioinf/muir |
| identification | MaxQuant | 2.6.3.0 | quay.io/medbioinf/maxquant:2.6.3.0 | https://github.com/medbioinf/muir |
| identification | MS Amanda | 3.0.22.071 | quay.io/medbioinf/msamanda:3.0.22.071 | https://github.com/medbioinf/muir |
| identification | MSFragger | 4.2 | medbioinf/msfragger | https://github.com/medbioinf/mspepid |
| identification | MS-GF+ | v2024.03.26 | quay.io/medbioinf/msgfplus:v2024.03.26 | https://github.com/medbioinf/muir |
| identification | Sage | 0.15.0-beta.1 | quay.io/medbioinf/sage:v0.15.0-beta.1 | https://github.com/medbioinf/muir |
| identification | X! Tandem | 2017.2.1.4 | quay.io/medbioinf/xtandem:2017.2.1.4 | https://github.com/medbioinf/muir |
| postprocessing | Percolator | 3.08 | ghcr.io/percolator/percolator:branch-3-08 | https://github.com/percolator/percolator |
| postprocessing | Oktoberfest | 0.10.0 | quay.io/medbioinf/oktoberfest | https://github.com/medbioinf/muir |
| postprocessing | mzid-merger | 1.4.26 | quay.io/medbioinf/mzid-merger:1.4.26 | https://github.com/medbioinf/muir |

Table 1: Utilized software for mspepid, version, and container, as well as the container's origin repository.
